# Dual Nanoparticle-Driven Therapeutics for Leishmaniasis: A Mathematical Model of Targeted Macrophage and Parasite Elimination

**DOI:** 10.64898/2026.03.27.714640

**Authors:** Divya Arumugam, Mini Ghosh

## Abstract

**Background:** To control leishmaniasis, chemotherapy drugs are currently under development. However, these drugs often exhibit poor efficacy and are associated with toxicity, adverse effects, and drug resistance. At present, no specific drug is available for the treatment of leishmaniasis. Meanwhile, vaccine research is ongoing. Recent studies have analysed some experimental vaccines using mathematical models.

**Aim:** In previous work, drug targeting was focused on the entire human body rather than specifically addressing infected macrophages and parasites. In our current approach, we aim to eliminate infected macrophages and parasites through nano-drug design. Specifically, we utilise two types of nanoparticles: iron oxide and citric acid–coated iron oxide. Moving forward, we plan to advance this strategy using mathematical modelling of macrophage–parasite interactions.

**Methods:** We design PDE-based models of macrophages and parasites, incorporating cytokine dynamics, to support nano-drug development. Drug efficacy is estimated using posterior distributions to analyse phenotypic fluctuations of macrophages and parasites during the design phase. We investigate implicit and semi-implicit treatment schemes, focusing on energy decay properties. To model drug flow during treatment, we introduce a three-phase moving boundary problem. Comparative analyses are conducted to evaluate macrophage and parasite behaviour with and without treatment. Finally, the entire framework is implemented within a virtual lab environment.

**Results:** The results show that the nano-drug exhibits better efficacy compared to combined drug doses. We analysed and compared two types of nano-drug particles: iron oxide and citric acid–coated iron oxide. We discuss how the drug effectively targets and eliminates infected macrophages and parasites.

**Conclusion:** Our model’s results and simulations will support researchers conducting further studies in nano-drug design for leishmaniasis. These simulations are performed within a virtual lab environment.

## 1 Introduction

Leishmania are classified into several forms, with three main types of leishmaniasis commonly known as visceral leishmaniasis, cutaneous leishmaniasis, and mucocutaneous leishmaniasis. Leishmania parasites comprise more than 20 known species and over 30 distinct variants. These protozoan organisms are obligate intracellular parasites transmitted to humans through the bite of an infected female sand fly [1, 2]. During transmission, the flagellated form of the parasite—known as the promastigote—is injected into the host. Promastigotes are endocytosed by phagocytic cells, where they transform into amastigotes. These amastigotes mature and proliferate within host cells, eventually causing the infected cells to rupture. In infected cells, granulomas or localized inflammatory foci can form, leading to infection. Particularly in visceral leishmaniasis, granulomas develop in the liver or spleen [9, 10].

According to WHO, approximately 350 million people across all continents suffer from leishmaniasis worldwide. Overall, an estimated seven hundred thousand to 1 million new cases occur annually. An estimated fifty thousand to ninty thousand new cases of visceral leishmaniasis (VL) occur worldwide each year, but only 25–45% are reported to WHO [3, 4, 28, 6, 29, 8]. There is ongoing research on leishmaniasis, including statistical analysis, stochastic models, and mathematical models used to study population dynamics and develop control strategies. Here we share a few for your reference [81, 82, 83]. In biology, there are many in vivo studies on leishmaniasis, but only very few within-host models have been developed from the perspective of mathematical modeling. Specifically, the interaction between macrophages in humans and parasite dynamics after disease onset has been studied. Our main focus is particularly to develop a PDE model for the interaction between macrophages and parasites.

Macrophages play a vital role in leishmaniasis, helping to combat pathogens and contributing to anti-leishmaniasis responses [51, 12, 13]. Phagocytes are necessary components of the immune system, contributing to both pathogen defense and tissue repair mechanisms.Let us briefly introduce *M*_1_ and *M*_2_ macrophages, which are literally known as polarized forms of macrophages. *M*_1_ macrophages are referred to as pro-inflammatory macrophages, while *M*_2_ macrophages are closely associated with anti-inflammatory responses. *M*_1_ macrophages produce cytokines that promote inflammation and pathogen clearance, whereas *M*_2_ macrophages contribute to immunosuppression and tissue healing. The Macrophages *M*_1_ and *M*_2_ macrophages are different phenotypes, but they do not exist in isolation.which also demonstrate the plasticity and transitioning. These kind of transitions always influenced by the cytokine environment.However the activation of a dynamic spectrum allows macrophages based immunology which is balancing pathogen elimination with tissue repair. [27, 15, 16].

There are two important phases of Macrophages activation.*M*_1_ is typically dominating during the time of infection it acts as acute phase,while *M*_2_ acts as a chronic phase. This M1 and M2 polarization is largely virus dependent and its also shapes the immune response.During the time of viral infection *M*_0_ and *M*_1_ macrophages are upregulated. This kind of phenomenon has been observed particularly in viral disease like covid-19 and Rabies. The Macropahge through polarization helps to modulate the immune response.Although Receptors and ligands are important for this process with ligands such as *IL* − *R, IFN* − *γ, IL* − 4, and *IL* − 13 [17]serving as mediators. The different inflammatory phenotypes during different stage of infection can be adopted by the Macrophages.At all stages some degree of macrophages driven inflammation is involved.However the play between Macrophage phenotypes and some viral factors are the critical determinant of diseases [23].

*M*_1_ and *M*_2*a*_ macrophages are typically converge and primarily governed by a cytokine-dependent mechanism.Generally the human parasites and macrophages are generally undergo a dynamic switch toward *M*_2_ polarization. *M*_2_ macrophages can be classified into *M*_2*a*_, *M*_2*b*_, *M*_2*c*_ and *M*_2*d*_ subtypes [18, 19, 20, 21, 22].Some of the studies demonstrated the core protein of the hepatitis C virus which is typically inhibits polarization.Anhhow *M*_2*c*_ phenotype shift may happen in specific conditions. The researchers expanding the field in immunology for novel therapeutic interventions they studied the mechanism of plasticity of macrophage and pathogen interaction.

In earlier research, many studies were conducted using Leishmania major, an Old World species responsible for cutaneous leishmaniasis. However, Leishmania donovani is the causative agent of visceral leishmaniasis. The genomes of L. donovani and L. major are different, and these differences are routinely used to classify species and strains [24, 25, 26]. The predominant Leishmania Indian strain has been extensively employed in genetic, biochemical, and immunological studies, helping to identify key genetic determinants. Significant differences in chromosomal banding patterns have been detected both within and between species.

The entire work in this paper is arranged into seven sections. Section 1 introduces leishmaniasis and the granuloma model in the liver, along with a discussion of *M*_1_, *M*_2_, and macrophage subtypes, followed by an overview of the various strains responsible for the disease. Section 2 focuses on the mathematical modeling of leishmania-induced granuloma formation, incorporating dendritic cells, macrophages, parasites, and cytokine dynamics, both in the presence and absence of drug interaction. In this section, we developed a partial differential equation (PDE) model that captures the Macrophage and parasite interaction based on cellular interactions, ligands, and cytokines. Section 3 explains the boundary conditions of the granuloma model, this will helpful for simulations. Section 4 studies the velocity of u and boundary conditions of the developed model. In Section 5 we developed the three-phase moving boundary conditions which will helpful for drug designing. Section 6 describes the implicit and semi-implicit schemes but we formulated in polar coordinates,for ensuring the computational stability and to enhance the accuracy of the model. Section 7 presents the results and discussion, we focused particularly on the efficacy of nano drug is based on iron oxide and citric acid-coated iron oxide treatments across different strains of leishmaniasis.these findings helps to researchers to the potential and optimizing therapeutic strategies. Finally, the conclusion highlights the key insights which is drawn from the preceding sections and subsections of our finding and highlights directions for future research.

## 2 Mathematical Modelling of Leishmania Based on Granuloma Formation

In this study, we present a mathematical model of *Leishmania* infection based on granuloma formation. The model is formulated using partial differential equations (PDEs) that describe the interactions among macrophages, parasites, cytokines, and various immune cell types. We consider five stages of macrophages: *M*_1_, *M*_2_,*M*_2*a*_, *M*_2*b*_, *M*_2*c*_, and a repeated *M*_1_ stage to account for reactivation or persistence. Additionally, we define transitional macrophage states: 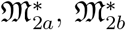, and 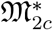, which represent intermediate stages between macrophage phenotypes. The parasite population is categorized into multiple stages: 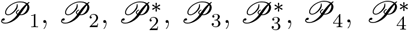, and *P*_5_, where the superscript ^∗^ denotes a transitional state from one stage to another. Typically, granulomas are composed of macrophages, dendritic cells, and T cells, which evolve dynamically within the granuloma and move with velocity *u*. In our model, we design the granuloma to include macrophages, dendritic cells, and various stages of T cells and B cells, all evolving with respect to time *T*. Furthermore, we assume that all species disperse or diffuse with appropriate diffusion coefficients. Somewhere within this framework lies the potential to simulate therapeutic interventions and immune responses. Anyhow, detailed descriptions of macrophage and dendritic cell dynamics are provided in the following sections.

### 2.1 Dendritic Cell Activation via Parasite Ingestion

This section provides a detailed explanation of how dendritic cells are stimulated through the ingestion of parasites. As initially described in reference [27], dendritic cells ingest parasites *P*_1_ and *P*_2_ as part of their antigen-presenting function. However, in our study, we extend this framework by considering five distinct types of parasites: *P*_1_, *P*_2_, *P*_3_, *P*_4_, and *P*_5_, each characterized by different densities and biological properties. Furthermore, we examine the following combinations of parasite interactions: (*P*_1_ + *P*_2_ + *P*_3_), (*P*_1_ + *P*_2_ + *P*_3_ + *P*_4_), and (*P*_1_ + *P*_2_ + *P*_3_ + *P*_4_ + *P*_5_), which represent increasingly complex infection scenarios. Let *N* denote the bursting number of parasites within a dendritic cell or macrophage, a parameter that plays a crucial role in determining the threshold for immune activation. In the context of our model, the mass of one macrophage is assumed to be equivalent to the combined mass of 150 parasites, providing a biologically relevant scaling factor for cellular interactions. Somewhere within this dynamic interplay of parasite ingestion and immune signaling lies the key to understanding the initiation of adaptive immunity. Anyhow, the activation of dendritic cells, denoted by *D*, is described in detail in the following section, where we explore the mathematical formulation and biological implications of this process.

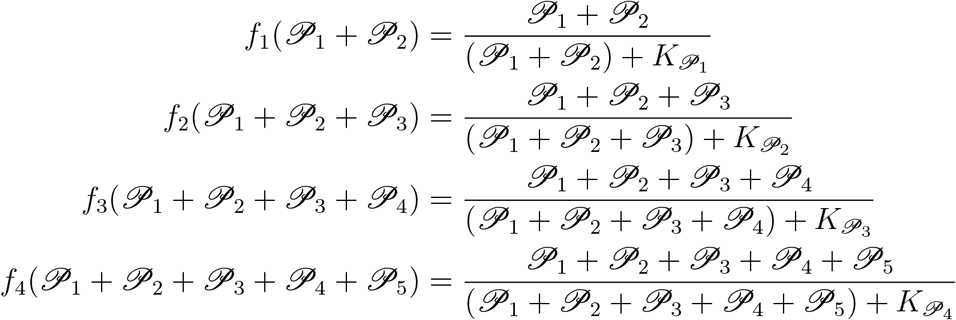

The average bursting pressure of the parasites is evaluated for the following combinations (*P*_1_ +*P*_2_),(*P*_1_ +*P*_2_ +*P*_3_),(*P*_1_ +*P*_2_ +*P*_3_ +*P*_4_),(*P*_1_ +*P*_2_ +*P*_3_ +*P*_4_ +*P*_5_). The Michaelis– Menten parameter is denoted by *K*_*P*_. In this study, we consider the specific parameters 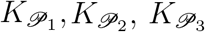, and 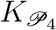. Additionally, the combinations of macrophages are defined as follows (*M*_1_ + *M*_2_),(*M*_1_ + *M*_2_ + *M*_2*a*_),(*M*_1_ + *M*_2_ + *M*_2*a*_ + *M*_2*b*_),(*M*_1_ + *M*_2_ + *M*_2*a*_ + *M*_2*b*_ + *M*_2*c*_).

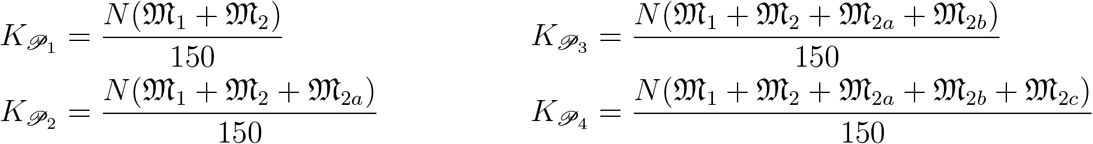

The following mathematical equation[1] describes the density of dendritic cells. The term *λ*_*D*_ *D*^0^ represents the rate of activation and is proportional to the density of inactive dendritic cells, denoted by *D*^0^. For modeling purposes, we assume that the number of inactive dendritic cells remains constant throughout the infection period. This equation captures both the activation of dendritic cells by parasites and their loss due to cell death.

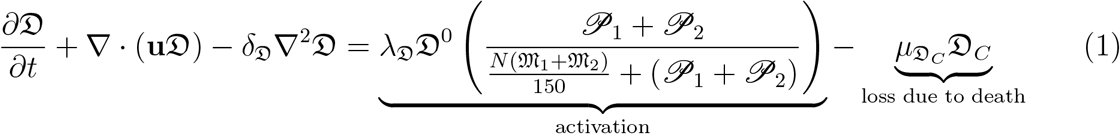

Figure 1 illustrates dendritic cell activation as a function of parasite identity. Each bar in the figure represents a unique combination of five different types of parasites (*P*_1_–*P*_5_), with the height of each bar indicating the total level of dendritic cell activation corresponding to that combination. The individual parasites—*P*_1_, *P*_2_, *P*_3_, *P*_4_, and *P*_5_—are color-coded for visual distinction. The first bar represents the combination (*P*_1_ + *P*_2_), while the second and third bars correspond to (*P*_1_ + *P*_2_ + *P*_3_) and (*P*_1_ + *P*_2_ + *P*_3_ + *P*_4_), respectively. Finally, the last bar—representing the full combination (*P*_1_ + *P*_2_ + *P*_3_ + *P*_4_ + *P*_5_)—shows the highest activation level.Furthermore, each type of parasite exhibits a distinct activation coefficient. Among them, *P*_4_ consistently enhances dendritic cell activation when included in combinations. *P*_5_, while contributing a smaller effect, shows a trend toward peak activation under certain conditions. Somewhere within these patterns lies insight into how parasite diversity influences immune response. Anyhow, the figure underscores the cumulative and differential impact of parasite identity on dendritic cell activation.

**Figure 1.**
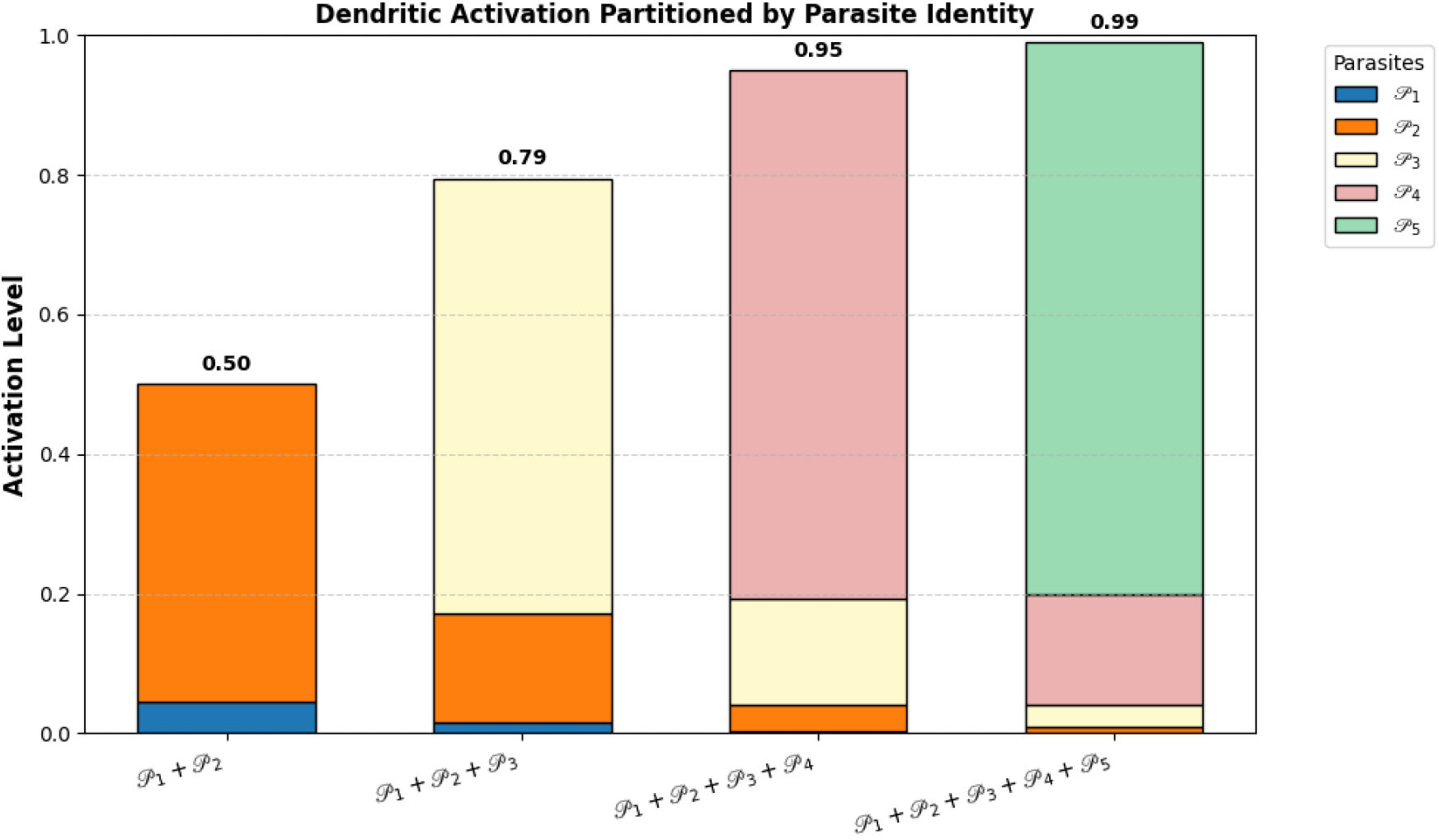
Dendritic activation Partitioned by parasite Identity.

Figure 2 illustrates dendritic cell dynamics at various sampled time points. We consider Equation [1], applied to different samples of ingested parasites as indicated in the corresponding dataset. Dendritic cell activation is analyzed in response to five distinct types of parasites and their associated macrophage states. The observed increase in cell density over time suggests progressive activation, recruitment, or proliferation of dendritic cells during infection or immune stimulation. The configuration *D*_1_ represents the combination of parasites (*P*_1_ + *P*_2_) and macrophages (*M*_1_ + *M*_2_), which exhibits a relatively lower density of dendritic cells. *D*_2_ includes (*P*_1_ + *P*_2_ + *P*_3_) and (*M*_1_ + *M*_2_ + *M*_2*a*_), showing moderate activation. *D*_3_ comprises parasites *P*_1_ through *P*_4_ and macrophages *M*_1_ through *M*_2*b*_, while *D*_4_ integrates all five parasite types (*P*_1_–*P*_5_) and all five macrophage states (*M*_1_–*M*_2*c*_).Furthermore, across these configurations, dendritic cell density peaks in *D*_3_ and *D*_4_ compared to *D*_1_ and *D*_2_, indicating a stronger activation effect at various sampled time points following parasite ingestion. Somewhere within these dynamic profiles lies a deeper understanding of how parasite diversity and macrophage phenotypes synergistically influence dendritic cell behavior. Anyhow, these findings underscore the importance of modeling immune cell interactions to capture the complexity of host-pathogen dynamics.

**Figure 2.**
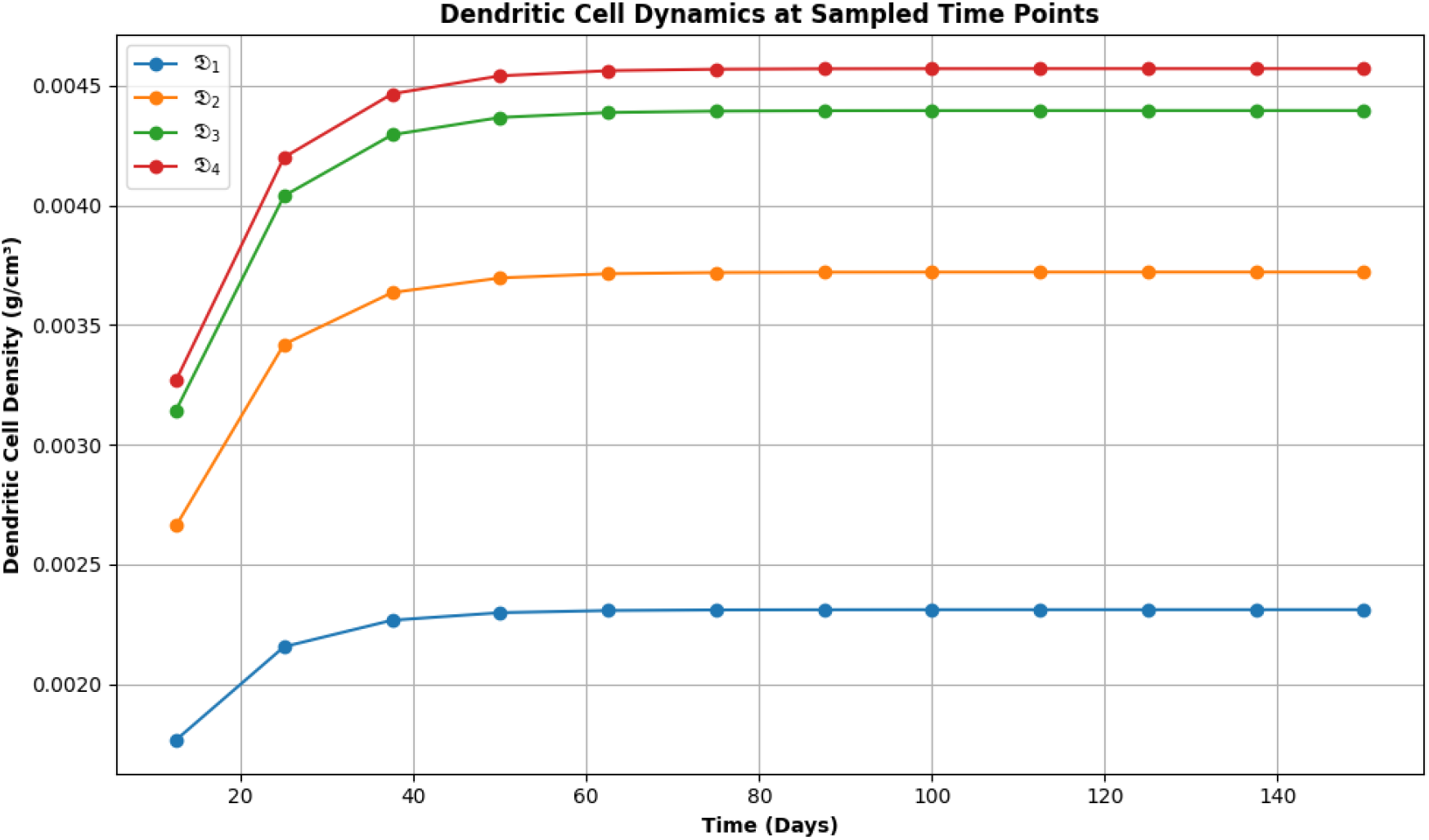
Dendritic cell activation dynamics at sampled time points.

### 2.2 Macrophages

In Equation (2), the right-hand side begins with a term describing the transition from *M*_2_ to *M*_1_, which is regulated by the cytokine *I*_*γ*_. The half-saturation constant for *I*_*γ*_ is denoted by *c*_*γ*_. The second term represents the loss associated with this transition. Furthermore, the activation of *M*_2_ is stimulated by the cytokine *I*_10_ and inhibited by *I*_*γ*_, with the halfsaturation constant for *I*_10_ denoted by *c*_10_. The third term accounts for the loss due to bursting, which follows Hill dynamics of the form:

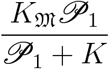

where 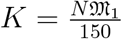, and *N* is the number of parasites inside the macrophages. Each macrophage is assumed to contain a mass equivalent to 150 parasites. The final term in Equation (2) represents the apoptosis of *M*_1_. Equation (3) is structurally similar to Equation (2), but includes a third term representing the loss due to bursting in parasite *P*_2_, and a fourth term representing apoptosis in macrophage *M*_2_. In Equation (4), the first term describes the recruitment of *M*_2_ macrophages, regulated by the cytokine IL-4. The half-saturation constant for IL-4 is denoted by *k*_IL4_.Somewhere within these equations lies the mechanistic insight into how cytokine signaling orchestrates macrophage transitions and immune regulation.Here, the following sections provide further elaboration on the mathematical structure and biological implications of these terms [28, 29, 30, 31, 32, 33, 34, 35, 36, 37].

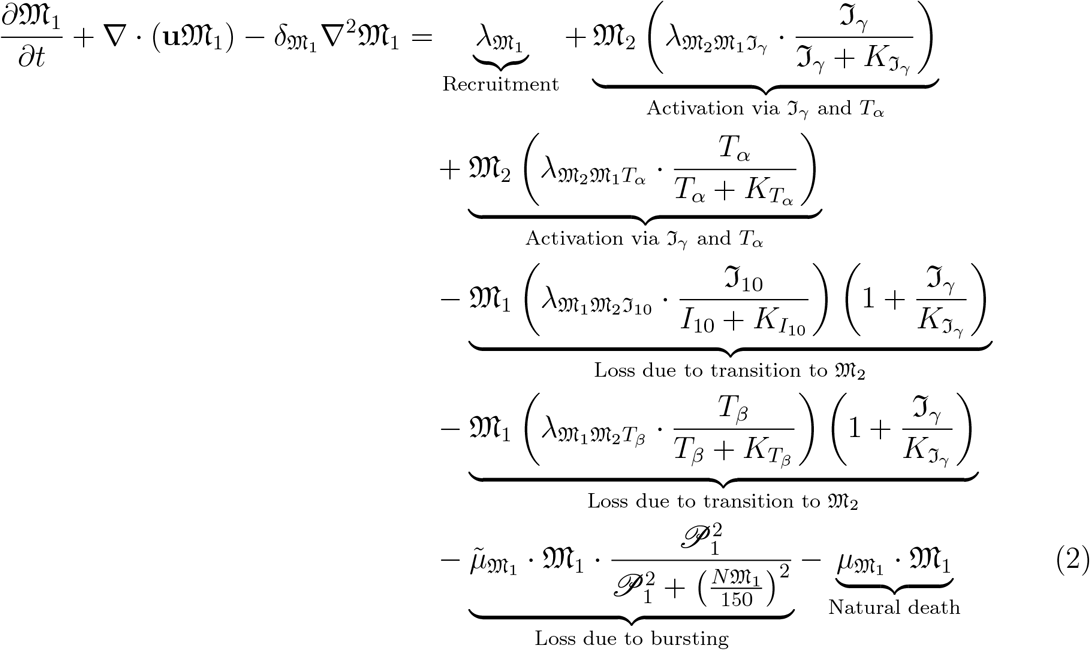

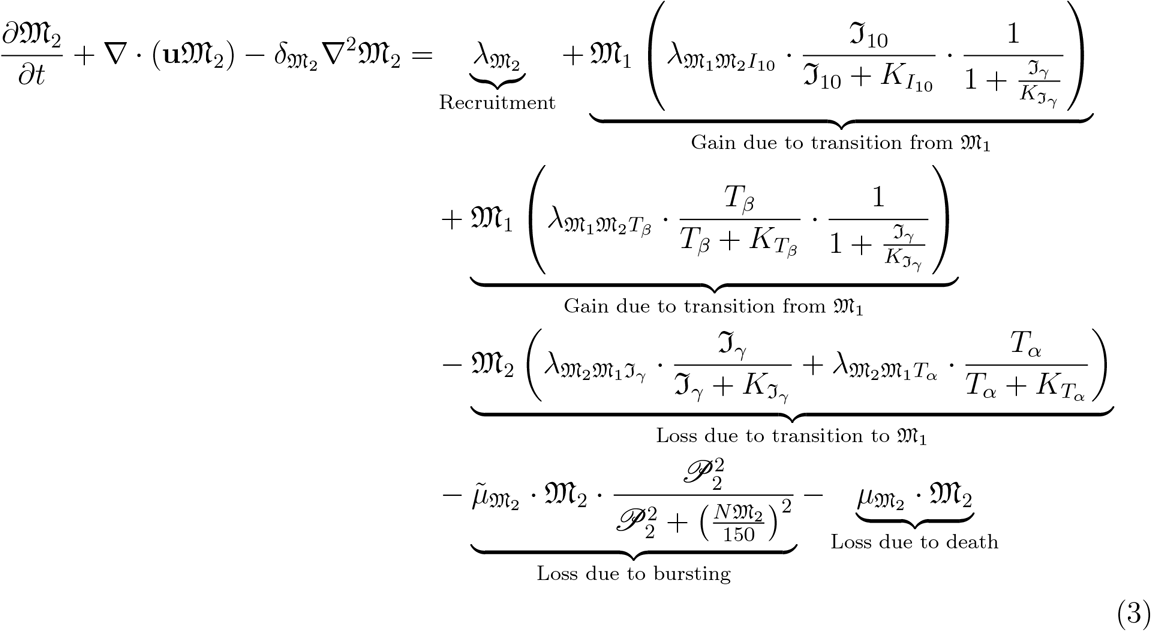

In our work, we assume that 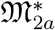 represents a transitional state between *M*_2_ and fully differentiated *M*_2*a*_ macrophages,emerging during the intermediate phase of the anti-inflammatory response and wound healing. This transition is primarily regulated by the cytokines ℑ*L*-4 and ℑ*L*-13. The second phase involves the acquisition of 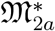 macrophages via transition from *M*_2_, mediated by the cytokine ℑL-10. The third phase accounts for the loss of 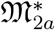 macrophages due to bursting, influenced by ℑFN-*α* and the parasite population 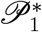, which will be detailed in subsequent sections. The fourth phase describes the natural death of 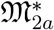 macrophages. The fifth phase involves tissue damage through fibrosis. The final phase represents tissue repair facilitated by parasite interaction, modulated by T-cell activity (T-modulated). In Equation (5), the remaining terms are consistent with those in Equation (4), except for: The third phase, which now depends on the parasite population 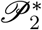 instead of 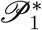. The final phase, which explicitly models tissue repair via parasite interaction, modulated by T-cell activity.For parameters we assume subparts of M2 as 60:20:20 for *M*_2*a*_,*M*_2*b*_,*M*_2*c*_.

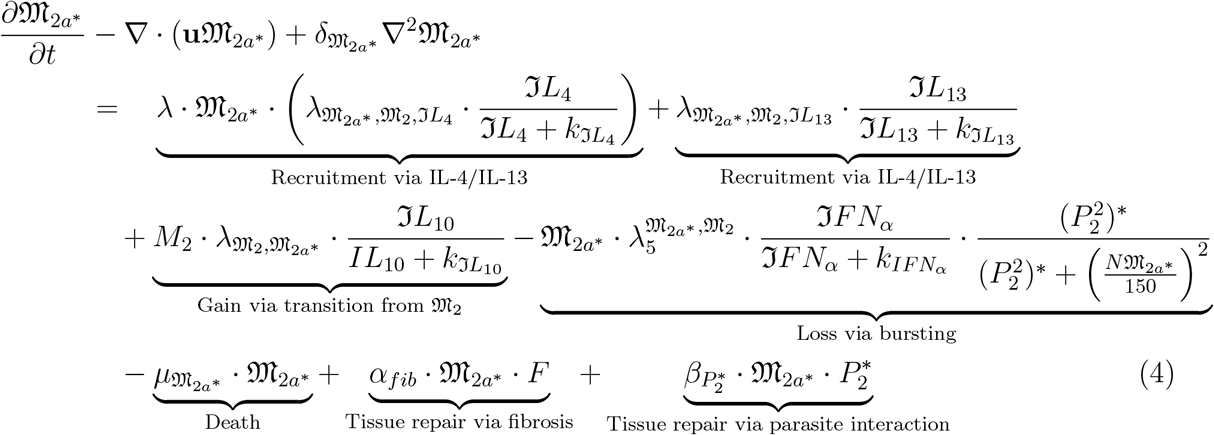

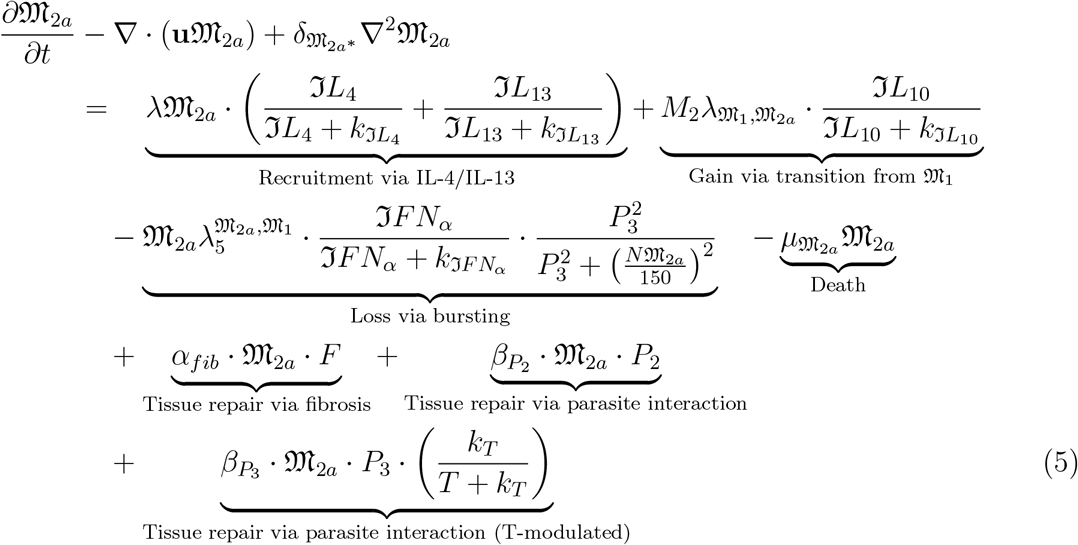

The equation for 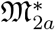 describes the transition from *M*_2*a*_ to *M*_2*b*_ macrophages, mediated by the cytokines IC and TLR. Additionally, 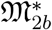 is acquired via transition from *M*_2_, regulated by the cytokine IL-10. The third phase represents the loss of 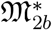 macrophages due to bursting, influenced by IFN-*α* and the parasite population (*P* 3)^∗^, as detailed in the following section. The fourth phase accounts for the death rate of 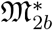 macrophages. The fifth phase involves tissue repair through interaction between the parasite population *P* 3^∗^ and the cytokine IL-10. The final phase models tissue repair via TNF–parasite interaction. The equation for *M*_2*b*_ is similar to that of 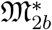, except that the last four phases are based on the parasite population *P* 4^∗^. Tissue repair in this context is regulated by the cytokines IL-10 and TNF.

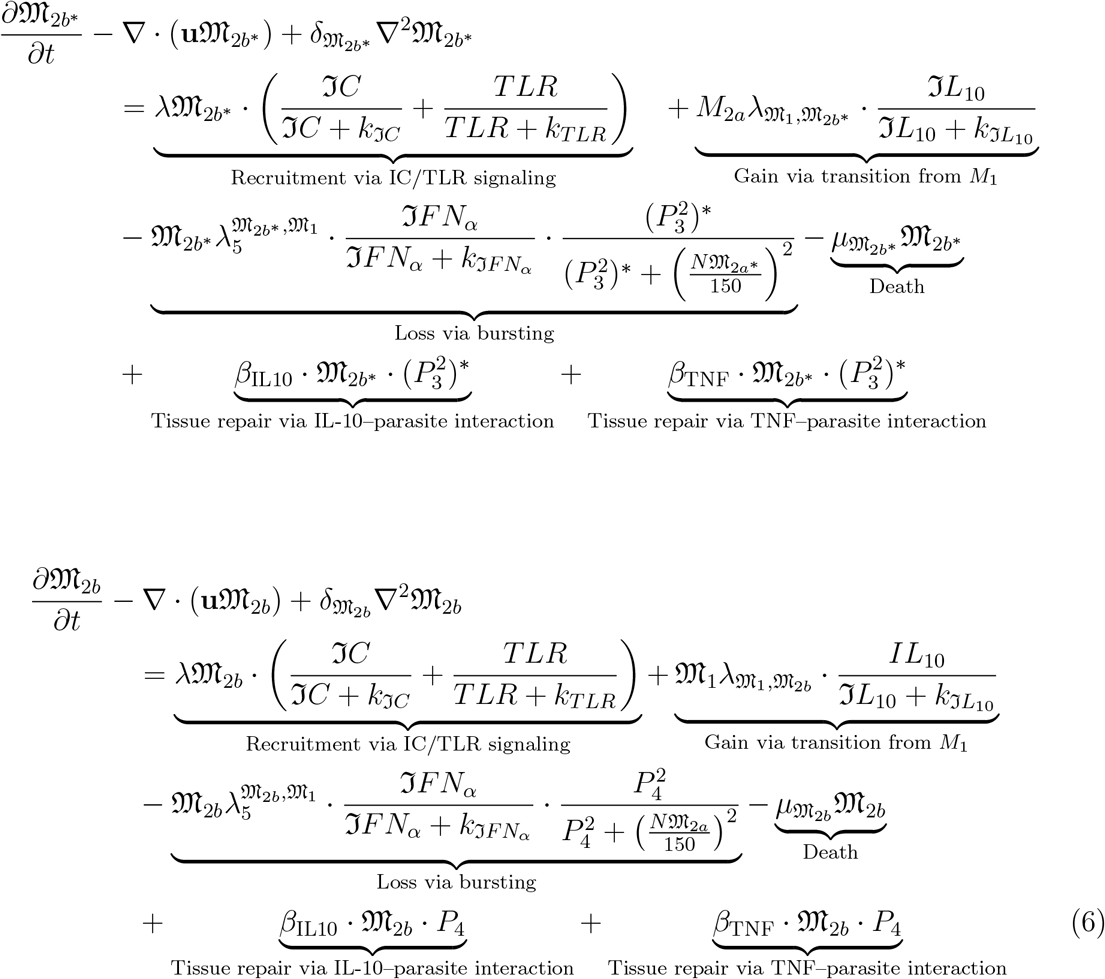

The equation for 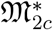 describes the transition phase from *M*_2*b*_ to *M*_2*c*_ macrophages, mediated by the cytokines ℑ*L*-10 and TGF-*β*. This phase also includes the gain of 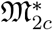 via transition from *M*_2*b*_. The loss term is governed by the cytokine ℑ*FN*-*α* in conjunction with the parasite population 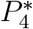. Tissue repair is modeled through the interaction of ℑ*L*-10 and TGF-*β* cytokines with the parasite population 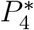. The final term in this equation corresponds to the parasite population 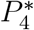. In the final macrophage equations, the last three terms are associated with the parasite population *P*_5_.

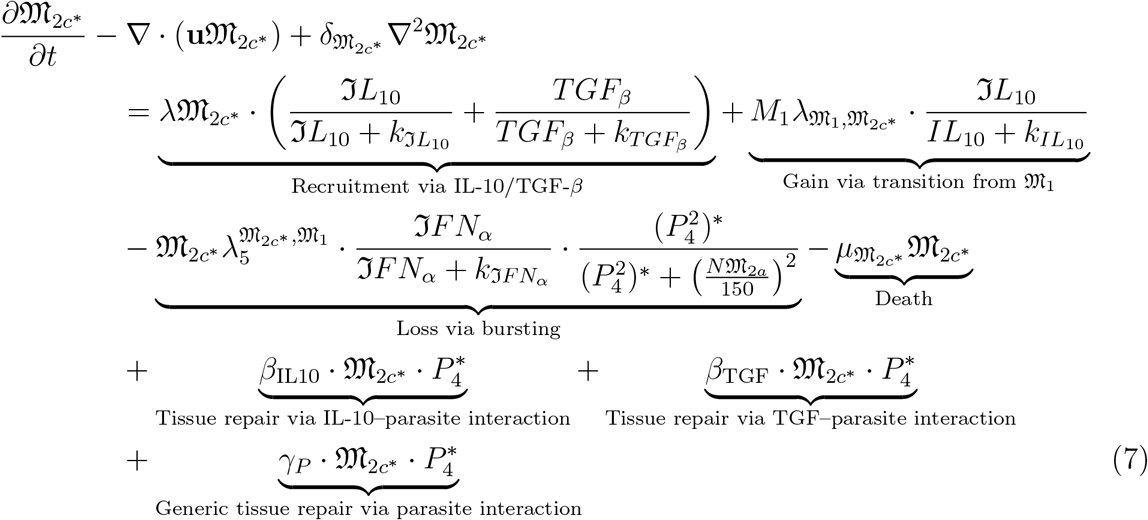

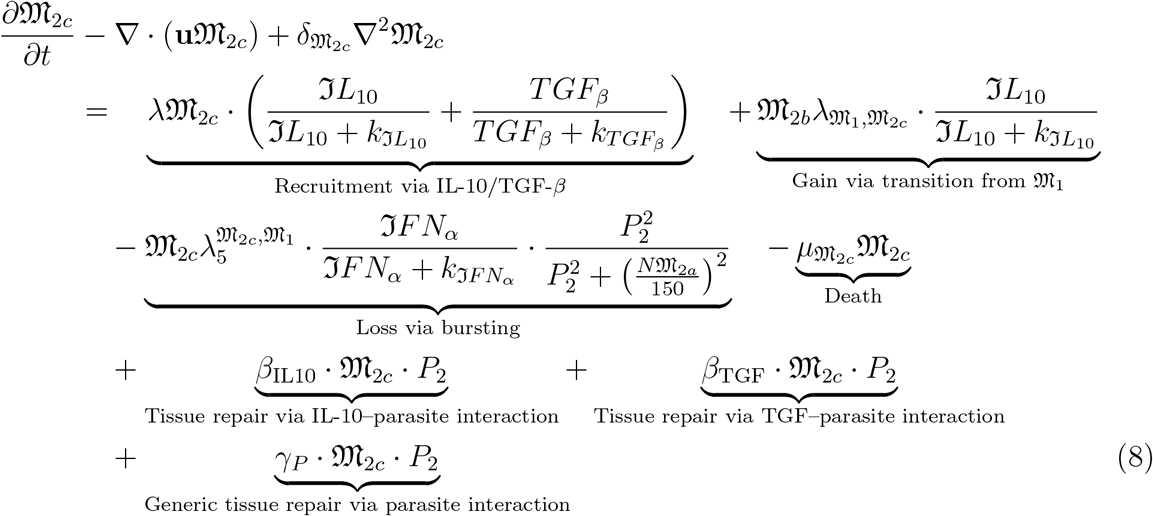

### 2.3 Modeling parasites based on strains

Parasites play a vital role in the progression of leishmaniasis; however, different parasite strains cause distinct forms of the disease. In our work, we focus on modeling the leishmania parasite dynamics, incorporating both classical leishmaniasis and post-kala-azar dermal leishmaniasis (PKDL). Based on this framework, we define specific strains such as *L. donovani, L. mexicana*, and *L. major*.Our study particularly concentrates on granuloma formation in the liver. We assume the granuloma has a diameter of 300 *µ*m, from which we derive the radius and fix the center at 0.5. The initial parasite growth and death rates are strain-dependent. Accordingly, we model drug diffusion using iron oxide-based nanodrugs. In our setup, iron oxide is typically coated with PEI-25, citric acid nanocomposites, with each coating tailored to the specific parasite strain. The growth rate of each strain is influenced by these drug formulations. The following figure illustrates the spatial-temporal model for the *L. donovani* and *L. mexicana* parasite strains.Our assumptions are based on the references [84, 85].

Figure 3 illustrates the spatial and temporal profile of granuloma formation, specifically in the liver. In this study, the granuloma diameter is fixed at 300. The figure presents results from testing two different parasite strains using a single nano-drug: citric acid-coated iron oxide. The temporal profile of parasite dynamics is shown for two strains—Leishmania donovani and Leishmania mexicana—which correspond to visceral leishmaniasis and cutaneous leishmaniasis, respectively. Initial growth and death rates of the parasites, based on strain-specific characteristics, are detailed in Table 1. These parameters were used to simulate the results in Figure 3, employing two numerical solvers: LSODA and RK45.Subplots A–E in Figure 3 represent simulations using L. mexicana with citric acid-coated iron oxide, as referenced in [citation]. In these plots, the temporal mean of parasite concentration reaches a value of 1 in some cases, while in others it approaches but does not fully reach this value.Subplots F–J depict simulations using the same nano-drug but with a different strain, L. donovani. Among these, only subplot I reaches the mean value of 1, while sub-plots F, G, H, and J approach this value but do not attain it completely.Notably, subplots A and B, when compared with F and G, show distinct differences in observed data values. The remaining subplots—C, D, E, H, I, and J—exhibit slight variations in observed values, suggesting nuanced strain-specific responses to the nano-drug treatment.

**Table 1:** Strain-specific parameters for Drug (*D*), growth rate (*λ*), and death rate (*µ*).

| Strain | Drug | $\lambda$ | $\mu$ | Drug | $\lambda$ | $\mu$ | Drug | $\lambda$ | $\mu$ |
| --- | --- | --- | --- | --- | --- | --- | --- | --- | --- |
| – | (CCIO) | (*L.D*) | (*L.D*) | (CCIO) | (*L.M*) | (*L.M*) | (IO) | (*L.D*) | (*L.D*) |
| P1 | $66 \times 10^{-9}$ | 1.00 | 0.10 | $66 \times 10^{-9}$ | 1.000 | 0.010 | $45 \times 10^{-9}$ | 1.00 | 0.10 |
| P2 | $66 \times 10^{-9}$ | 1.00 | 0.12 | $66 \times 10^{-9}$ | 1.000 | 0.010 | $45 \times 10^{-9}$ | 1.00 | 0.10 |
| P2* | $66 \times 10^{-9}$ | 0.80 | 0.11 | $66 \times 10^{-9}$ | 0.058 | 0.010 | $45 \times 10^{-9}$ | 0.80 | 0.11 |
| P3 | $66 \times 10^{-9}$ | 0.85 | 0.08 | $66 \times 10^{-9}$ | 0.058 | 0.010 | $45 \times 10^{-9}$ | 0.85 | 0.08 |
| P3* | $66 \times 10^{-9}$ | 0.85 | 0.09 | $66 \times 10^{-9}$ | 0.058 | 0.010 | $45 \times 10^{-9}$ | 0.85 | 0.09 |
| P4 | $66 \times 10^{-9}$ | 0.89 | 0.10 | $66 \times 10^{-9}$ | 0.058 | 0.010 | $45 \times 10^{-9}$ | 0.89 | 0.10 |
| P4* | $66 \times 10^{-9}$ | 0.89 | 0.10 | $66 \times 10^{-9}$ | 0.058 | 0.010 | $45 \times 10^{-9}$ | 0.89 | 0.10 |
| P5 | $66 \times 10^{-9}$ | 0.90 | 0.10 | $66 \times 10^{-9}$ | 0.058 | 0.010 | $45 \times 10^{-9}$ | 0.90 | 0.10 |

**Table 2:**
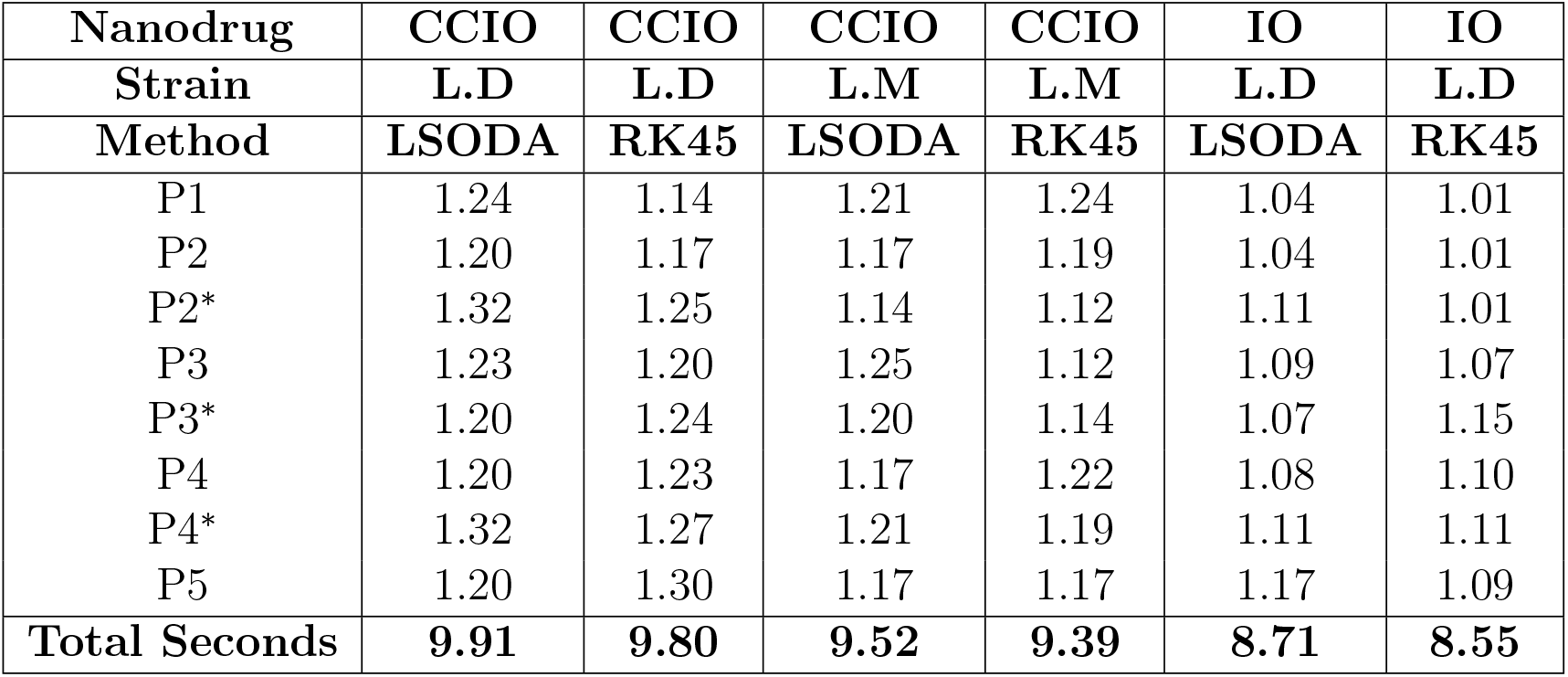
Performance Comparison of LSODA and RK45 Methods for Nanodrug Strains.

**Figure 3.**
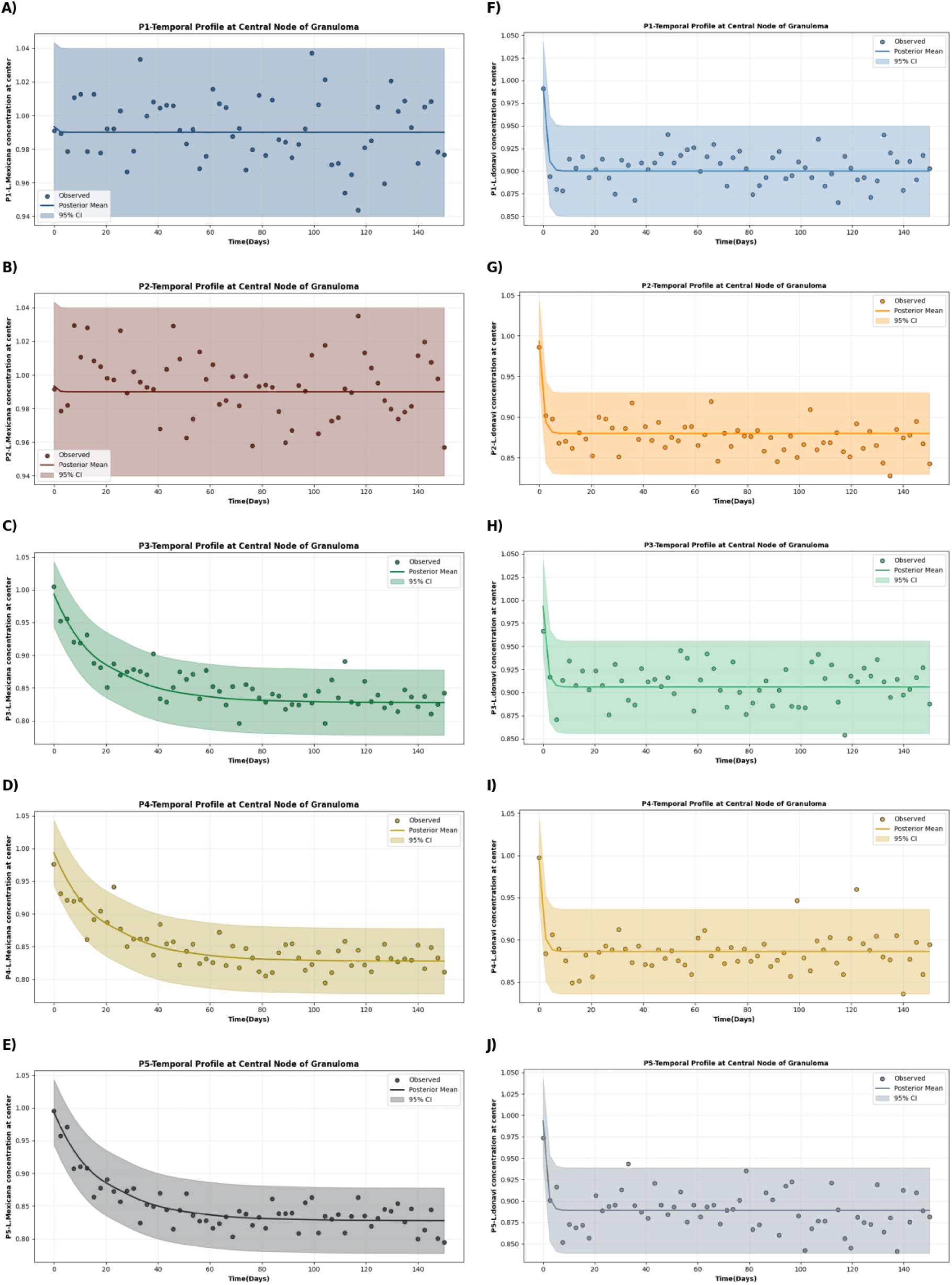
Dendritic activation Partitioned by parasite Identity.

### 2.4 Mathematical Equations describes the density of Parasites changes over time

Leishmania is an infectious disease in which parasite density plays a vital role in capturing the progression from the initial stage of infection to more advanced stages. This section explains the density of parasites across different stages—denoted as *P*_1_, *P*_2_, *P*_3_, *P*_4_, and *P*_5_.Parasite *P*_1_ typically survives in a pro-inflammatory environment, residing within macrophages of type *M*_1_. The equation governing *P*_1_ satisfies the Leishmania density condition and describes the growth dynamics of *M*_1_, capturing the logistic proliferation of Leishmania within the macrophage–parasite interaction [28, 29, 30, 31, 32, 33, 34, 35, 36, 37]. In contrast, parasite *P*_2_ survives in anti-inflammatory macrophages, specifically within *M*_2_ cells. A detailed description of parasites *P*_1_ and *P*_2_ is provided in [reference]. We assume the transition stage between *P*_2_ and *P*_3_ is denoted as 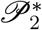, the transition from *P*_3_ to *P*_4_ as 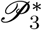, and the transition from *P*_4_ to *P*_5_ as 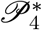.

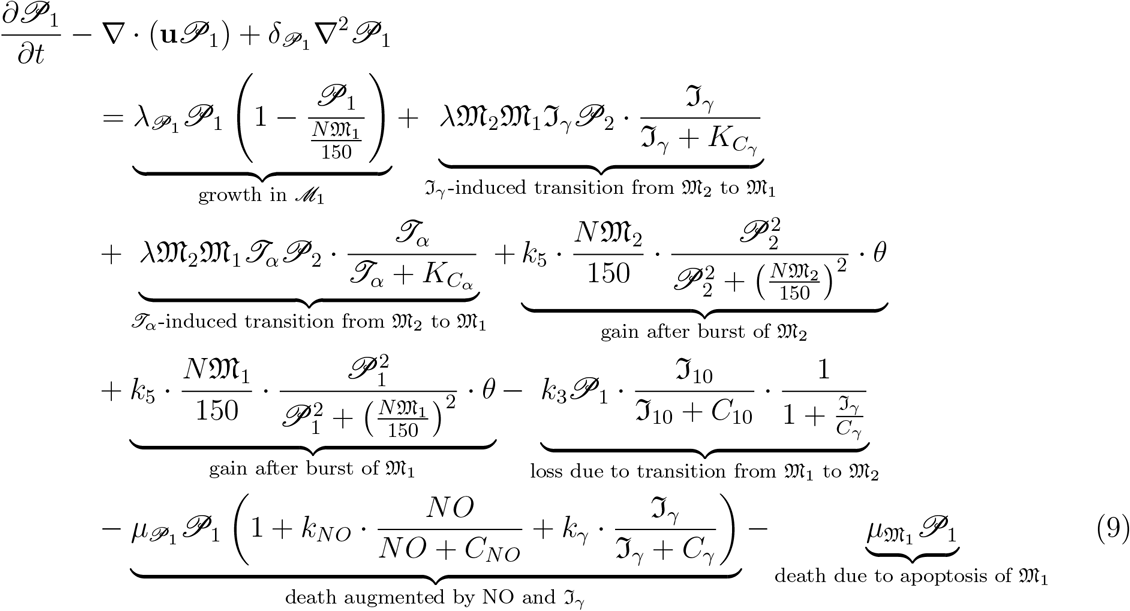

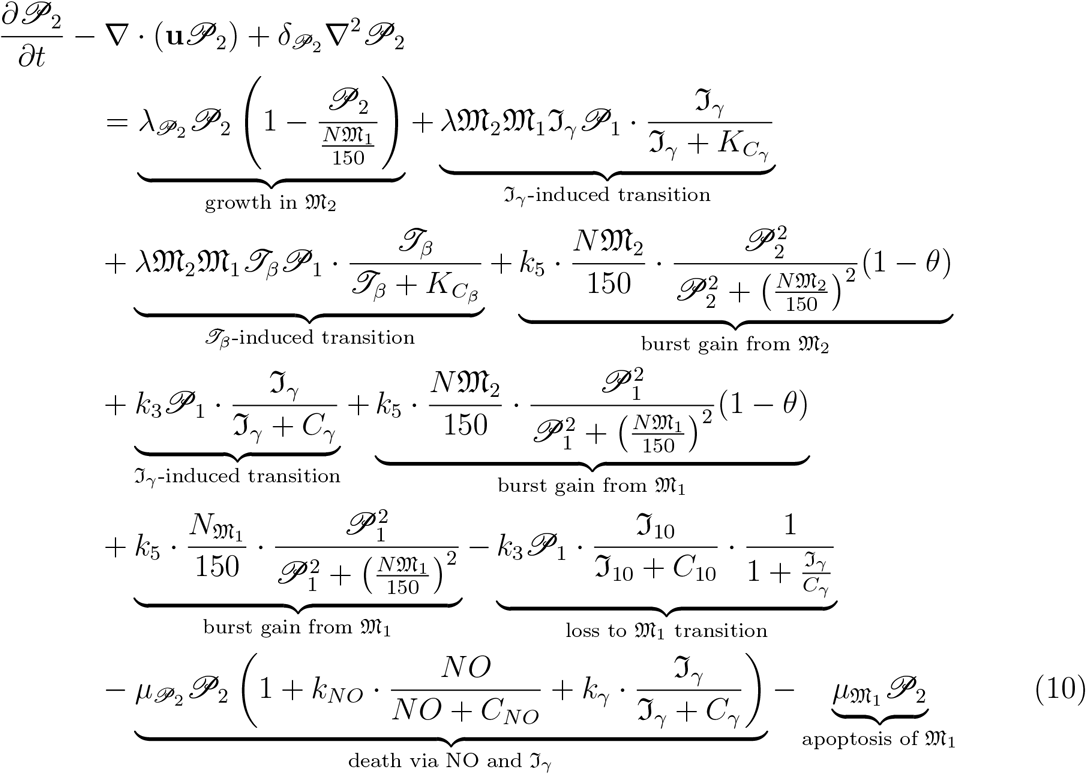

Let us begin with the new parasite density, denoted as 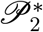, which differs significantly from *P*1 and *P*2. The parasite *P*2 may transition into *P*3 through various cytokinesis mechanisms. However, it produces either a very low amount or no nitric oxide at all. We consider the intermediate stage of this transition as 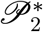. This parasite resides within *M*_2*a*_ macrophages, whose detailed description is provided in the previous section. During the transition, the density of 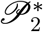contributes to the development of Leishmania rather than its decay. The left-hand side of the equation representing 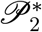 describes its growth within *M*_2*a*_, modeled using a logistic growth form. Here, N denotes the carrying capacity of 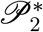 in *M*. If 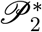 exceeds 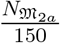, growth ceases—similar to the behavior observed in *P*_1_ and *P*_2_. The cytokine *I*_10_ induces the transition, and when *M*_2*a*_ bursts, it releases 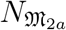 parasites. The parameter *θ* represents the ingested parasite *P*_2_. An alarmin-induced drift facilitates the transition from 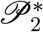 to *P*_3_, mediated by the cytokine ℑ*L* − 33. During this transition, *P*_2_ experiences loss due to cytokines such as ℑ_10_, *J NFβ*, and ℑ*L* − 33. Occasionally, cell death occurs via nitric oxide (NO) and *Iγ*. Importantly, *M*_2*a*_ macrophages cannot replicate themselves like *M*_2_ macrophages; they produce very little or no progeny. Apoptosis of *M*_2*a*_ may also occur during this process.

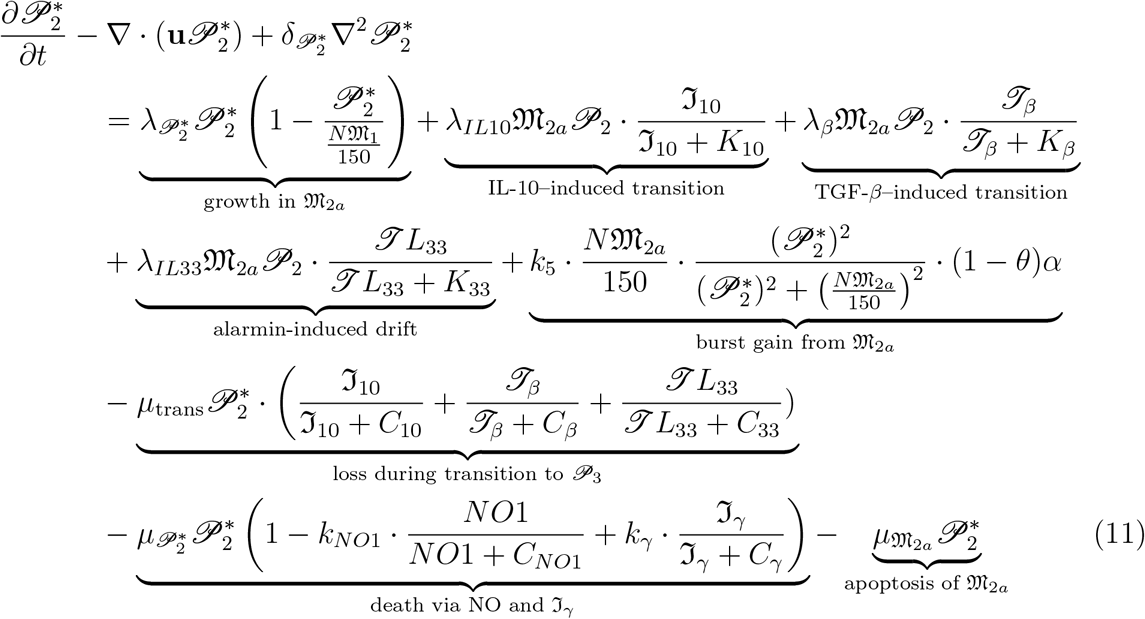

*P*_3_ represents the complete transition from the intermediate state 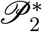 to the fully developed *P*_3_ state. This transition appears to follow a logistic growth pattern. When *M*_2*a*_ macrophages burst, they release 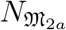 parasites, which include the matured *P*_3_ forms. These *P*_3_ parasites continue to reside within *M*_2*a*_. The transition is facilitated by specific cytokines, and unlike the previous equation describing 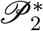, this stage includes a gain term that accounts for the accumulation of parasites during the transition. All remaining descriptions and biological context are provided in the section above.

Let us begin with 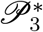, which represents the transitional parasite density residing within *M*_2*b*_ macrophages. This stage marks the transition from *P*_3_ to *P*_4_, and is biologically distinct from earlier phases. The gain in parasite density during this transition is driven by burst events originating from *M*_2*b*_. The cytokines involved in this process differ from those in previous stages and include ℑ*L* − 10, ℑ*L* − 6, *J NF* − (*α*), and *J LR* − 4, among others. However, there is also a loss component associated with the transition to *P*4. The resulting density *P*_4_ contributes to the development of more severe manifestations, such as post-kala-azar dermal leishmaniasis PKDL. The gain in *P*_4_ arises from the transition of 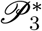, while the remaining terms governing its dynamics are analogous to those described for 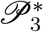.

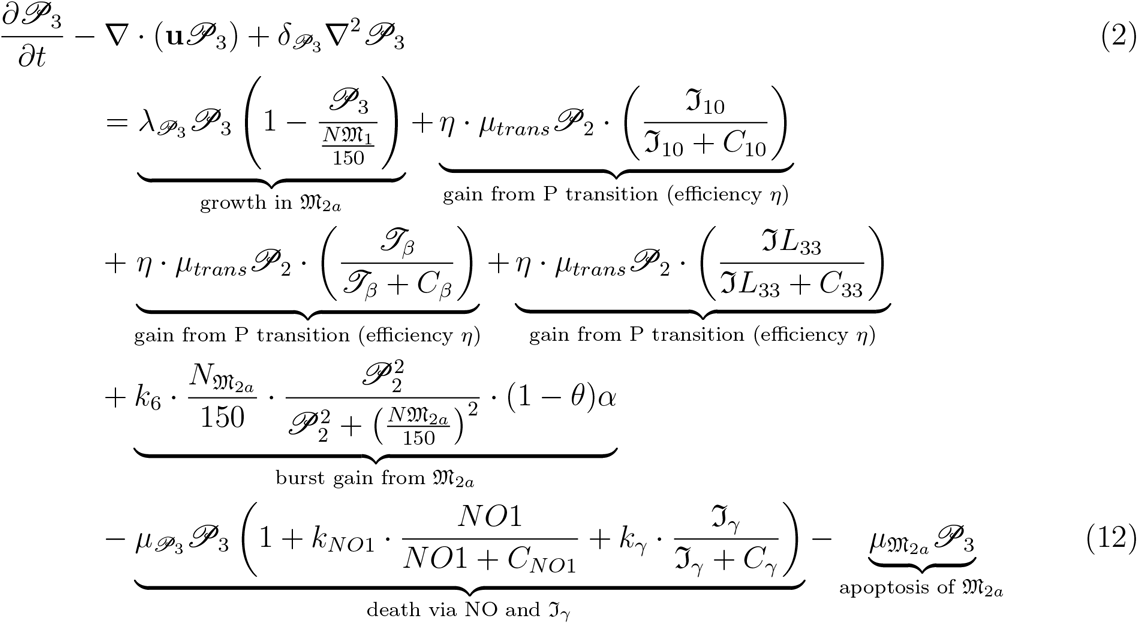

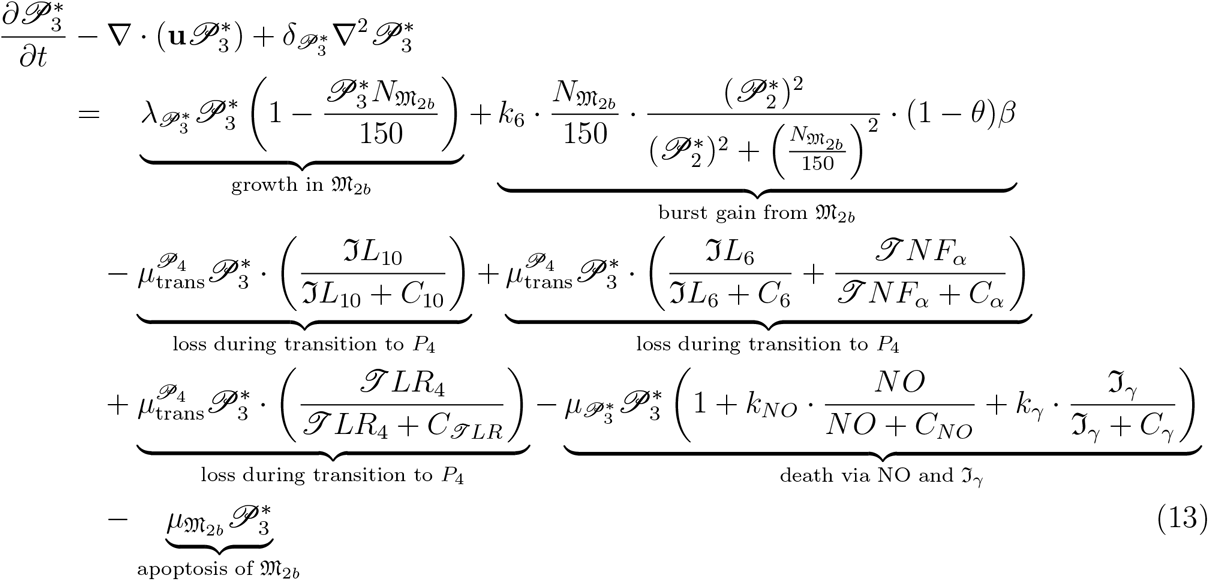

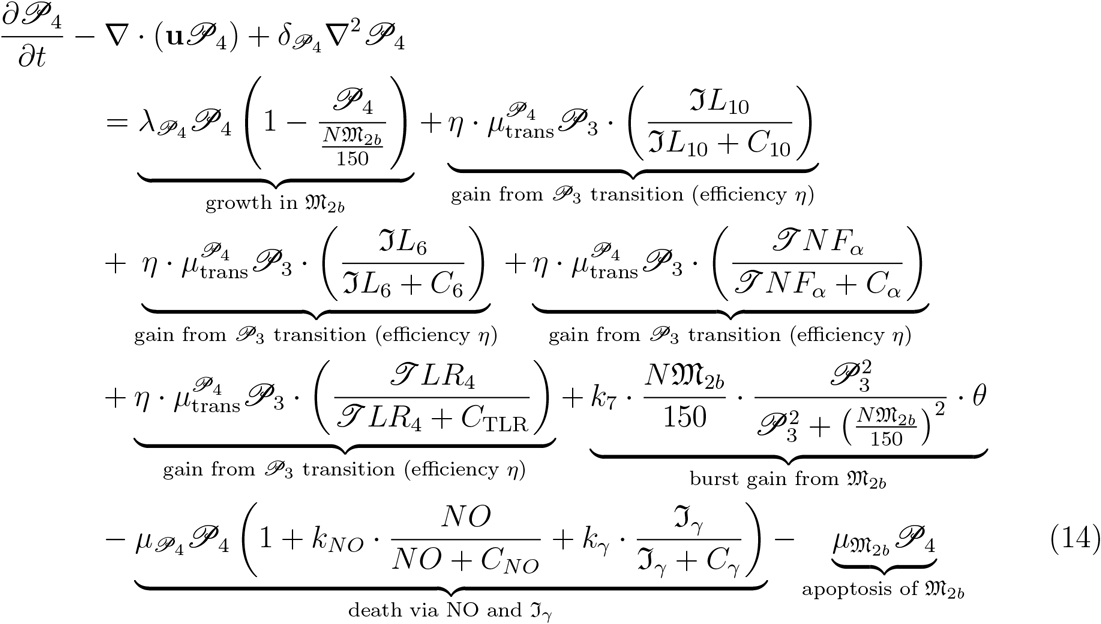

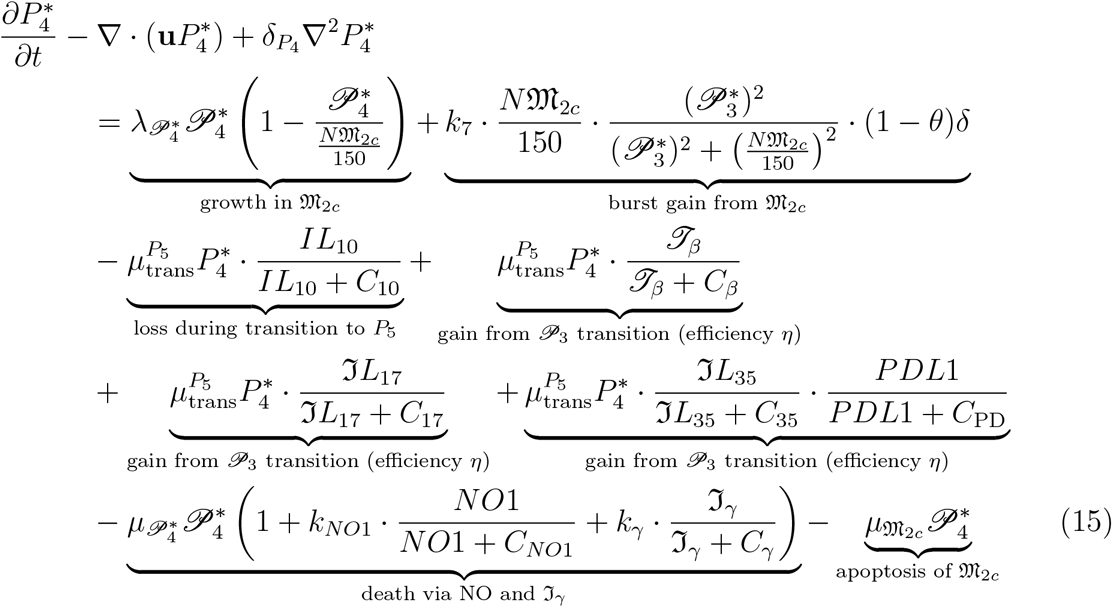

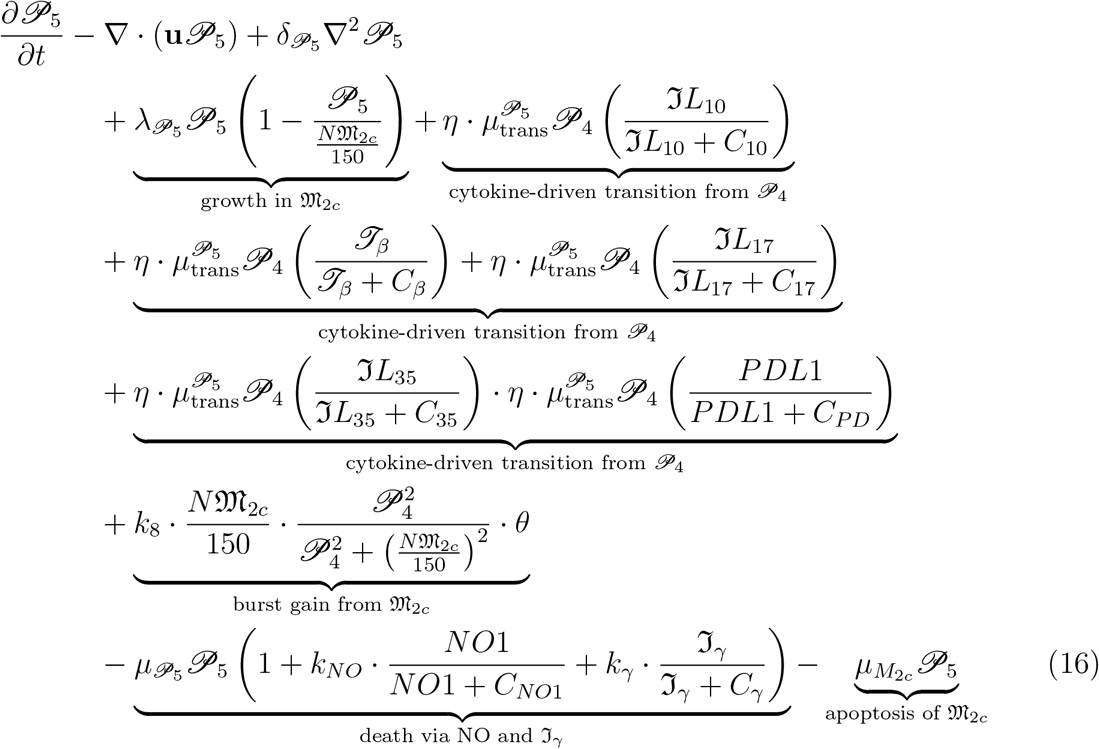

The transitional stage 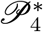 marks the beginning of the shift toward the more advanced parasite state, *P*5. If the density of parasites at this stage becomes excessively high, the severity of Leishmania infection also increases significantly. The transition from 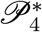 to *P*_5_ is mediated by a distinct cytokine profile, including ℑ*L* − 10, *J β*, ℑ*L* − 27, ℑ*L* − 35, and *PDL*1. These cytokines modulate the environment and contribute to the loss of parasites occurring specifically within the *P*_4_ population. Within *M*_2*c*_ macrophages, nitric oxide production is either very low or entirely absent, which further impairs parasite clearance. Apoptosis of *M*_2*c*_ macrophages may also occur during this stage.On the other hand, the emergence of *P*_5_ is driven by the transition dynamics of 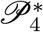, where a gain in parasite density is observed. This gain is directly linked to the cytokine-mediated progression from 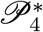 to *P*_5_.

### 2.5 Mathematical modelling of Cells

Mathematical modeling of immune cells such as *J*-cells, *B*-cells, *J*_1_-cells, *J h*2-cells, *J r*-cells, and *J h*17-cells focuses primarily on the diverse roles of *J*-cells in immune defense. *J* cells play a vital role in identifying foreign particles and recognizing specific pathogens. Except for *B* cells, the remaining cell types mentioned are subtypes of *J* cells, each with distinct properties:*J h*1 cells enhance cytokine production, which boosts macrophage activity and promotes pathogen destruction.*J h*2 cells are essential for defense against extracellular parasites.*J r* cells (regulatory *J* cells) maintain immune tolerance and prevent excessive inflammation.*J h*17 cells contribute to early-stage parasite clearance.*B* cells support the immune response by targeting pathogens that float freely in body fluids. The following equations describe the dynamics of these cell populations and their interactions.

#### 2.5.1 Mathematical model of T-cells

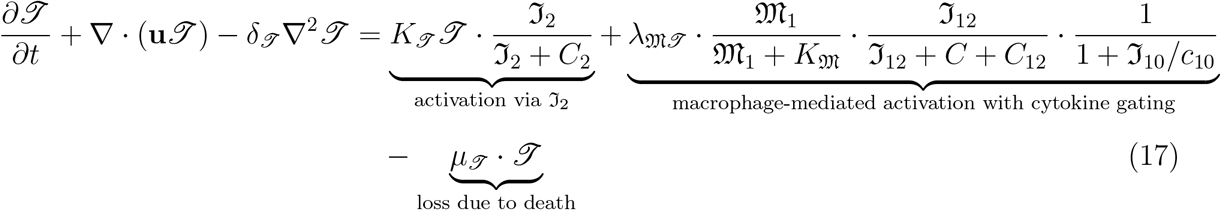

The equation describing *J* cell dynamics captures the activation of interleukin ℑ_2_, primarily through interactions with macrophages *M*_1_ within a cytokine environment enriched with ℑ_12_ and ℑ_10_. Ultimately, the model accounts for the natural death of *J* cells as a loss term [46, 47].

#### 2.5.2 Mathematical model of B-cells

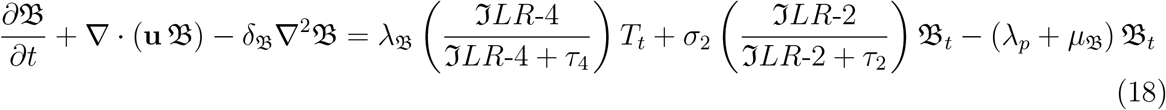

*B* cells begin functioning during the early stages of infection, activated through the combined influence of antigens and cytokines. Initial activation occurs via interleukin ℑ_2_, followed by antigen-driven stimulation mediated by ℑ_6_. Macrophage-mediated activation also plays a role, particularly through cytokines ℑ_6_ and ℑ_10_, within the *M*_2*a*_ macrophage environment. The natural loss of *B* cells is accounted for as part of their physiological turnover [87].

#### 2.5.3 Mathematical model of Th1 cells

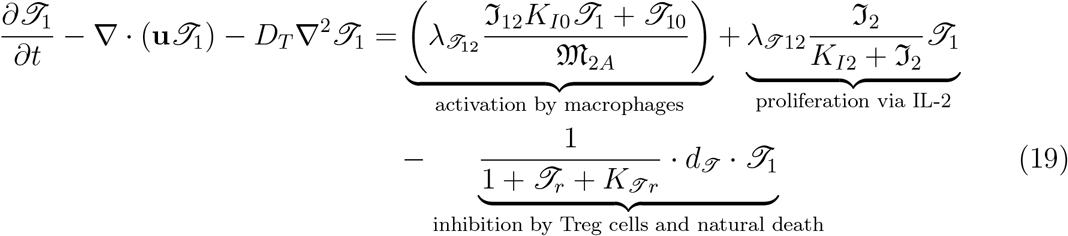

*J h*1 cells begin their activation through interactions with *M*_2*a*_ macrophages, primarily mediated by cytokines *J*_1_ and *J*_10_. The proliferation of *J*_1_ cells may occur via stimulation by interleukin ℑ*L ™*2. However, this process is subject to inhibition by regulatory *J* cells (Tregs), and natural cell death is also accounted for [94].

#### 2.5.4 Mathematical model of *J h*2 cells

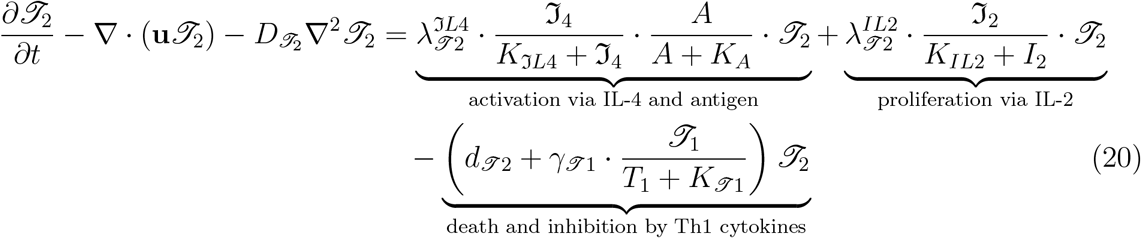

#### 2.5.5 Mathematical model of *J r* cells

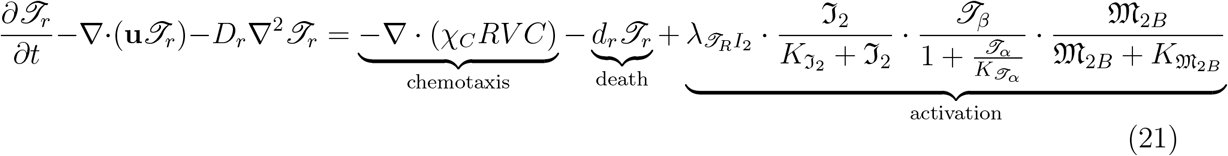

#### 2.5.6 Mathematical model of *J h*17 cells

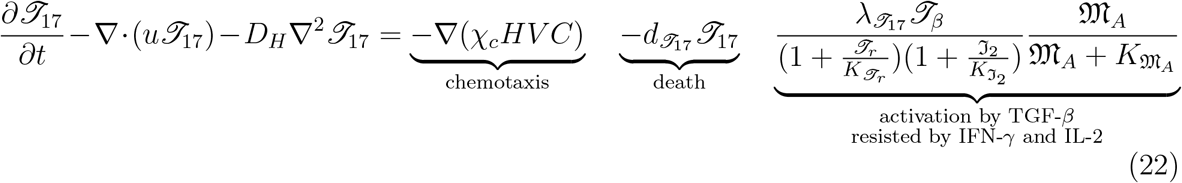

*J h*2 cells are activated via ℑ*L* − 4 in response to antigen exposure. Their proliferation is supported by ℑ*L*− 2, while death and inhibition are mediated by *J h*2-associated cytokines. Regulatory *J* cells Tr cells contribute to chemotaxis and are activated through *J β*, with their activity proportionally influenced by *J NFα* [94]. *J h*17 cells are activated in the presence of *J GF* − (*β*), but this activation is resisted by cytokines such as ℑ*FN* − (*γ*) and ℑ*L* − 2 [94].

### 2.6 cytokines without Drug

Cytokines are a diverse group of small signaling proteins that play a critical role in the regulation and coordination of immune responses. They are predominantly secreted by immune cells such as T lymphocytes, B lymphocytes, macrophages, and dendritic cells, and function as key mediators in the body’s defense mechanisms against infection and inflammation [48, 49, 51, 52, 53].

### 2.7 Mathematical model of Interleukin-2

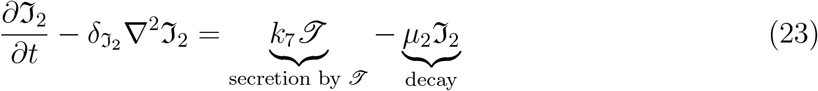

### 2.8 Mathematical model of Interleukin-10

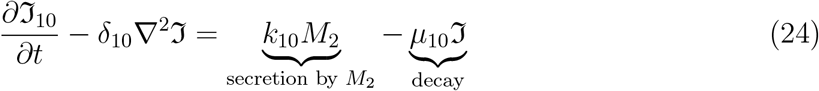

### 2.9 Mathematical model of Interleukin-12

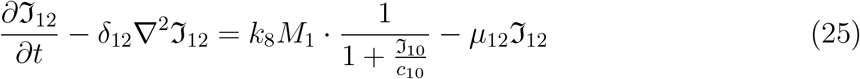

### 2.10 Mathematical model of Interferon-*γ*

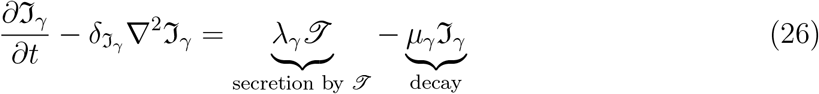

### 2.11 Mathematical Model of Nitric oxide NO and NO1

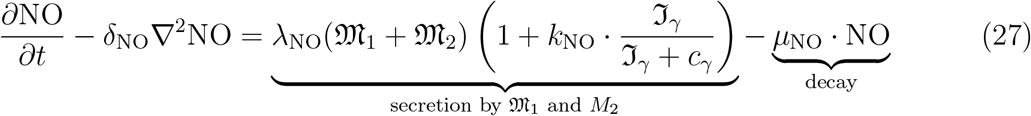

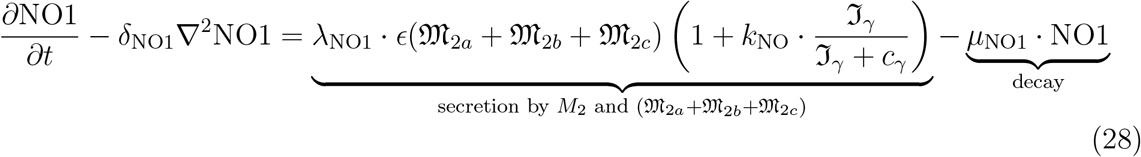

Nitric oxide is typically produced by *M*_1_ and *M*_2_ macrophages, primarily through stimulation by the cytokine *Iγ*. However, in our case, we additionally incorporate *M*_2*a*_, *M*_2*b*_, and *M*_2*c*_ macrophage subsets, which appear to produce either very low levels or no nitric oxide at all. To represent this behavior, we define the equation *NO*_1_ to capture the minimal or absent nitric oxide production in these macrophage types [54].

### 2.12 Mathematical Model of *J NF* − *α*(*J*_*α*_) **and** *J NF* − *β*(*J*_*β*_)

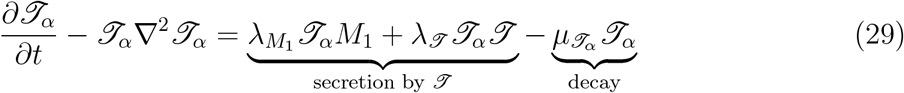

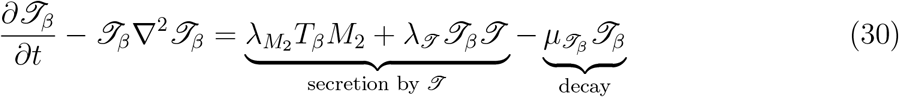

Tumor necrosis factor alpha (*T N* ℱ-*α*, denoted *J*_*α*_) is produced by *J* cells and M1 macrophages, while tumor necrosis factor beta (*T N* ℱ-*β*, denoted *J*_*β*_) is secreted by M2a macrophages [55, 56, 57, 58, 59, 94, 87, 88, 89, 90, 91, 92, 93, 94].

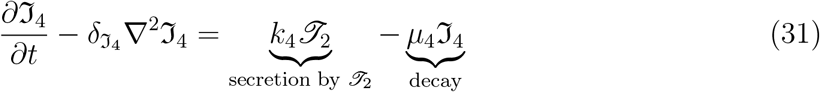

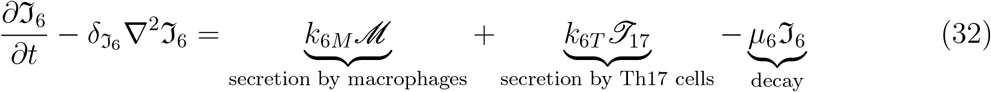

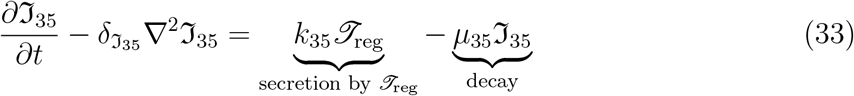

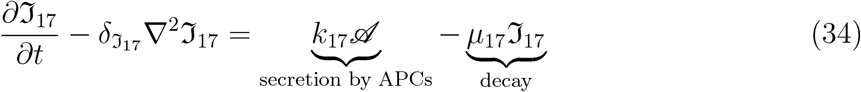

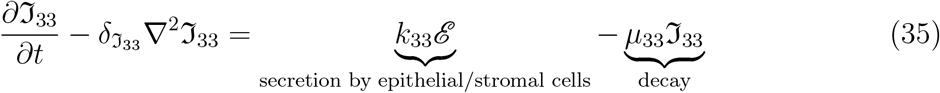

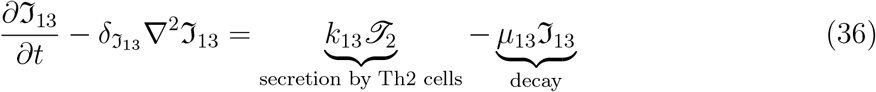

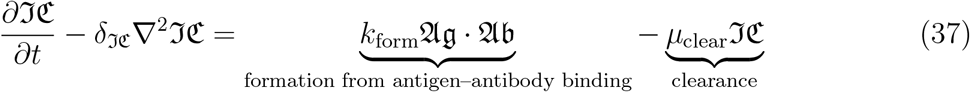

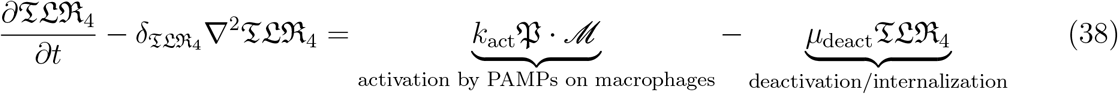

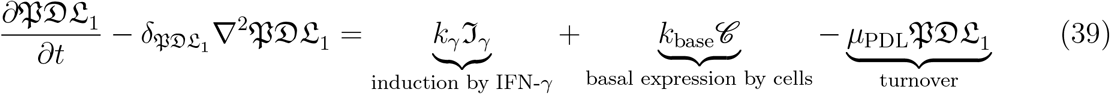

### 2.13 Equations for Nano-drugs Iron-oxide and cirtic acid Coated iron oxide

Equations (40, 41, and 42) represent the pulsed equations for iron oxide and citric acid coatings.

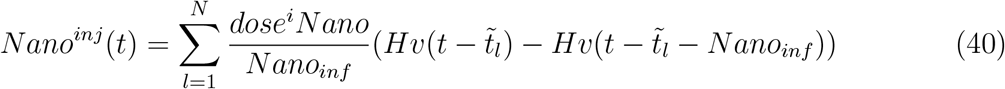

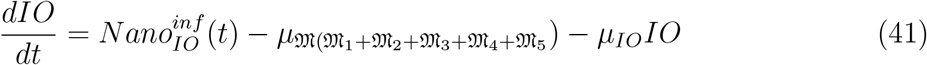

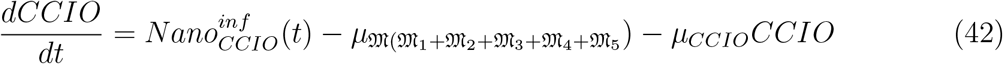

## 3 The equation for the velocity of cells u

To analyze computational complexity in a simplified framework,In this particular literature [16] they assume that all cells in the granuloma are distributed uniformly with a cellular density of 1 g/cm^3^. In addition, we consider the inclusion of B cells, which also contributes to a density of 1 g/cm^3^ within the granuloma.

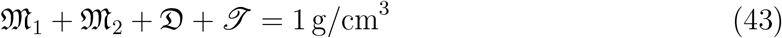

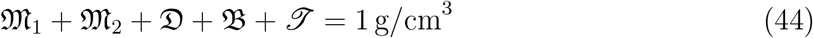

In the dispersion coefficients in Equation 43 are equal. However, this assumption may not always hold, particularly in cases where D cells are greater than T cells, which could influence their diffusion behavior. In another scenario, we assume uniform cellular density and additionally incorporate B cells into the system.

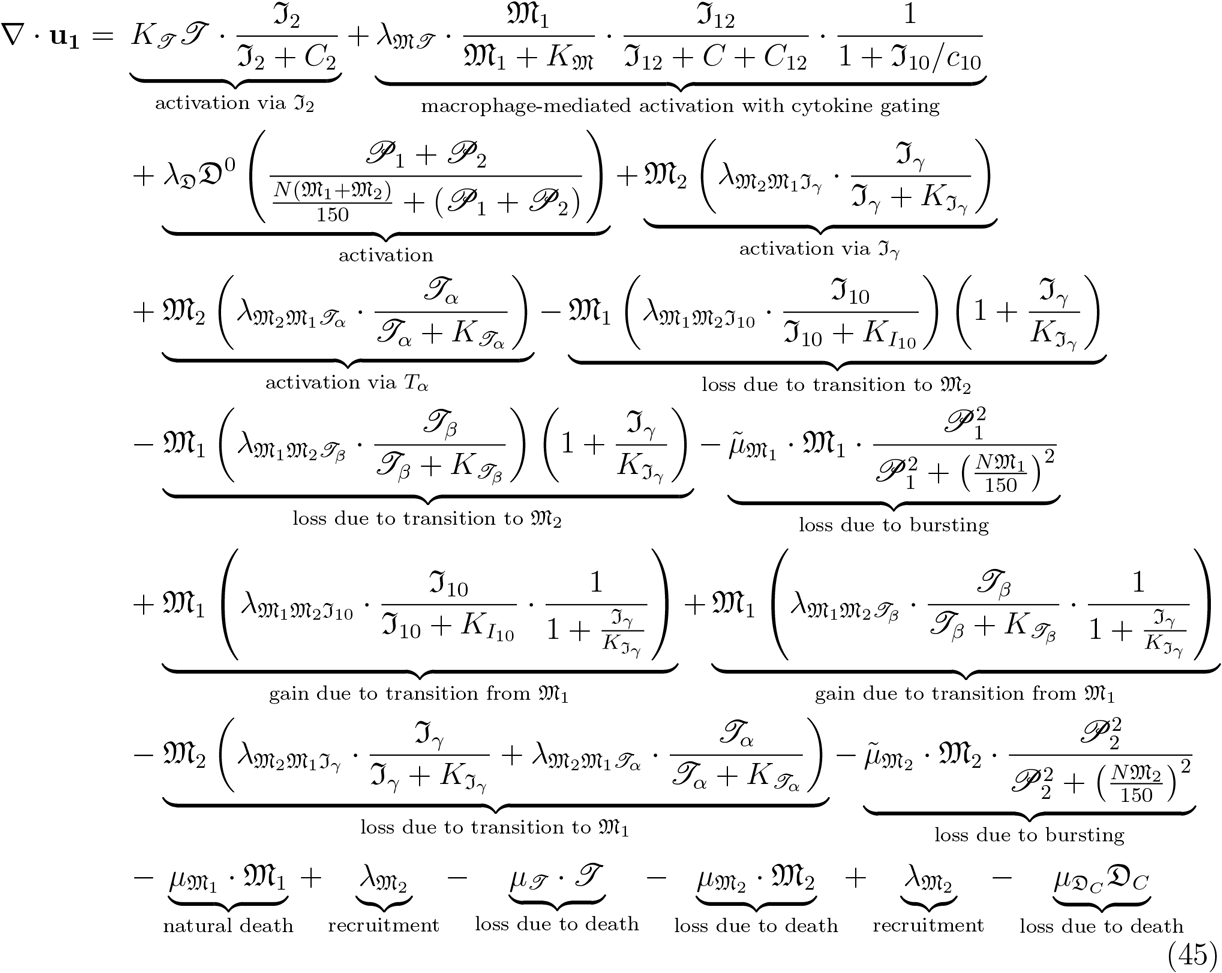

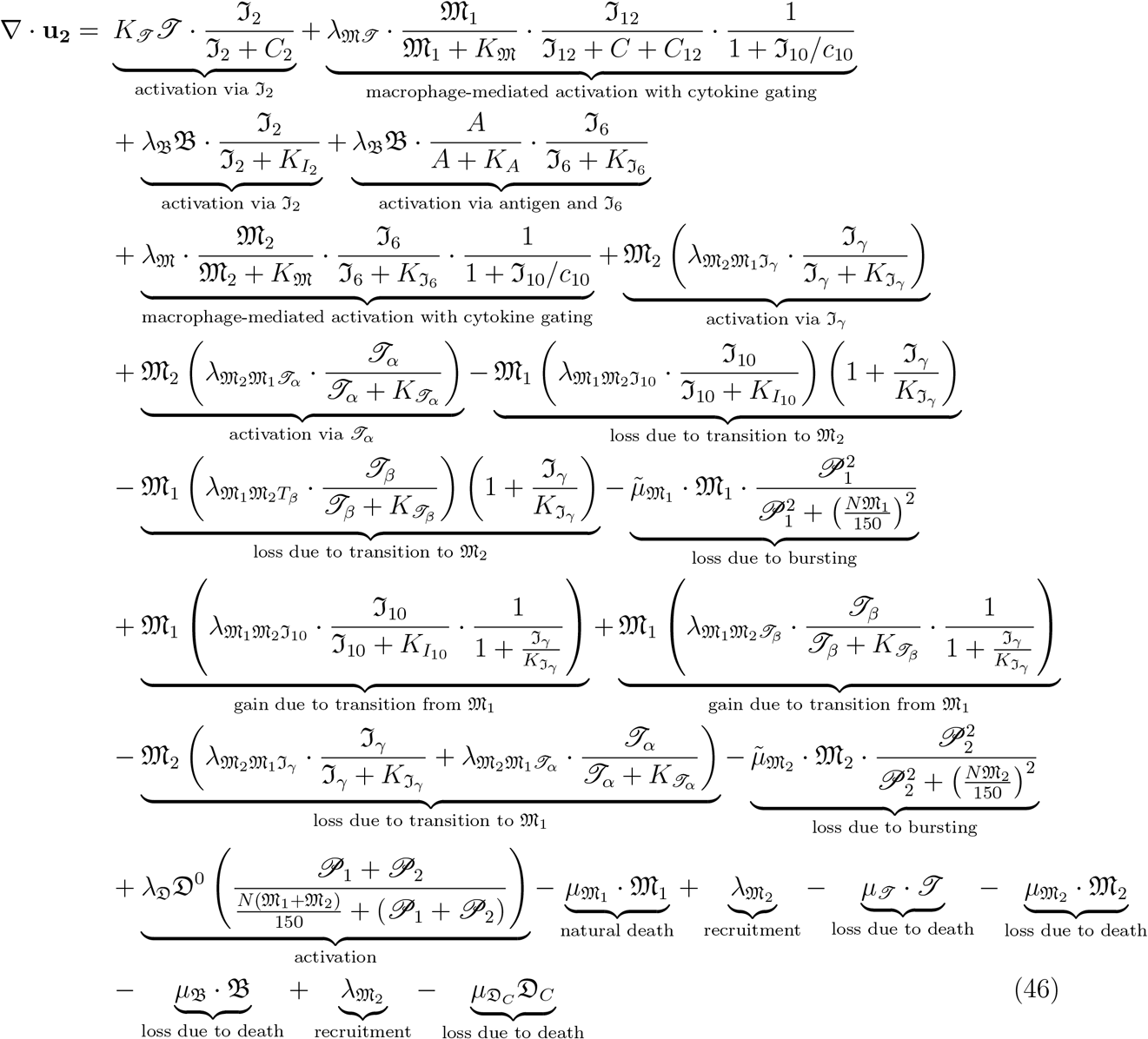

## 4 Boundary Condition

The shape of the granuloma is assumed to be spherical. We consider the radius as a function of time, denoted by *r* = ℜ(*t*), and assume that all variables are radially symmetric. Therefore, they are functions of *r* and *t*, i.e., *f* = *f*(*r, t*). At the center of the granuloma *r* = 0, each *r*-th derivative of any variable is zero due to symmetry.

Let the average number of parasites in each *M*_1_ macrophage be *N*_1_, and in each *M*_2_ macrophage be *N*_2_, where *N*_1_ *< N*_2_. *M*_1_ macrophages are considered healthier than *M*_2_ macrophages. However, despite containing fewer parasites, *M*_1_ macrophages migrate into the granuloma. The immigration and arrival of *M*_1_ macrophages at the boundary increase with the factor 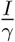.

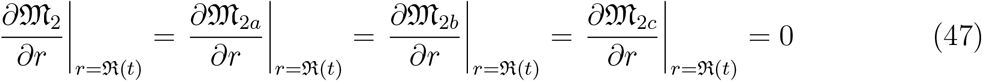

The strength of the adaptive immune response is influenced by *β*, a phenomenological parameter that represents the intensity of migration from the lymph nodes. This migration involves macrophages and *J* cells entering the granuloma. *M*_0_ denotes the density of macrophages, which is assumed to remain constant. In particular, within the lymph nodes associated with the granuloma, dendritic cells are in motion under flux conditions.

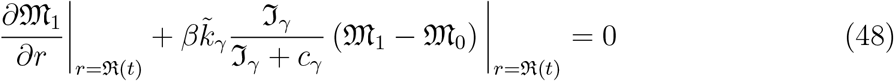

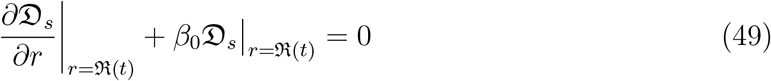

We denote by *J*_0_ the density of inactive *J* cells migrating from the lymph nodes. *J*_0_ is assumed to be constant. Some of the *J*_0_ cells are activated at the granuloma boundary by dendritic cells under an ℑ_12_ cytokine environment.

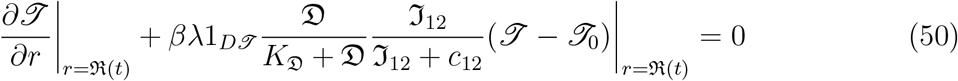

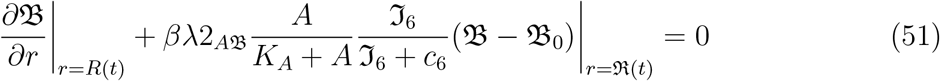

Similarly, *B*_0_ denotes the inactive state of *B* cells, which is also assumed to be constant. *B*_0_ cells are activated at the granuloma boundary by dendritic cells in the presence of antigen and an *J*_6_ cytokine environment.

We assume a no-flux boundary condition for following paarsites densities

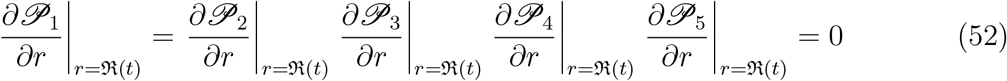

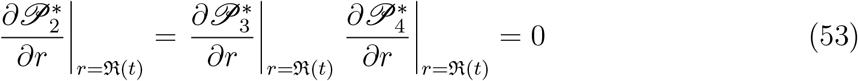

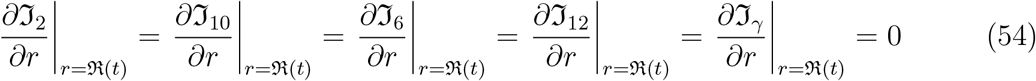

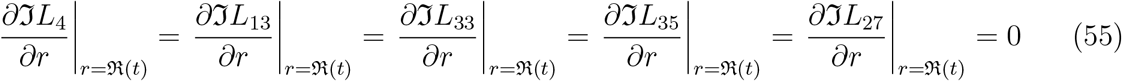

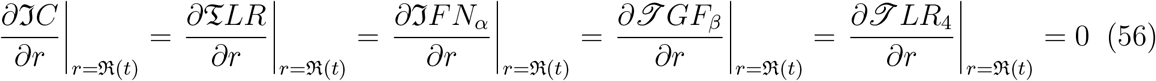

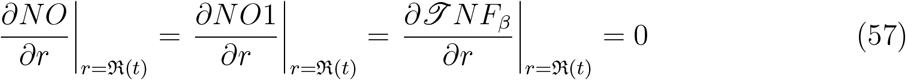

We also assume that all cytokines are produced by immune cells within the granuloma, as well as by those located outside the granuloma. The granuloma boundary is represented by *r* = ℜ(*t*).

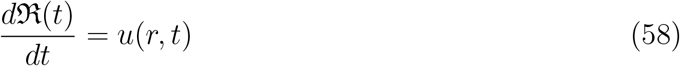

## 5 Moving Boundary condition for three phase

The Stefan problems describe moving boundaries in systems governed by differential equations [61, 62]. However, most formulations are heavily dependent on the specific nature of the problem—sometimes influenced by the discretizations scheme—which can lead to physically unrealistic interpretations. Interface problems typically arise at two-phase boundaries, as clearly explained in [63, 64, 65, 66]. Generally, the concentration in one phase is fixed by a thermodynamic constraint. The solute then diffuses toward the interface through Phase A, while Phase B is removed. These fluxes are typically unequal. In our model, we consider three distinct concentrations based on the densities of healthy macrophages, infected macrophages, and nano-drugs. Specifically: **Phase A** contains the number of healthy macrophages.**Phase B** includes the number of infected macrophages, i.e., macrophages affected by parasites. **Phase C** represents a reactive phase that eliminates parasites within infected macrophages, depending on the concentration of nano-drugs [67, 68, 69, 70]. These concentrations are typically unequal across the three phases.So far *s* = *s*(*t*) is the interface position.However in our case we design *S* = *S*_1_(*t*) and *S* = *S*_2_(*t*) are two moving boundaries

The following system of differential equations can be formulated to describe the dynamics across Phases A, B, and C:

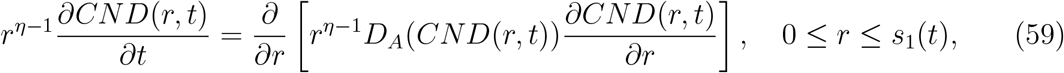

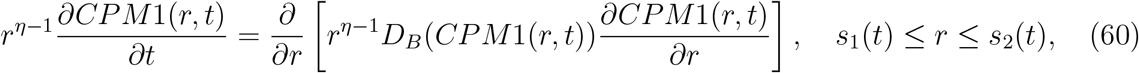

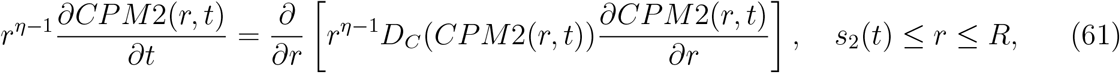

Equations (59)–(61) describe the diffusion dynamics across three distinct phases. Equation (59) governs diffusion in Phase A, located to the left of the interface. Equation (60) pertains to diffusion within the central region, Phase B. Finally, Equation (61) addresses diffusion to the right of the interface, represented as Phase C. This paper defines the moving boundary dynamics of drug diffusion into infected macrophages. We consider three equilibrium concentrations:*CPM*_1_, CND, and *CPM*_2_. Specifically, *CPM*_2_ denotes the concentration of healthy macrophages, CND represents the concentration of the nano-drug, and *CPM*_2_ corresponds to the concentration of infected macrophages. To complete the formulation of diffusion-controlled dynamics across all three moving phases, fixed boundary conditions are imposed at *r* = 0 and *r* = *R*, with moving interfaces at *r* = *S*_1_(*t*) and *r* = *S*_2_(*t*). The model assumes that the diffusion coefficients *D*_*A*_, *D*_*B*_, and *D*_*C*_ for Phases A, B, and C, respectively, are independent of concentration.

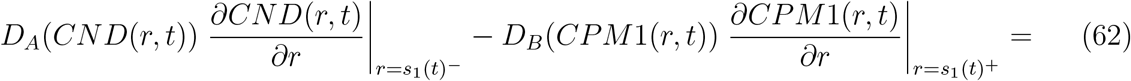

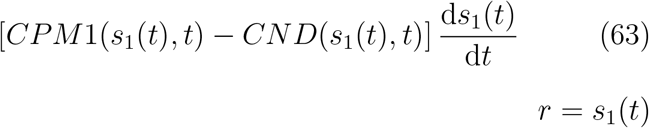

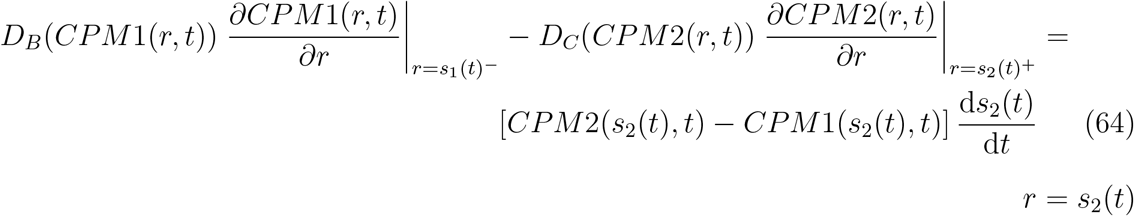

For all three phases of the moving boundary problem, we extend the Landau transformation to the coordinate system. When the spatial boundaries are fixed, we model the diffusion-controlled phase based on nano-drug dynamics. The numerical techniques employed here were originally developed by Murray and Landis (1959). The following equations involve multiple interphase boundaries and concentration-dependent diffusion coefficients [71, 72, 73].

**Phase A** is defined with a boundary between 0 ≤ *r* ≤ *s*_1_(*t*). Here we define Phase A: 0 ≤ *r* ≤ *s*_1_(*t*),

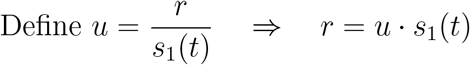

This mapping implies: *r* = 0 ⇒ *u* = 0, *r* = *s*_1_*(t)* ⇒ *u* = 1

Then **Phase B**: in between *s*_1_(*t*) ≤ *r* ≤ *s*_2_(*t*).

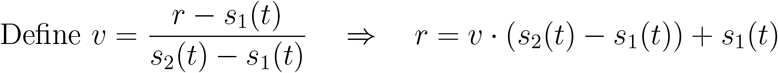

This mapping implies: *r* = *s*_1_(*t*) ⇒ *v* = 0, *r* = *s*_2_(*t*) ⇒ *v* = 1

**Phase C**: *s*_2_(*t*) ≤ *r* ≤ *R*.

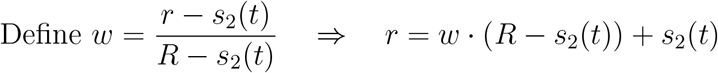

This mapping implies: *r* = *s*_2_(*t*) ⇒ *w* = 0, *r* = *R* ⇒ *w* = 1

In phase A the diffusion equation (59) becomes

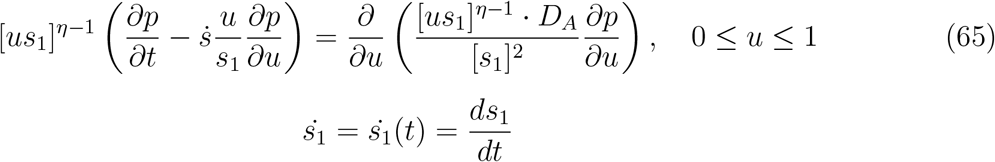

In phase B the diffusion equation (60) becomes **For Phase B**, define the transformation:

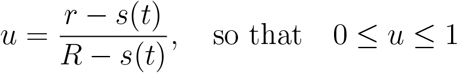

Let *q*(*u, t*) = *c*(*r, t*) be the concentration in the transformed coordinate. Then the diffusion equation becomes:

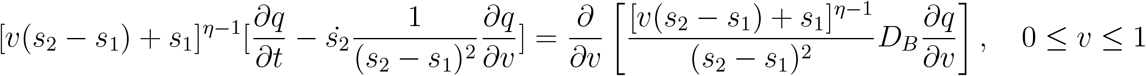

In phase C the diffusion coefficient equation (61) becomes

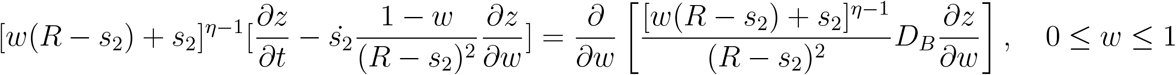

The transformed version of the interface equation (62) and (63) becomes.

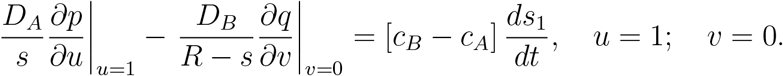

The transformed version of the interface equation (61) becomes

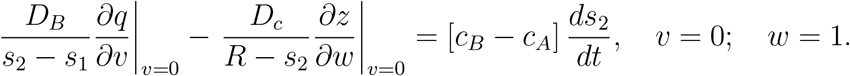

A conservative scheme Derivation At any time t,this quantity is given by

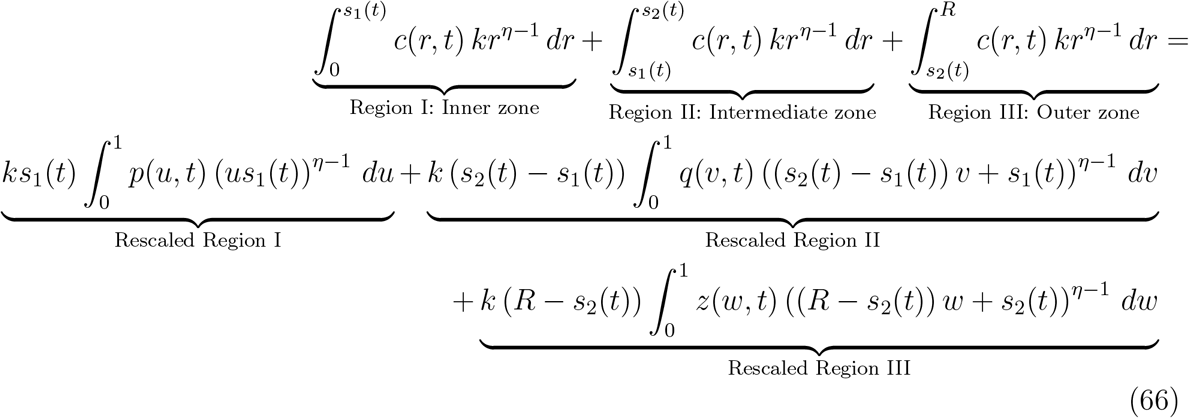

This value must be time invariant

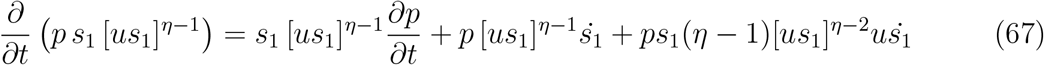

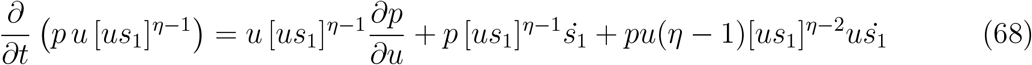

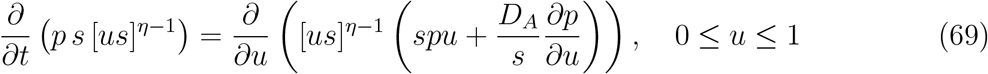

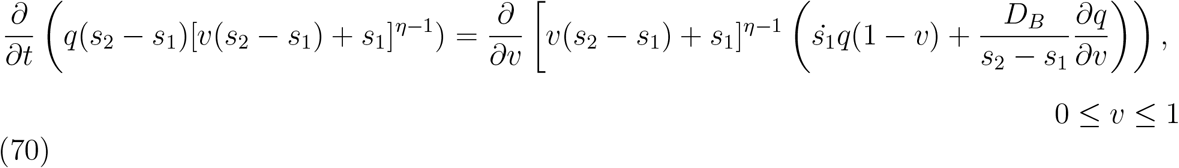

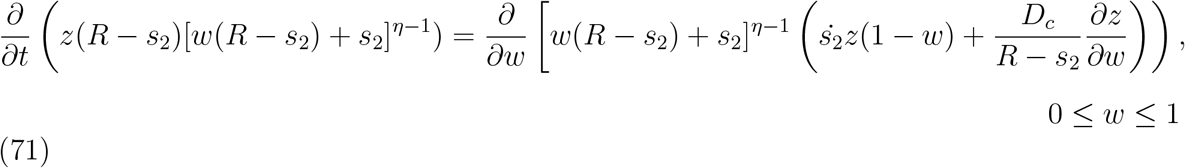

Now integrate the divergent form of the diffusion equations (67) (68) around each node over one time step in phase

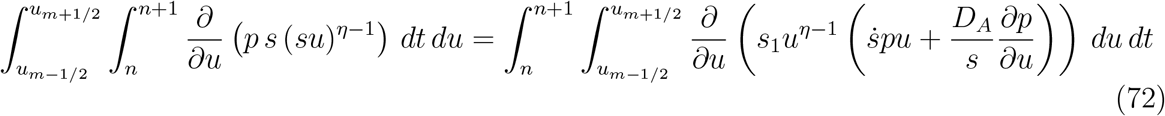

can be written as

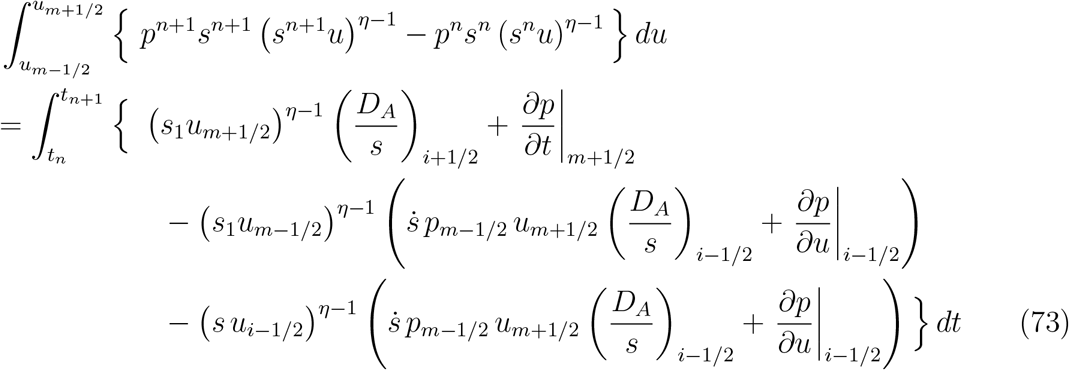

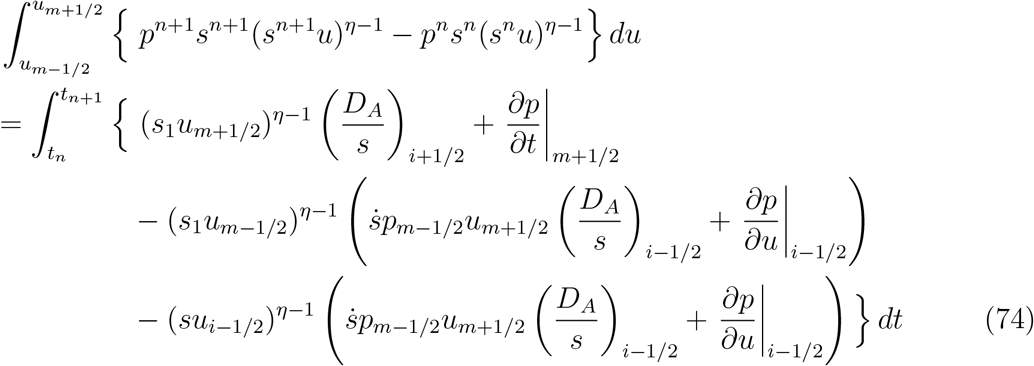

Taking a constant 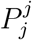 to approximate the variable *P* ^*m*^ = *p*(*r, t*^*m*^) over the interval

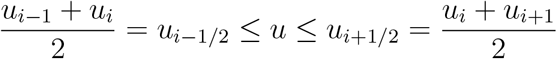

and it is similar for 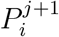, it is possible to simplify the integral on the left-hand side of the equation 73.

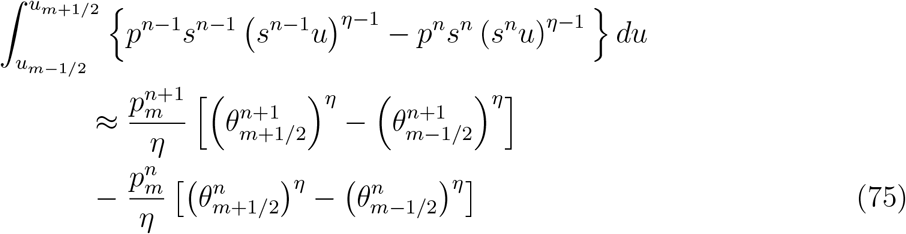

Let the notation 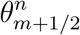 represent *S*^*j*^*u*_*i*+1*/*2_.

To discretize the right-hand side, introduce *σ* ∈ [0, 1], and define: 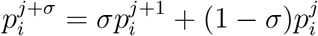. This expression approximates the concentration near *u*_*i*_. over the interval [*t*^*j*^, *t*^*j*+1^].

The integral can then be evaluated to yield the following equations represent the finite difference scheme

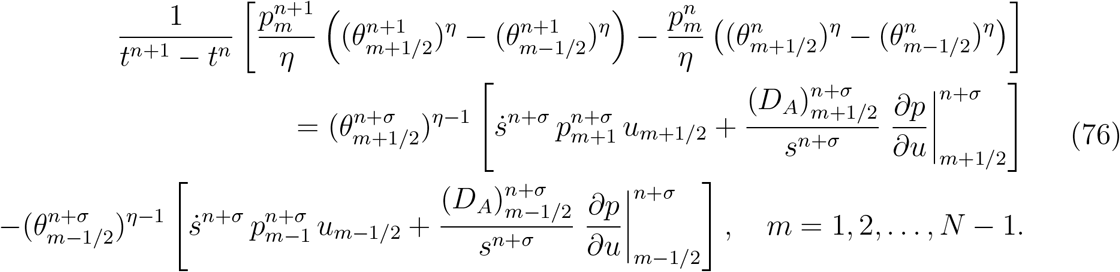

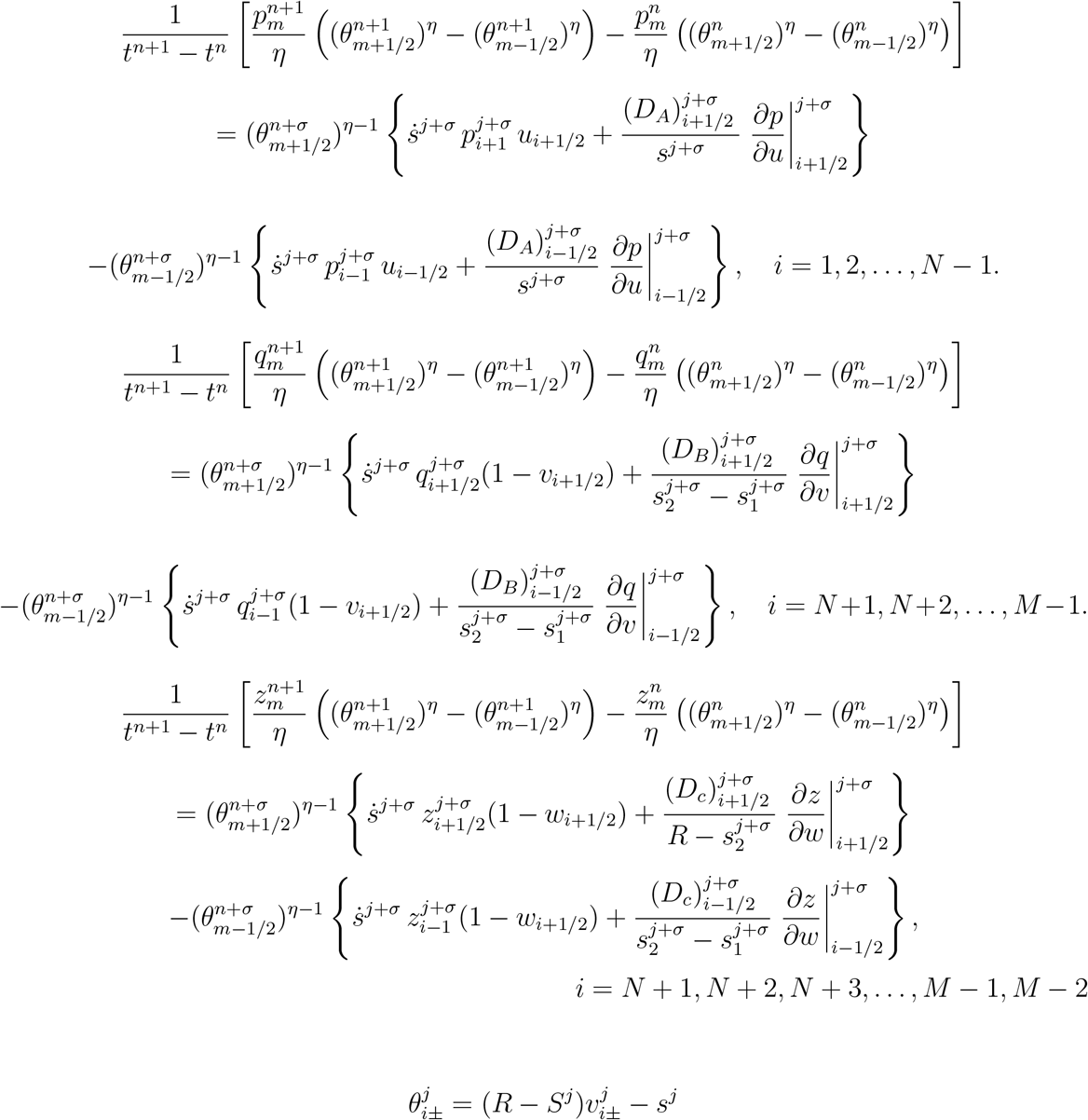

The quantity 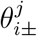 is the interval between *t* = *t*^*j*^ and *t* = *t*^*j*+1^ is the amount of solute in element *i*. To generate a finite difference form of the conservation equation (73), it is also necessary to derive the finite difference approximations for the concentrations at the given boundaries, specifically at *i* = 0, *i* = *N*, and *i* = *M*.At the moving interface,the local equilibrium implies:

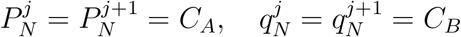

For the fixed boundaries, zero flux conditions are imposed:

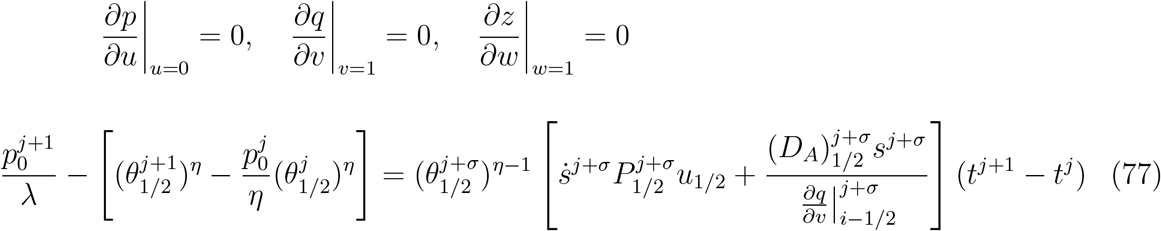

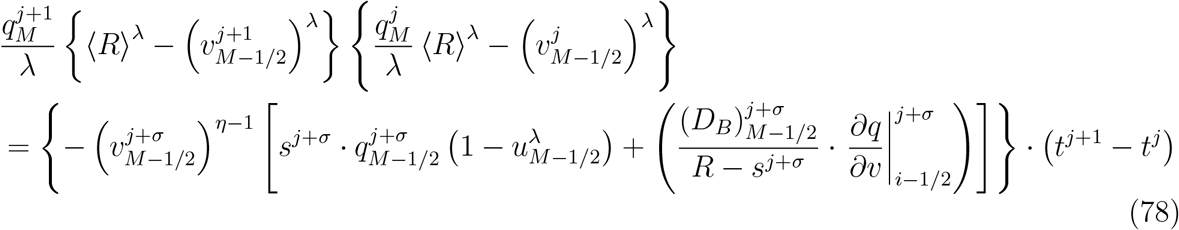

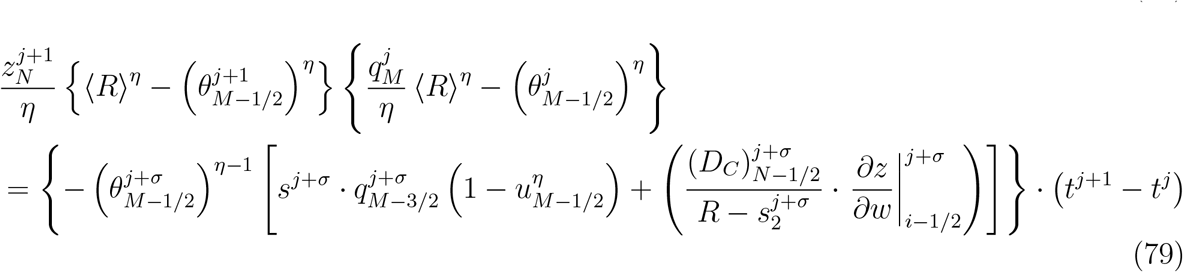

these equations 77,78 and 79 can be written as

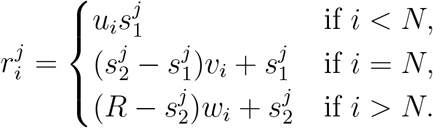

To discretise the interface equation

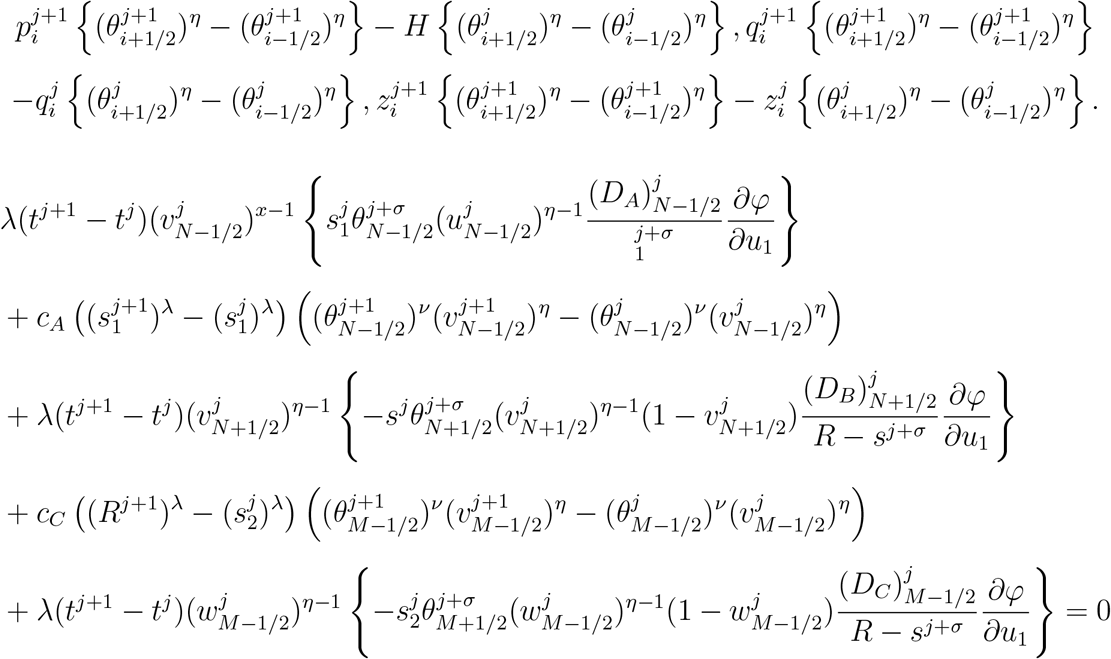

## 6 Implicit and Semi-Implicit Modeling in Polar Coordinates

This section presents drug diffusion using implicit and semi-implicit formulations. We predict drug flow in circular polar coordinates [74]. A detailed study of gradient flow in the formation of biological transport networks—via implicit and semi-implicit formulations—was previously developed using the Cai-Hu model [86]. This model demonstrates unconditional energy decay with respect to mesh size. In our approach, we investigate implicit and semi-implicit energy decay based on a moving mesh, which enables more precise analysis of mesh size compared to the previous model. We introduced circular polar coordinates to better capture the radial symmetry observed in granuloma formation during Leishmaniasis. This coordinate system aligns naturally with the biological geometry, making it well-suited for modeling drug transport in such contexts.

Here, *D*^Λ*m*^ represents a **stiff reaction-diffusion system** in a nonlinear and non-differentiable physical form. This is particularly relevant for cases where 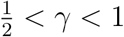, based on the **metabolism term** |*m*|^2^(*γ* − 1), which becomes stiff as *m* → 0. The **activation term** *C*^2^∇*p* ∇*P* has eigenvalues of zero, leading to *C*^2^|∇*P*|^2^ when |∇*P*| is large. The reference [74] also explains the **implicit and semi-implicit schemes** using a **rectangular grid**, where (∇_*h*_)^2^ denotes the **discretized Laplacian operator**.

### 6.1 Implicit Nonlinear Equations in Polar Coordinates

In our model, we consider the polar coordinates (*r, θ*), Here *r* represents the distance for radial and theta *θ* denotes an angular position. We introduce factors involving 1*/r*, which are characteristic of a two-dimensional polar coordinate system. This framework allows us to classify the fields into two types: the electric field *E*(*r, θ*) and the magnetic field *H*(*r, θ*). The governing equations are expressed in divergence form.

The governing equation is written in **divergence form**:

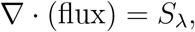

This indicates the conservation of a quantity with a source term *S*_*λ*_. The left hand side of equation (80) combines the derivatives of 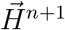 and ^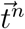^. The second term is radially weighted and involves the material coefficients 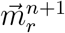 and 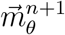, which act on the derivatives of 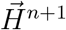. The time levels *n* and *n* + 1 are coupled through the average 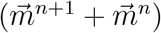. The field 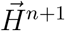 appears within both the derivative terms and the divergence operator, and it is updated at each time step using numerical schemes [75, 76, 77, 78, 79].

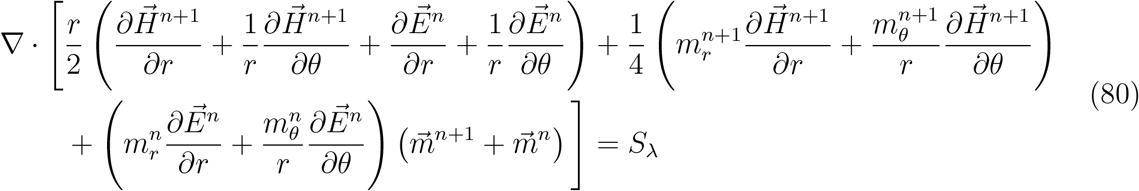

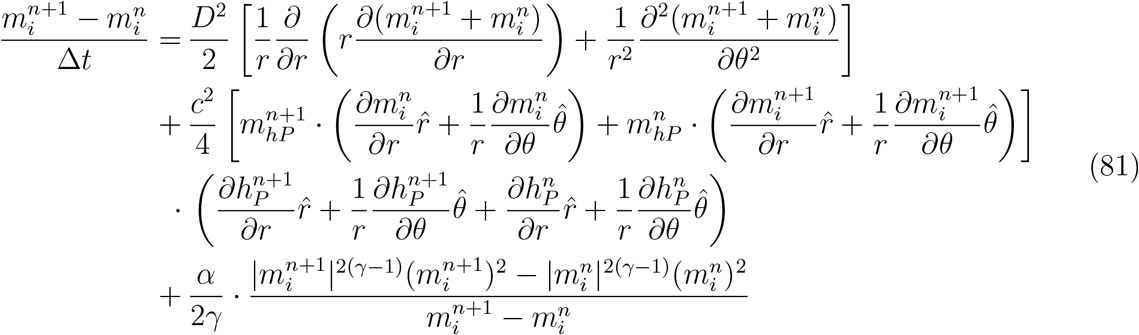

### 6.2 Semi-Implicit Nonlinear Equations in Polar Coordinates

In this model, we incorporate a semi-implicit function that plays a dual role—acting both implicitly and explicitly. We consider semi-implicit nonlinear equations in polar coordinates, involving two scalar fields *m*_1_(*r, θ, t*) and *m*_2_(*r, θ, t*). Nonlinear multipliers such as 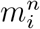 or 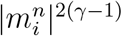 are treated explicitly at the previous time level *n*. The term *D*^2^ represents drug diffusion, while *c*^2^ captures nonlinear interactions driven by curvature. These curvature effects are both self-dependent and involve cross-interactions, which are relevant in contexts such as magnetization dynamics, elasticity, and nonlinear wave coupling. Let *m*_1_(*r, θ, t*) and *m*_2_(*r, θ, t*) be two coupled scalar fields defined on a circular domain. The nonlinear reaction term is given by:

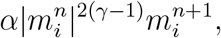

where *γ* controls the degree of nonlinearity and *α* determines the strength and sign of the interaction. This term is treated semi-implicitly, with *m*^*n*+1^ appearing implicitly.

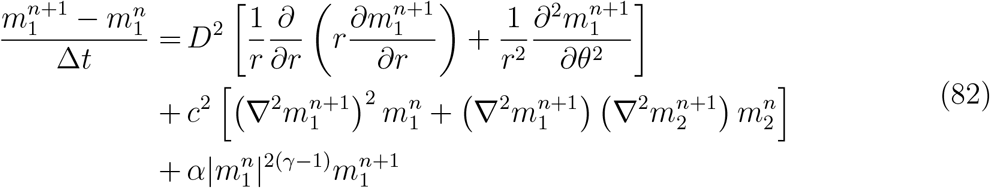

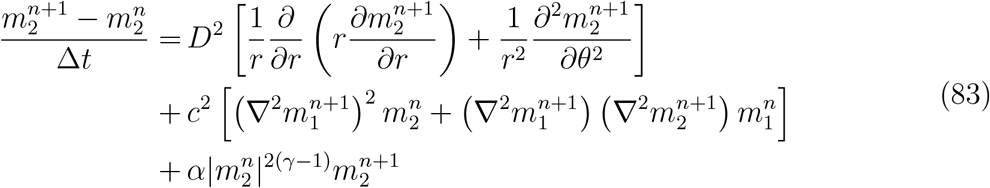

The semi-implicit terms include the component *m*, 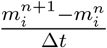 which represents the time derivative evaluated across time levels *n* and *n* + 1, 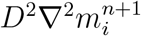 accounting for diffusion and smoothing at the time level *n*+1. We further consider nonlinear curvature coupling,*c*^2^(∇^2^*m*)^2^*m*^*n*^ where the Laplacian is evaluated 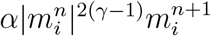 at the time level *n* + 1 and the field at time level *n*. Then the nonlinear reaction term is treated semi-implicitly in time.

In reference to mesh movement, we adopt a mesh-with-submesh approach. The following steps are applied using an *r*-moving mesh, where Poisson pressure is introduced. Here, MST denotes the mass source term, and 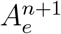 represents the area of element *e* at time step *n* + 1. The superscripts inj vary from 1 to 3, while the index *j* also ranges from 1 to *j*. In the *j*^th^ coordinate direction, the mesh velocity computation, MST source term, and Poisson pressure equation are clearly explained in the equation given below [80]. The semi-implicit time stepping scheme is given by:Here, ∇^2^*m* denotes the Laplacian in polar coordinates:

#### Mesh Velocity Computation

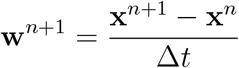

#### MST Source Term

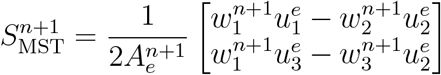

#### Pressure Poisson Equation

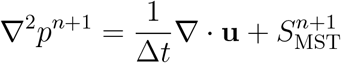

## 7 Results and Discussion based on Nano-drugs Ironoxide and Critic acid coated Iron-oxide

The host cells for *Leishmania* parasites are Macrophages which plays a important role during infection. In our study, we focused on iron oxide-based nanomedicine for leishmaniasis, with particular emphasis on nano-drugs. Among the candidates, PEI_25_-decorated Fe_3_O_4_ and citric acid-coated iron oxide nanoparticles emerged as our primary focus. This work clearly explore the interplay between the macrophages, *Leishmania*, and iron oxide. Although numerous drugs are currently under trial, there remains no approved treatment for visceral leishmaniasis. Our central hypothesis is that nano-drugs offer a highly targeted approach for infected macrophages, making them ideal candidates for therapeutic intervention.One of the most critical battles between *Leishmania* and macrophages during that scenario, the iron is the competitor element for the survival as well as death. *Leishmania* has evolved sophisticated mechanisms to acquire iron necessary for its proliferation. To counter this, we aim to increase intracellular iron levels in macrophages, leveraging the closerelationship between cellular iron metabolism and antimicrobial mechanisms, particularly those based on reactive oxygen species (ROS).

Iron, a transition metal, exhibits the multiple oxidation states ranging from –2 to +7. It is almost involved in various biological functions, including oxygen transport and storage, oxygen utilization in muscle tissue, DNA synthesis, cellular respiration, and electron transport. Its key biological utility lies in the reversible conversion between its most common and biologically relevant forms: divalent ferrous iron (Fe^2+^) and trivalent ferric iron (Fe^3+^). This redox flexibility enables iron to act as a catalyst by readily accepting or donating electrons, thereby playing a central role in host-pathogen dynamics and therapeutic strategies.

Iron oxide nanoparticles (IONPs) are widely used in medicine for both diagnostic and therapeutic applications due to their ability to reveal intricate details of tissues and disease conditions. IONPs coated with carboxydextran have been approved as imaging contrast agents for the detection of liver lesions. The biodistribution of IONPs is influenced by several factors, including particle size, shape, surface coating, charge, dosage, and route of administration.Owing to their ability to selectively accumulate in macrophages—where *Leishmania* parasites reside—and the critical role of intracellular iron balance in regulating macrophage polarization, IONPs have emerged as a promising strategy for treating leishmaniasis. However, this approach presents certain limitations. Beyond the accessibility of IONPs to infected macrophages via parenteral or local administration, a key challenge lies in targeting macrophages as an immunomodulatory strategy for leishmaniasis therapy. Based on these considerations, we designed and tested our model using iron oxide nanoparticles to evaluate their therapeutic potential in the context of macrophage-targeted treatment for visceral leishmaniasis [95].

### 7.1 Energy Decay and Implicit Behavior of Iron Oxide Nanoparticles

In this section, we discuss the results concerning the implicit and semi-implicit functions applied to PEI_25_-decorated Fe_2_O_3_ and citric acid-coated Fe_3_O_4_. This discussion is particularly relevant for iron oxide-based nanodrugs targeting leishmaniasis. The detailed formulation of the implicit scheme in polar coordinates is provided in Section 6. The results demonstrate that the energy dynamics vary significantly depending on the nanoparticle formulation and the numerical scheme applied. The implicit treatment enhances stability and resolution, especially under conditions of high diffusion and pressure coupling. The behavior of iron oxide nanoparticles (IONPs) was studied in a dynamic, biologically inspired environment by comparing two formulations: PEI_25_-decorated *γ*-Fe_2_O_3_ and citric acid-coated Fe_3_O_4_. An initial energy level was assumed, and its decay over time was defined. A key focus of the study was the role of the drug diffusion coefficient, which differs between the two nanoparticle types.

Figure 4 presents the angular distribution of normalized intensity for PEI 25-coated *γ*-Fe_2_O_3_ nanoparticles with a mean diameter of 45 nm. The system was simulated from time step 0 to 99 at an energy level of 5.00 *µ*m. The polar grid spans radial divisions from 100 to 360 degrees. Figure 4**(A)** shows the distribution at time step 0, with energy 5.00 *µ*m. The polar grid and energy decay were modeled using implicit functions. The intensity, denoted as *m*_*i*_ intensity, ranges from 0 to 0.9063. A peak intensity is observed within a narrow angular sector. Although the implicit domain spans from −0.25 to 0.25, the high intensity remains confined within this range, indicating minimal energy dissipation. Figure 4**(B)** shows the *m*_*i*_ intensity distribution at time step 1, ranging from 0 to 0.9061, with an energy decay to 4.76. The simulation was performed on a 95×344 grid, and the intensity appears lower compared to subfigure A. Figure 4**(C)** corresponds to time step 50, where *m*_*i*_ intensity spans from 0 to 0.6933. The energy has decayed to 0.41, indicating a gradual loss over time. The grid resolution used was 17×62. Figure 4**(D)** represents the final time step (t = 99), with *m*_*i*_ intensity ranging from 0 to 0.4233. The energy has decayed significantly to 0.04, and the simulation was conducted on a 10×38 grid.

**Figure 4.**
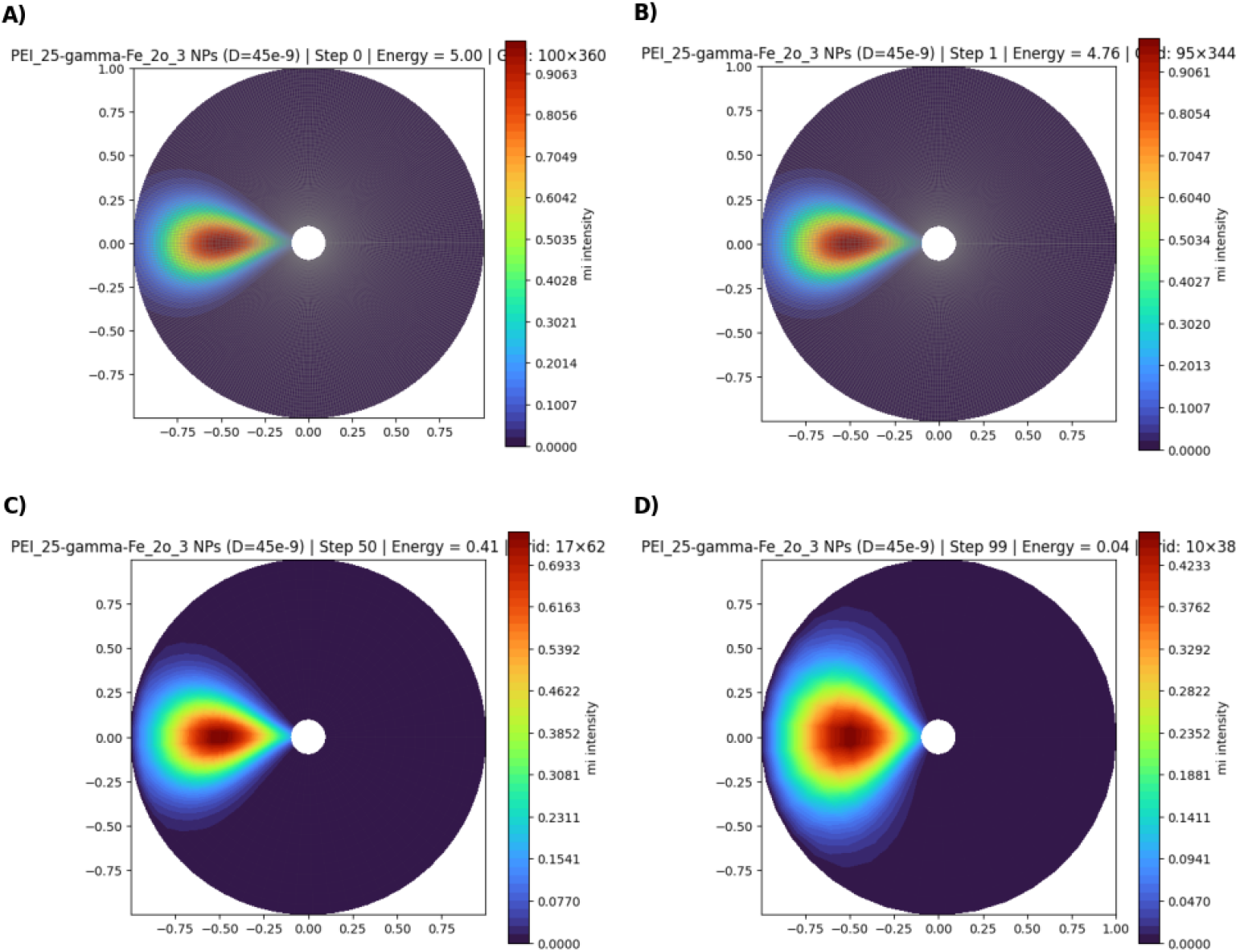
Properties Implicit in PEI_25_-Decorated *γ*-Fe_2_O_3_ Nanoparticles.

Figure 5 illustrates the normalized intensity distribution for citric acid-coated Fe_3_O_4_ nanoparticles with a mean diameter of 66 nm. The system was simulated from time step 0 to 99 at an energy level of 5.00. The polar grid spans angular divisions from 100^°^ to 360^°^. The intensity, denoted as *m*_*i*_ intensity, ranges from 0 to 0.9063, consistent with the values observed in Figure 5. The remaining subfigures A,B,C,D exhibit results qualitatively similar to those of the PEI 25-coated nanoparticle plots, showing comparable trends in angular intensity distribution and energy decay over time.

**Figure 5.**
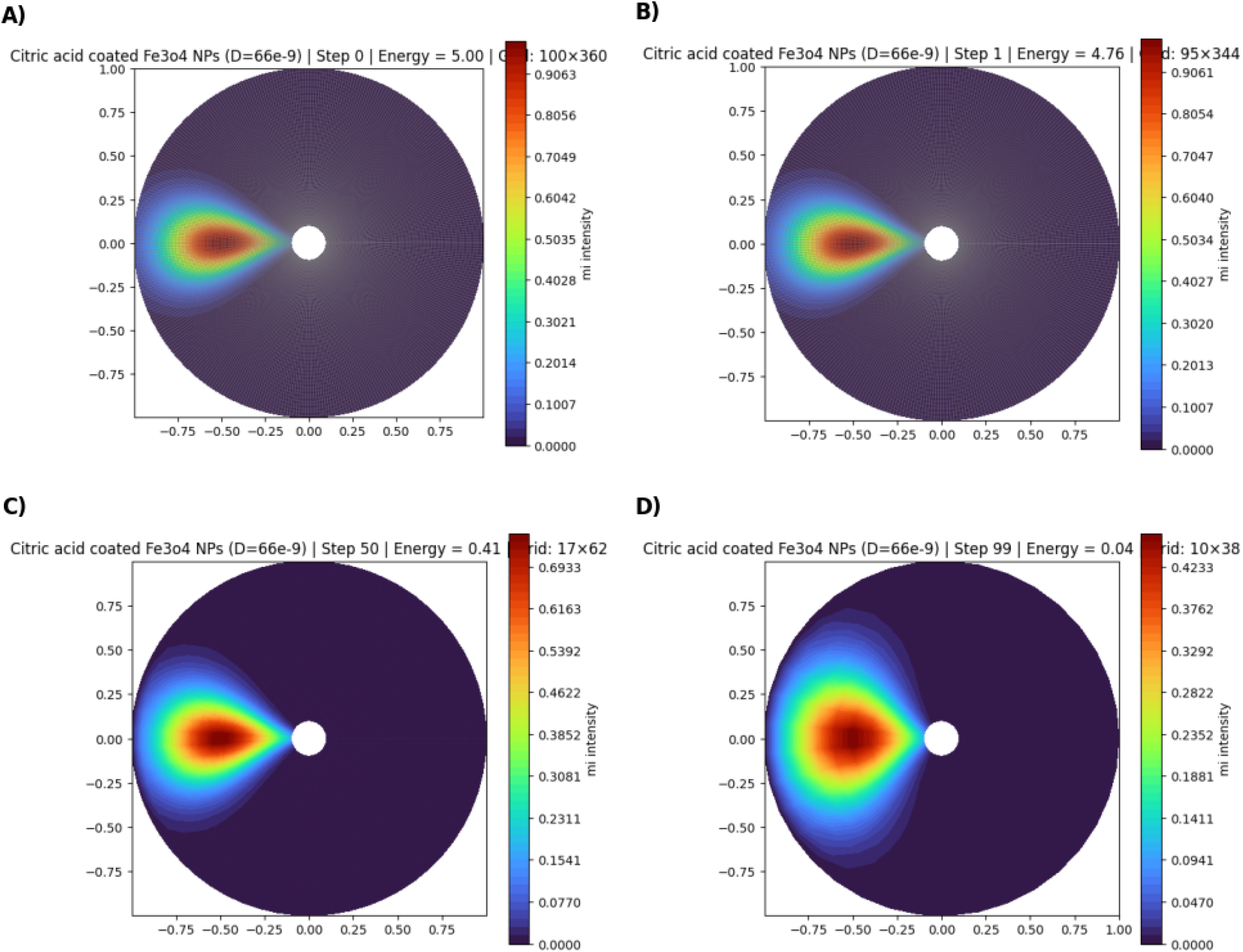
Properties Implicit in Citric Acid-Coated Fe_3_O_4_ Nanoparticles.

Figure 6 illustrates the *semi-implicit* properties of PEI_25_ *γ*-Fe_2_O_3_ nanoparticles. While Figures 4 and 5 clearly depict the *implicit* characteristics, the semi-implicit behavior shown here plays a dual role—functioning both implicitly and explicitly—which significantly enhances drug diffusion. Figure 6(A) Depicts macrophages *m*_1_ and *m*_2_ under semi-implicit conditions. The high-intensity range for *m*_1_ lies between −0.25 and 0.25. For *m*_2_, also centered around −0.25 to 0.25, the intensity begins at a medium level, with energy 4.756 and grid size 95×344. Figure 6(B) Shows *m*_1_ exhibiting high intensity between −0.50 and 1.00 at time step 4. *m*_2_ displays high intensity only along the positive axis, ranging from 0.50 to 1.00, with energy 4.094 and grid size 83 × 301. Figure 6(C) Presents a broader spread of *m*_1_ intensity, appearing medium to slightly high compared to earlier figures. Figure 6(D) Represents *m*_1_ with medium intensity, while *m*_2_ shows low intensity. However, high-intensity regions of *m*_2_ appear distributed across the entire circular domain. Figure 6(E) Demonstrates that even at step −99, *m*_2_ continues to exhibit high intensity, with energy 0.035 and grid size 10 × 38. The same Poisson pressure is applied uniformly across all steps.

**Figure 6.**
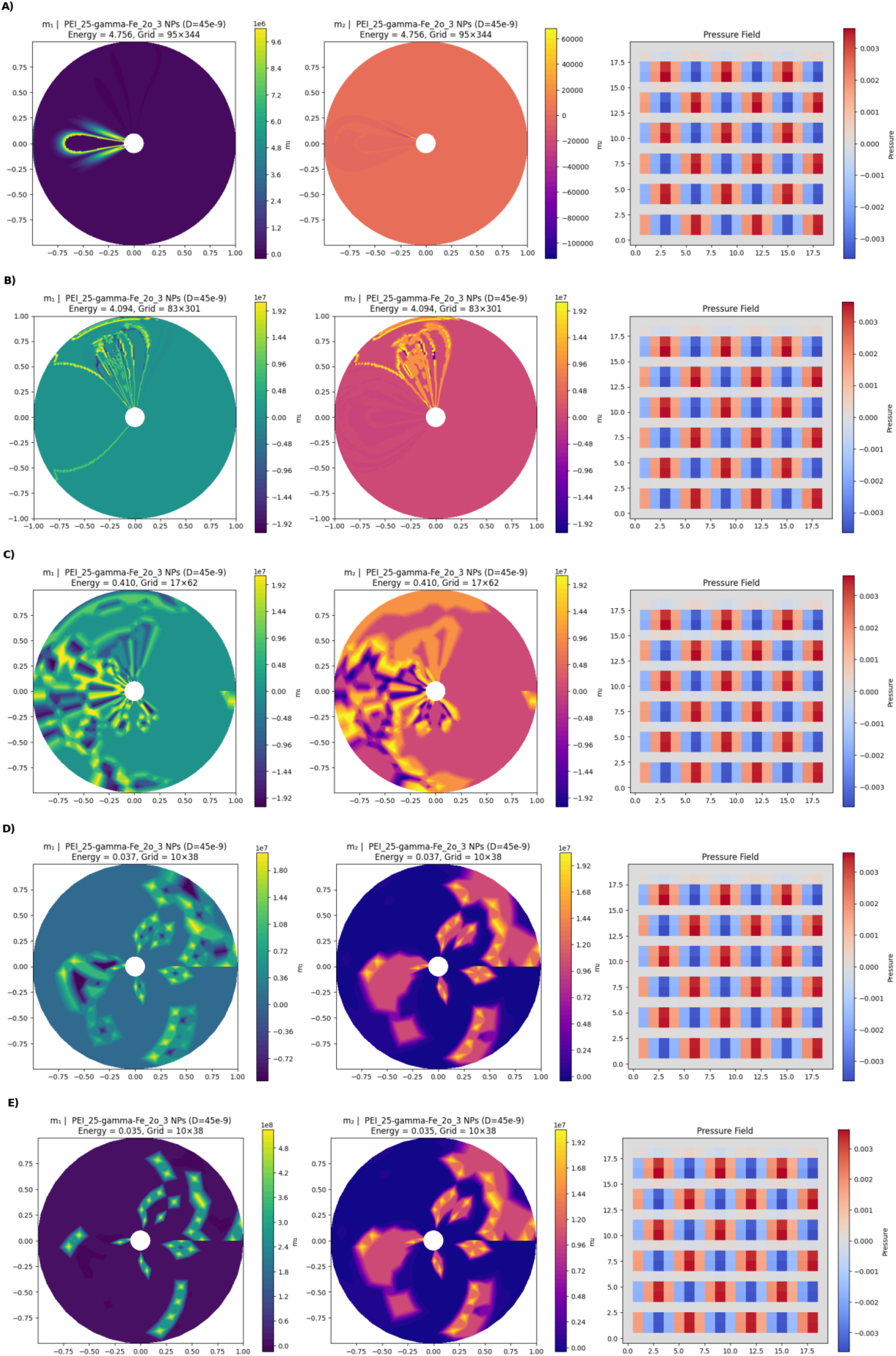
Semi-implicit Properties of PEI_25_-Decorated *γ*-Fe_2_O_3_.

Figure 8 represents the citric acid-coated iron oxide nanoparticles, beginning similarly to Figure 8. **(A)** The initial intensity range spans from −0.25 to 0.25, with energy 4.756 and grid size 95 × 344, comparable to Figure 8(B) Mirrors the spatial configuration of Figure 8(C) Shows variation: *m*_1_ begins with medium intensity at step 1, with only a few regions of high intensity. *m*_2_ displays medium intensity and slight high-intensity regions, with energy 0.410 and grid size 17 × 62. Figure 8(D) Depicts *m*_1_ transitioning to low intensity across the entire circular domain, while *m*_2_ shows medium intensity with slight high-intensity features—resembling implicit behavior. The same Poisson pressure scale is applied uniformly across all steps.

Figure 7 shows the moving mesh grid for PEI_25_-decorated *γ*-Fe_2_O_3_. Equal Poisson pressure is applied, as explained in Section 6. The figure ultimately exhibits very high intensity, with approximately half of the mesh grid showing strong activation. Figure 9 shows the semi-implicit behavior of the moving mesh grid for citric acid-coated Fe_3_O_4_ nanoparticles under Poisson pressure. The drug diffusion spreads uniformly across the mesh, with regions of high intensity appearing as localized patches.

**Figure 7.**
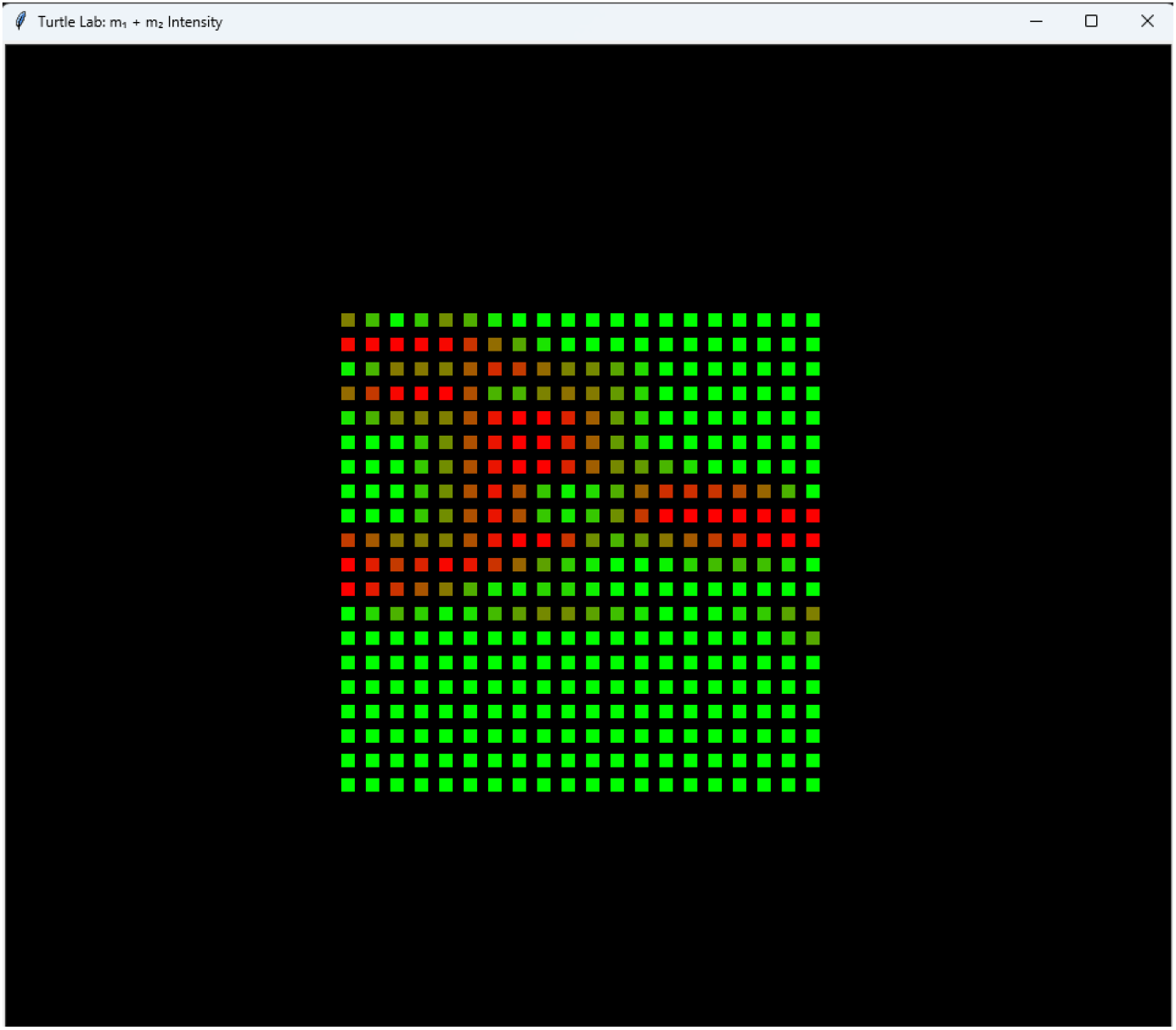
Semi-implicit of moving mesh grid for PEI_25_-Decorated *γ*-Fe_2_O_3_.

**Figure 8.**
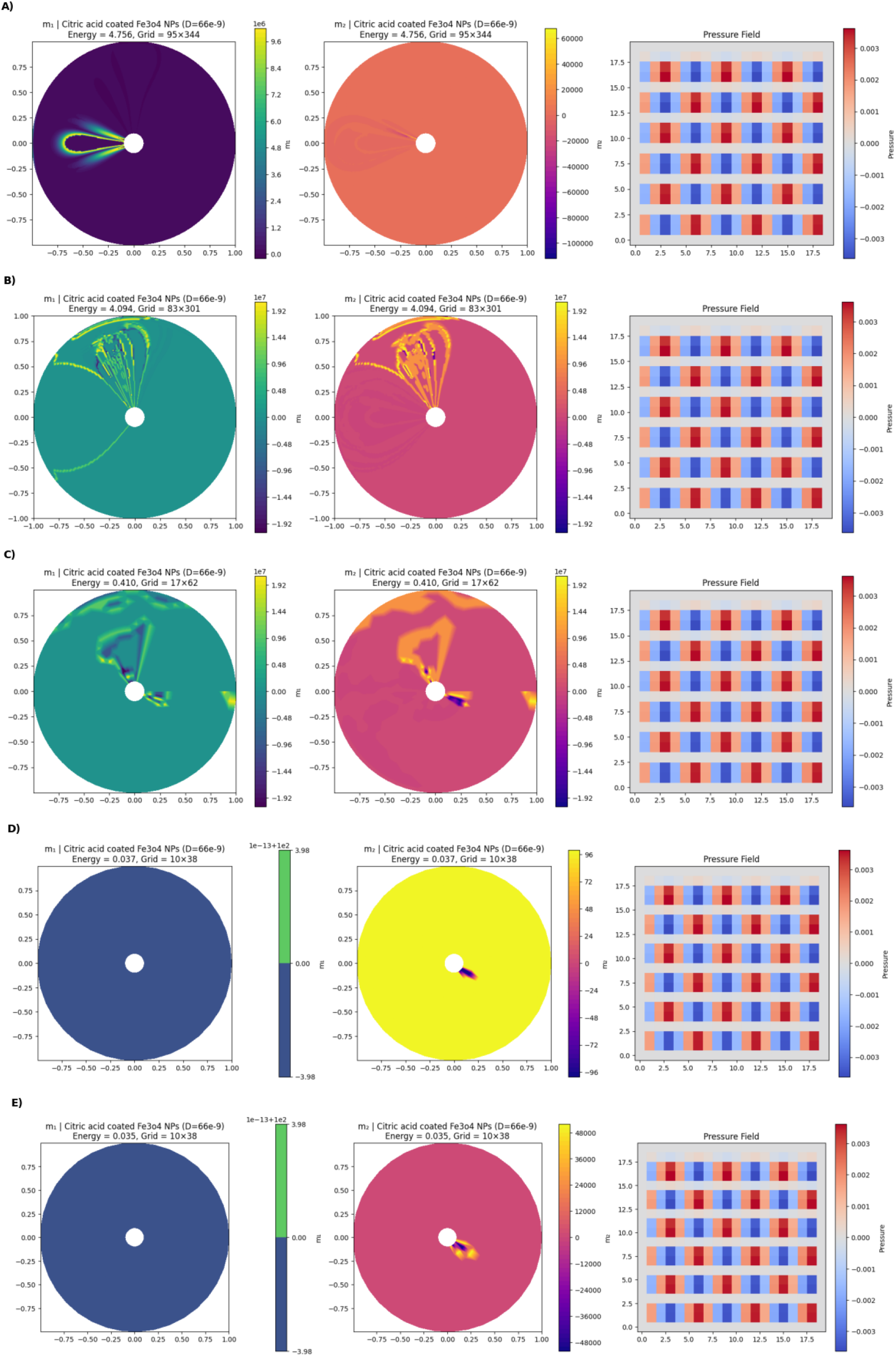
Semi-implicit Properties of Citric Acid-Coated Fe_3_O_4_ Nanoparticles.

**Figure 9.**
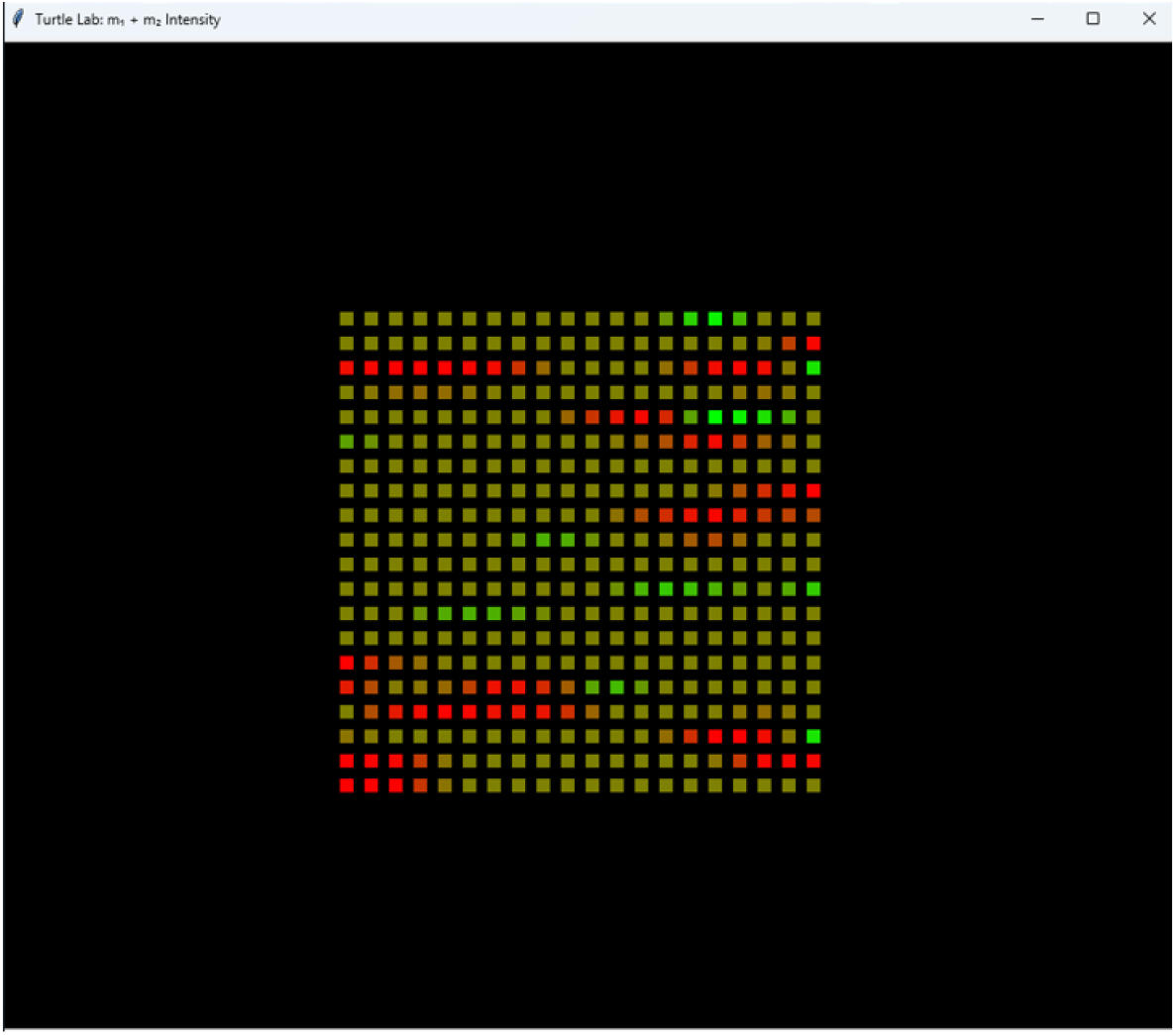
Semi-implicit of moving mesh grid for Citric Acid-Coated Fe_3_O_4_ Nanoparticles.

Figure 11 illustrates the three-phase moving boundary problem, as explained in Section 5. Based on drug diffusion dynamics, we designed the boundary evolution using three biological components: healthy macrophages, unhealthy macrophages, and nano-drug-based ligands. The model incorporates Phase-A, Phase-B, and Phase-C equations to simulate nano-drug diffusion. We tested the system with seven infected macrophages, defining the boundary range from *S*_1_(*t*) to *S*_2_(*t*). Over time, the clearance state progresses until the number of remaining infected macrophages approaches zero.

Figure 10 presents the virtual lab simulation developed based on the three-phase moving boundary problem. This simulation models drug diffusion and macrophage clearance by assigning drug concentration according to the nano-particle infection rate and the three-phase framework discussed in Section 5. In this framework:**Phase A** represents the nano-drug zone,**Phase B** is the transition zone, and **Phase C** corresponds to the infected macrophage region. Figure 11 contains nine subplots (A to I), each representing macrophage infection and clearance dynamics: **(A)** Initial boundary: *S*_1_(*t*) = 104.0, *S*_2_(*t*) = 182.0; all 7 macrophages are infected. **(B)** *S*_1_(*t*) = 112.0, *S*_2_(*t*) = 186.0; 0 cleared, 7 remaining.**(C)** *S*_1_(*t*) = 117.6, *S*_2_(*t*) = 188.8; 1 cleared, 6 remaining. **(D)** *S*_1_(*t*) = 124.8, *S*_2_(*t*) = 192.4; 2 cleared, 5 remaining. **(E)** *S*_1_(*t*) = 127.2, *S*_2_(*t*) = 193.6; 3 cleared, 4 remaining. **(F)** *S*_1_(*t*) = 124.8, *S*_2_(*t*) = 192.4; 4 cleared, 3 remaining.**(G)** *S*_1_(*t*) = 127.2, *S*_2_(*t*) = 193.6; 5 cleared, 2 remaining. **(H)** *S*_1_(*t*) = 130.4, *S*_2_(*t*) = 195.2; 6 cleared, 1 remaining.**(I)** *S*_1_(*t*) = 132.48, *S*_2_(*t*) = 196.4; 7 cleared, 0 remaining.**(J)** Final state: *S*_1_(*t*) = 145.6, *S*_2_(*t*) = 202.8; 7 cleared, 0 remaining.Our virtual lab system for simulation with the help of three phase moving boundary problem acts well for the drug designing scenario.so the final time acts as the boundary *S*_2_(*t*) =202.8.

**Figure 10.**
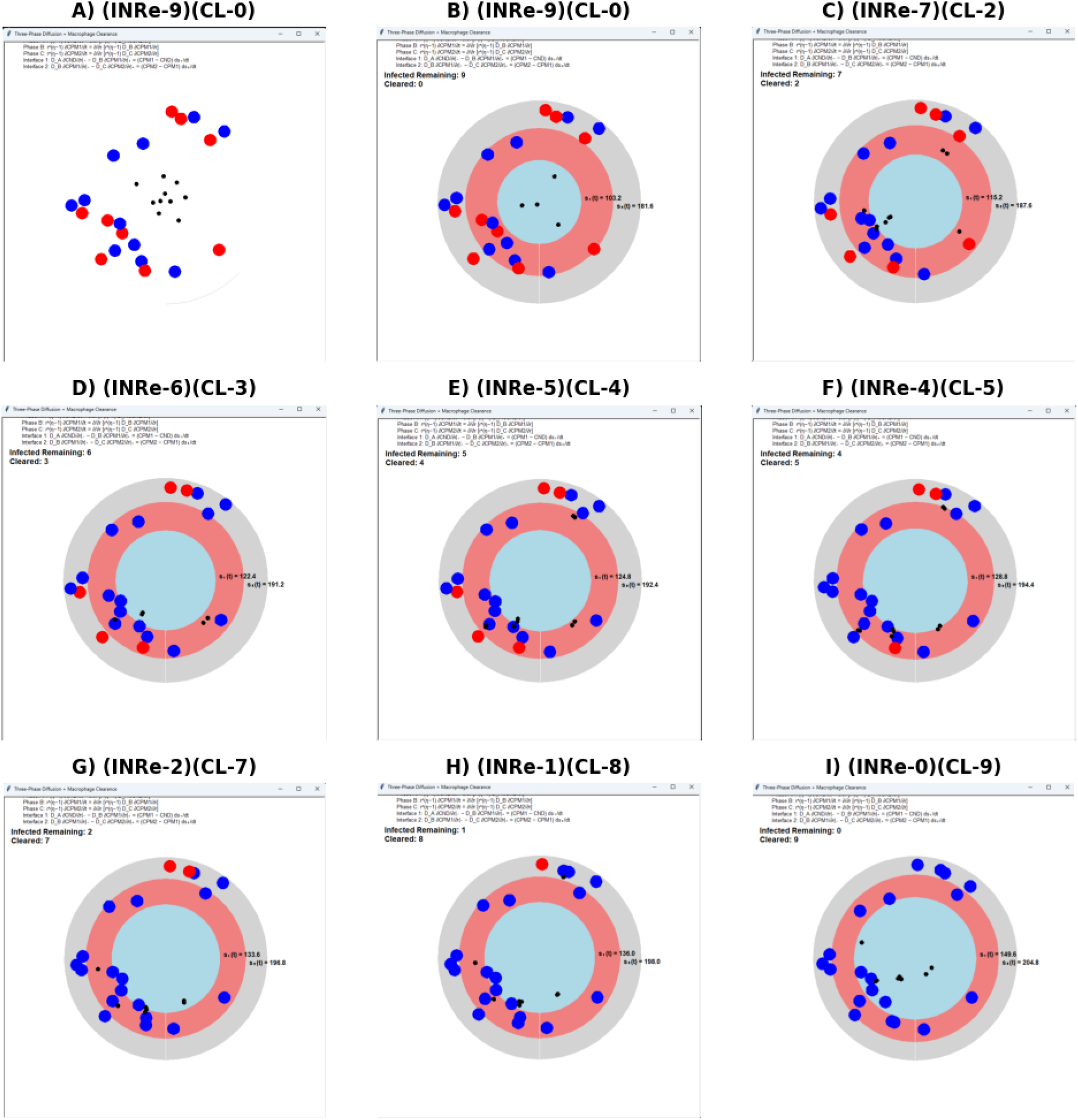
Three phase infected macrophage clearance.

**Figure 11.**
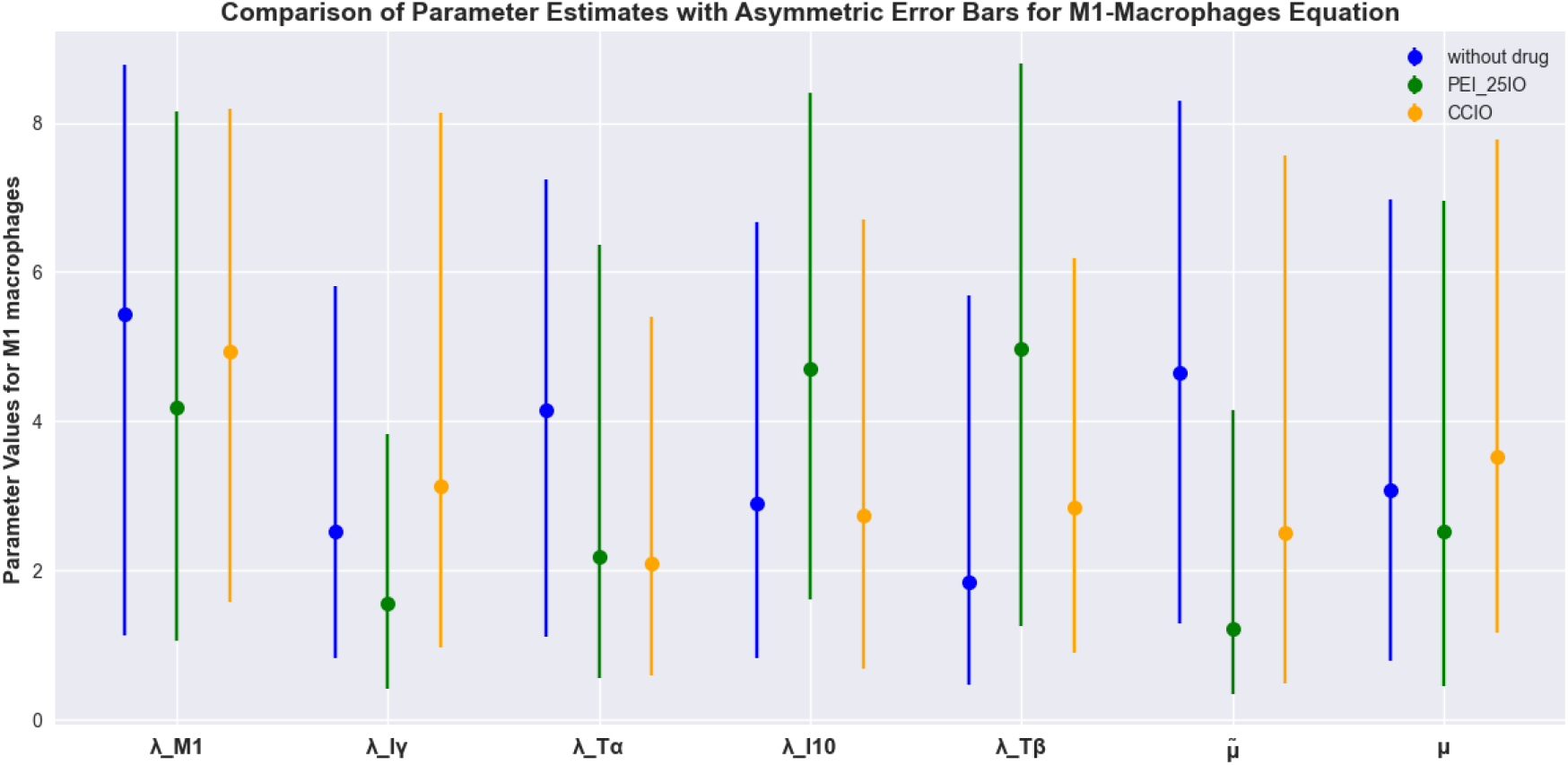
Asymmetric Error Bar for M1 macrophage.

### 7.2 Numerical results and discussion

Our main aim in this work is to study the efficacy of the drug in the transition phase, also known as the intermediate state. Our target is drug design for macrophages, parasites, and cytokines. In Section 2, we additionally derived equations for the M2 and M2a states. We introduced *M* 2*a*^∗^, which we consider as the intermediate state. These are regarded as hidden states of macrophages, so we assumed and derived equations for *M* 2*a*^∗^, *M* 2*b*^∗^, and *M* 2*c*^∗^. In parallel, parasites also play an important role in macrophage dynamics during infection. Therefore, we derived equations for *P* 2^∗^, *P* 3, *P* 4, *P* 4^∗^, and *P* 5, all belonging to the same genetic species.

The initial conditions for these derivations are discussed in the supplementary section. We investigated the uncertainty of the system using statistical methods. In the computational part, we applied Bayesian inference to estimate the posterior distribution of the parameters. Initially, we performed log-likelihood estimation, and then we applied the Monte Carlo method to quantify uncertainty [96]. Based on this, we derived asymmetric error bars corresponding to the uncertainty values of the parameters. Figures 11 shows that the uncertainty in the parameter values for M1 macrophages, specifically *λ*_*I*_, *γ*, and *µ*, is very low. This result is consistent, since in pro-inflammatory macrophages the death rate is relatively low. For M2 macrophages (anti-inflammatory), after infection the system slows down for certain parameters, as shown in Figure 12. The remaining figures are provided in the supplementary section.

**Figure 12.**
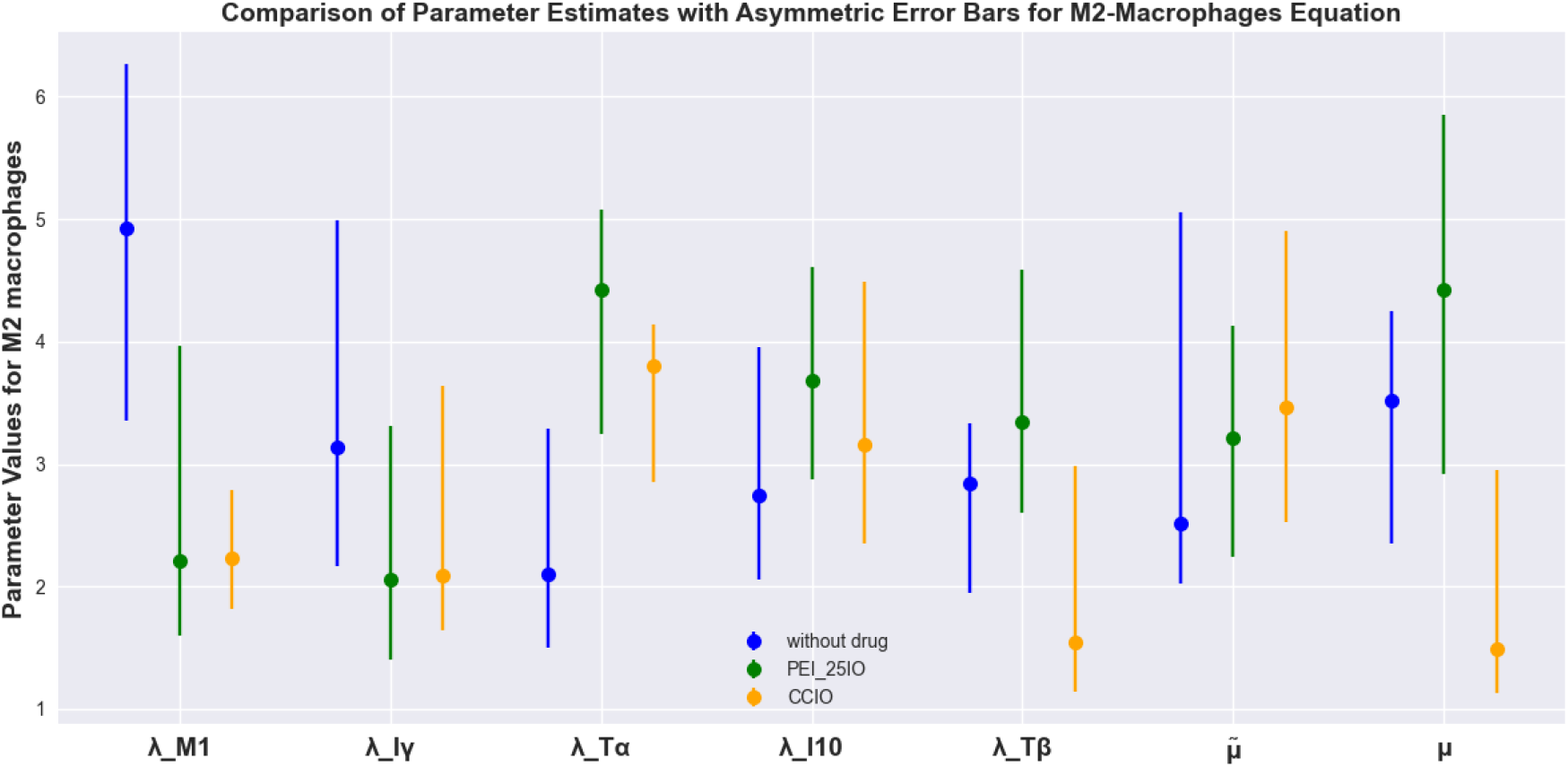
Asymmetric Error Bar for M1 macrophage.

We tested cytokine IL-4 levels with and without drug treatment using PEI25 and citric acid–coated iron oxide, based on the pulsed drug effect described in Section 2. With the CCIO drug, the cytokine level was significantly lower compared to IO. Similarly, for cytokines IL-10, IL-13, IFN-*γ*, IL-33, IL-35, NO, NO1, IL-12, IL-2, IL-6, IL-17, IC, TLR4, and PDL1, the drug effect was evident. However, for T-*α* and T-*β*, the values were nearly equal, with only slight deviations at the decimal level.

## 8 Conclusion

In this final section, we conclude based on the results obtained in the previous sections. From Section 2, we applied the parasite strains Leishmania donovani and Leishmania mexicana and tested them numerically. When combining L. mexicana drug treatment with L. donovani, our results showed better efficacy using numerical techniques such as LSODA. In Section 3, the results demonstrated that the velocity term *δ* U produced better outcomes in the presence of B cells. Section 5 highlighted that implicit and semi-implicit schemes applied to PEI25 and citric acid–coated iron oxide yielded different results. Compared to the implicit scheme, the semi-implicit method showed better efficacy in the case of citric acid–coated iron oxide. Section 6 addressed three-phase moving boundary problems. During very small boundary intervals, the drug successfully targeted infected macrophages while sparing healthy macrophages. Our model achieved this scenario effectively. Based on the two drugs introduced in Section 2, we tested both drug and no-drug conditions across cytokines, macrophages, cells, and parasites. The model consistently showed better efficacy with drug treatment, as discussed earlier. Finally, we conclude that for visceral leishmaniasis, citric acid–coated iron oxide provides better efficacy. However, the effectiveness may vary depending on the concentration of the coating. We have demonstrated this mathematically through numerical simulations and virtual laboratory environments. Nevertheless, additional clinical trials are required before the drug can be used in practice.

## Disclaimer

The work/opinion is based on research findings by the authors and not the opinion of the government.

## Data Availability

All relevant data are within the manuscript.

