## Supplemental File for "Dual Nanoparticle-Driven Therapeutics for Leishmaniasis: A Mathematical Model of Targeted Macrophage and Parasite Elimination"

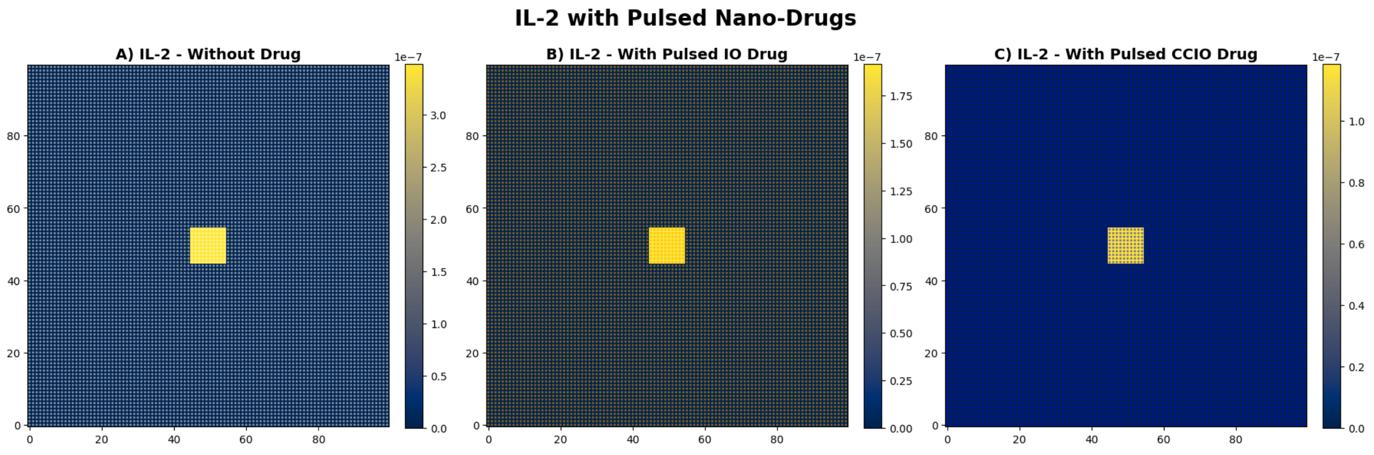


Figure S1: **Comparison of pulsed drug effect for $IL-2$.**

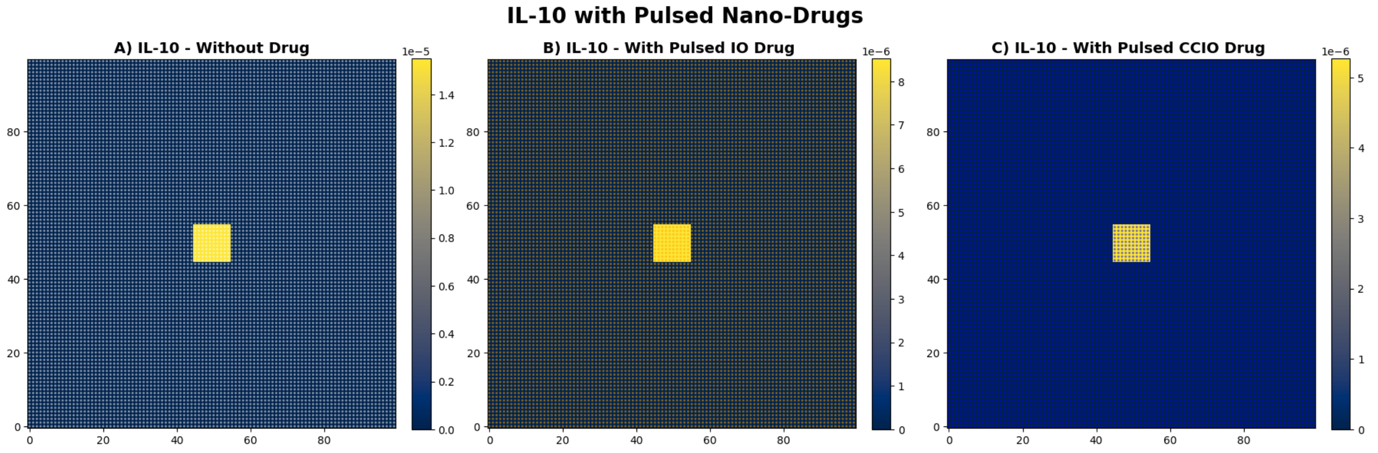


Figure S2: **Comparison of pulsed drug effect for $IL - 10$.**

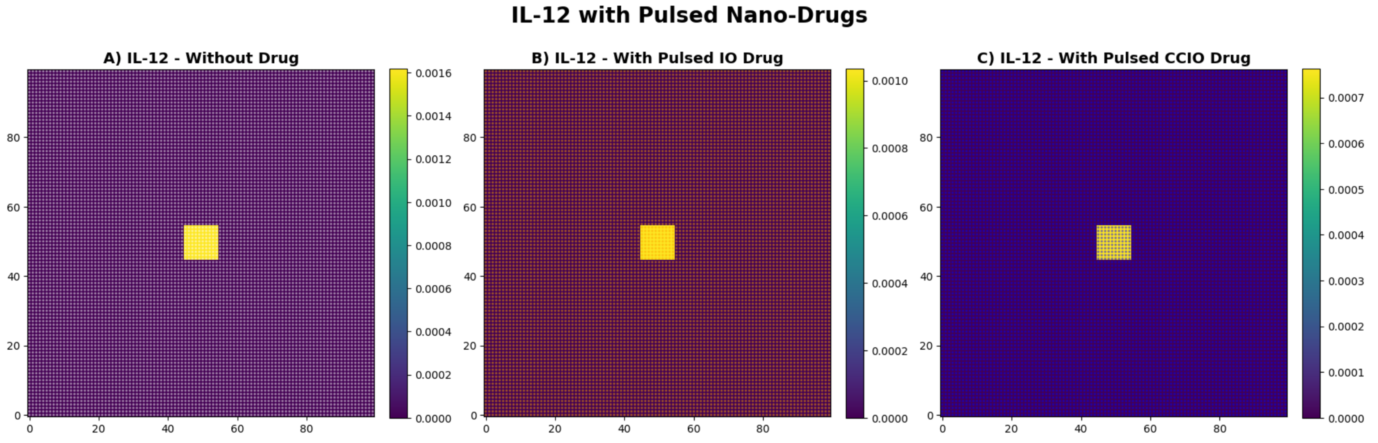


Figure S3: **Comparison of pulsed drug effect for $IL - 12$.**

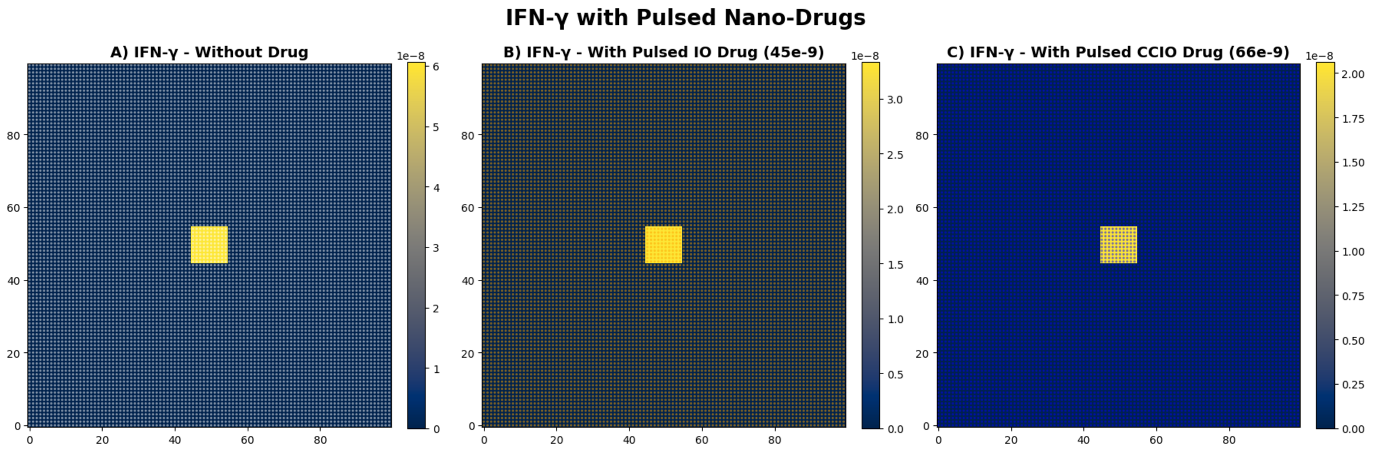


Figure S4: **Comparison of pulsed drug effect for $INF - \gamma$.**

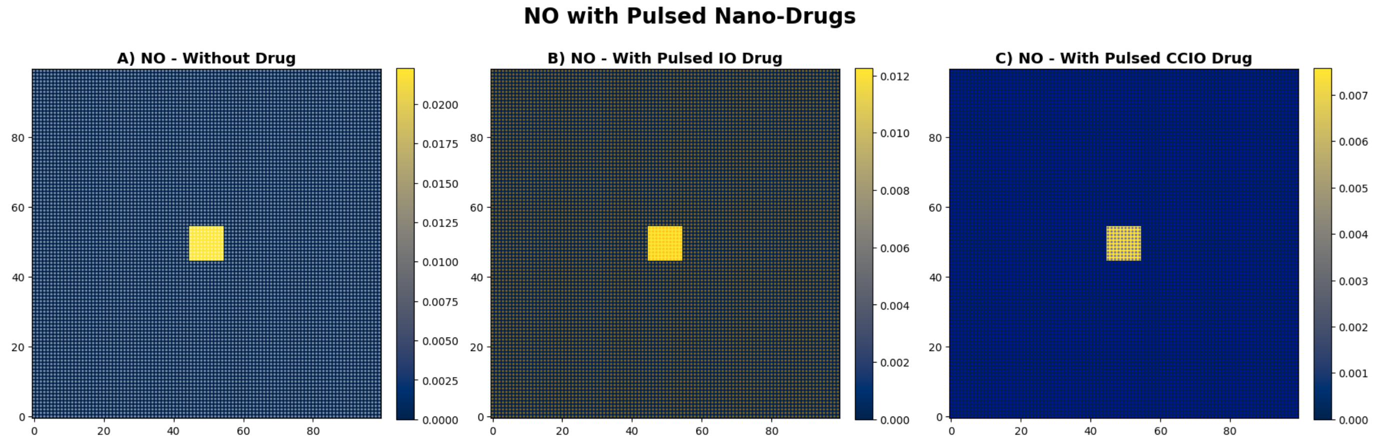


Figure S5: **Comparison of pulsed drug effect for NO.**

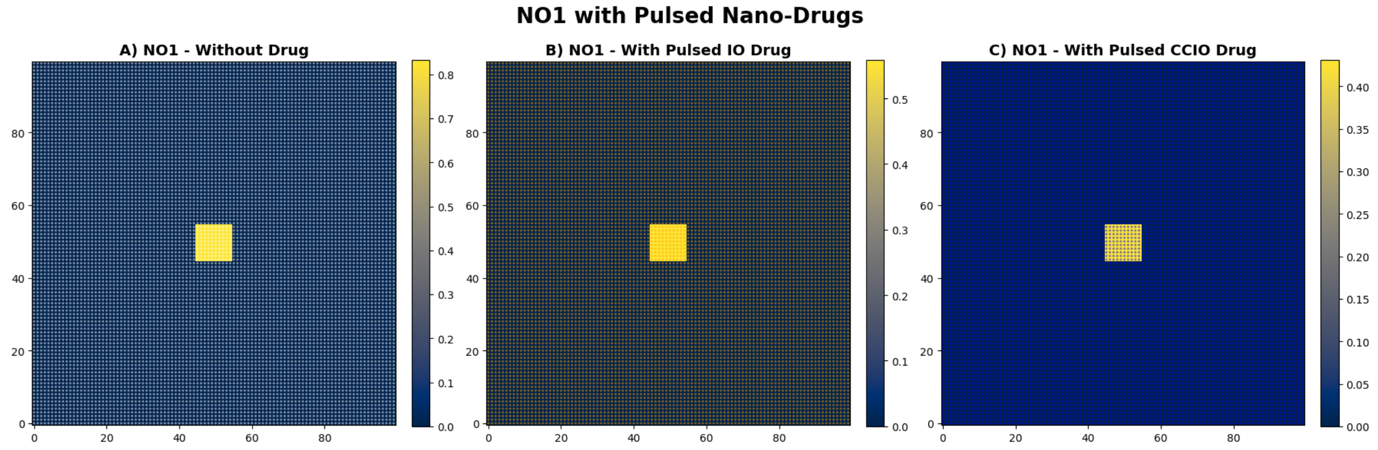


Figure S6: **Comparison of pulsed drug effect for NO1.**

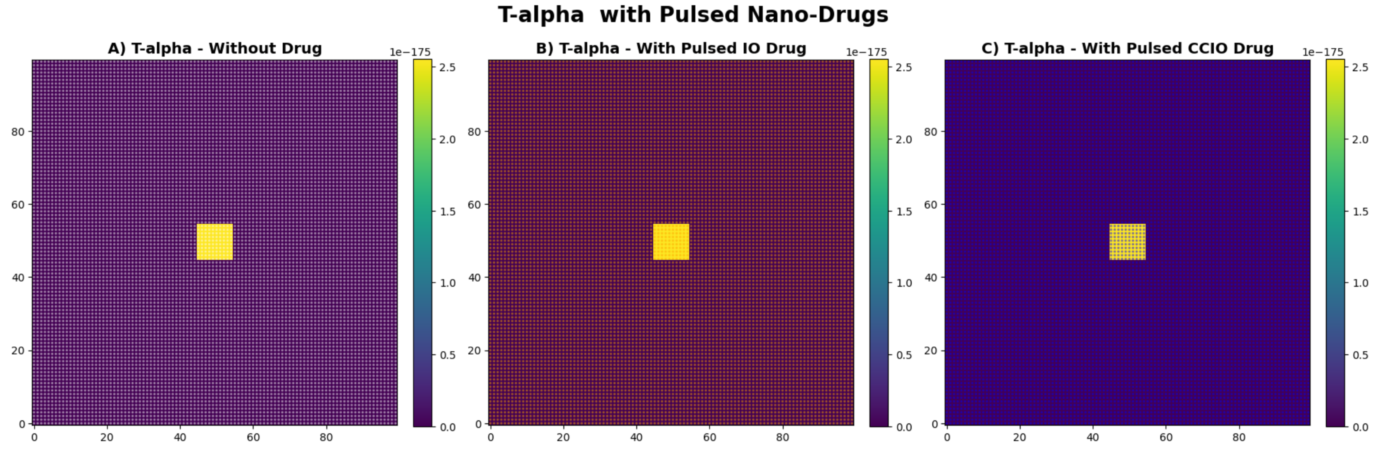


Figure S7: **Comparison of pulsed drug effect for T-alpha.**

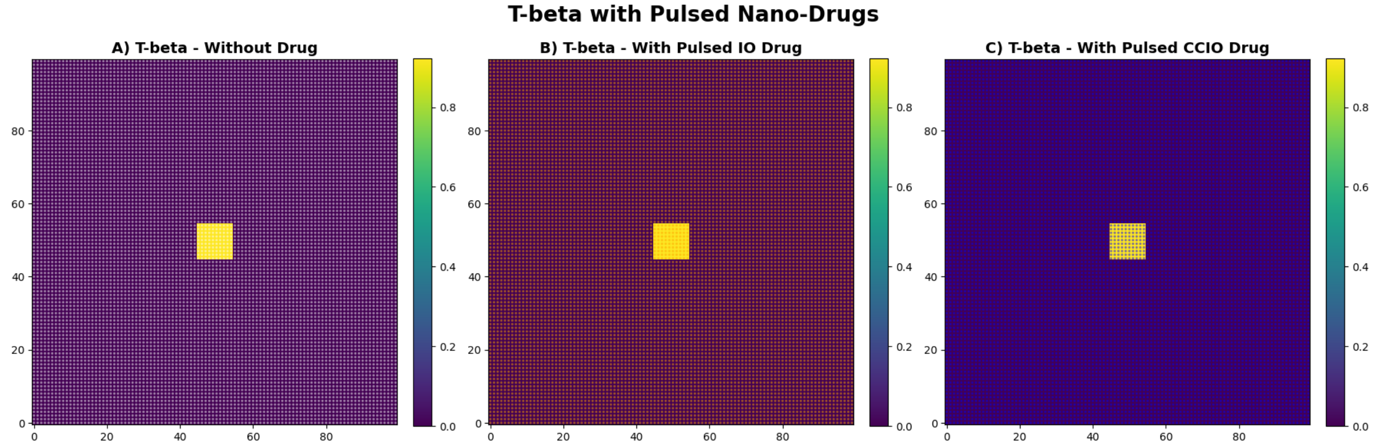


Figure S8: **Comparison of pulsed drug effect for T-beta.**

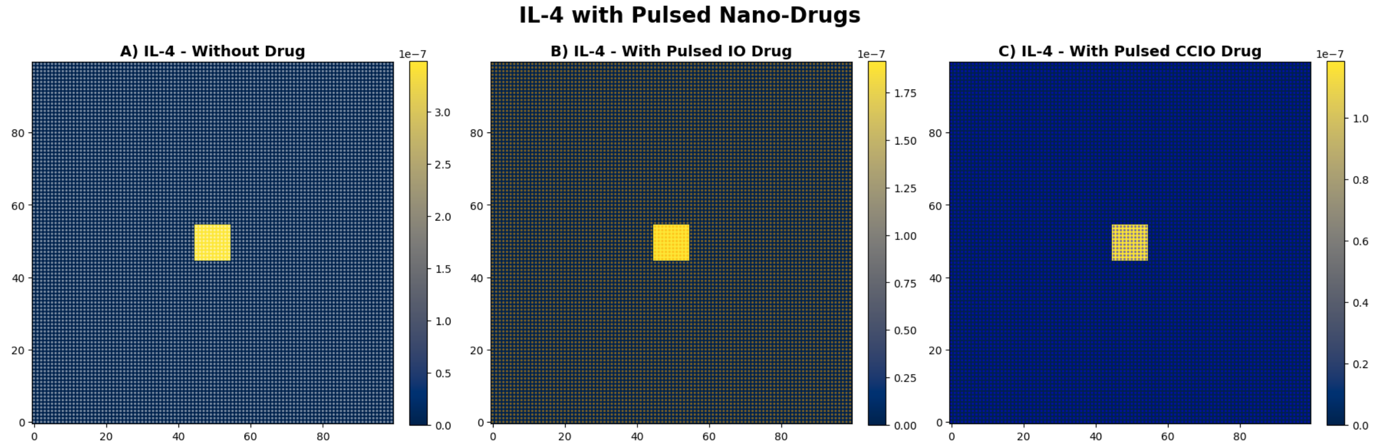


Figure S9: **Comparison of pulsed drug effect for** $IL-4$**.**

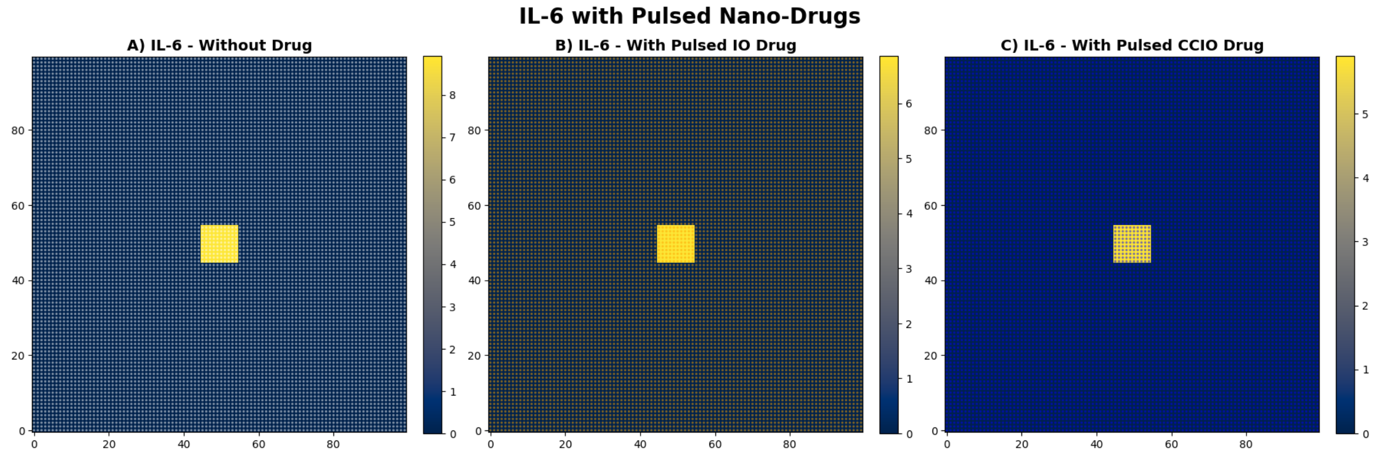


Figure S10: **Comparison of pulsed drug effect for** $IL-6$**.**

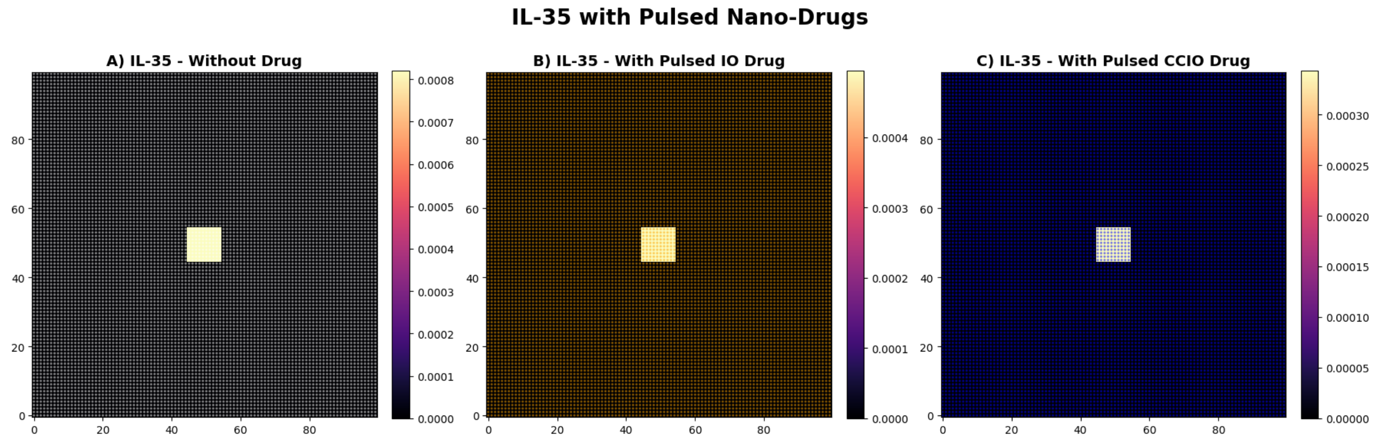


Figure S11: **Comparison of pulsed drug effect for $IL-35$.**

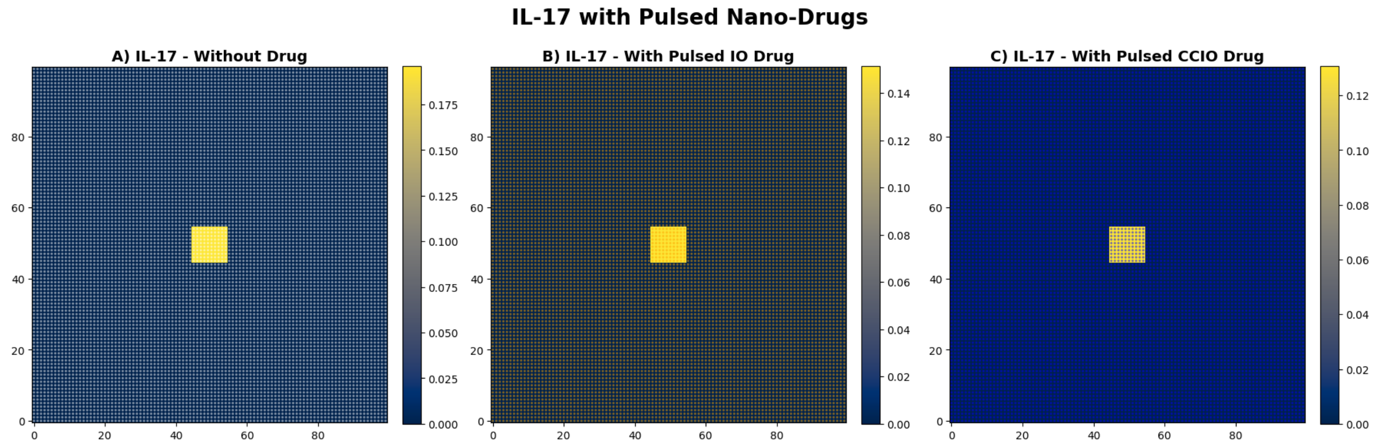


Figure S12: **Comparison of pulsed drug effect for $IL-17$.**

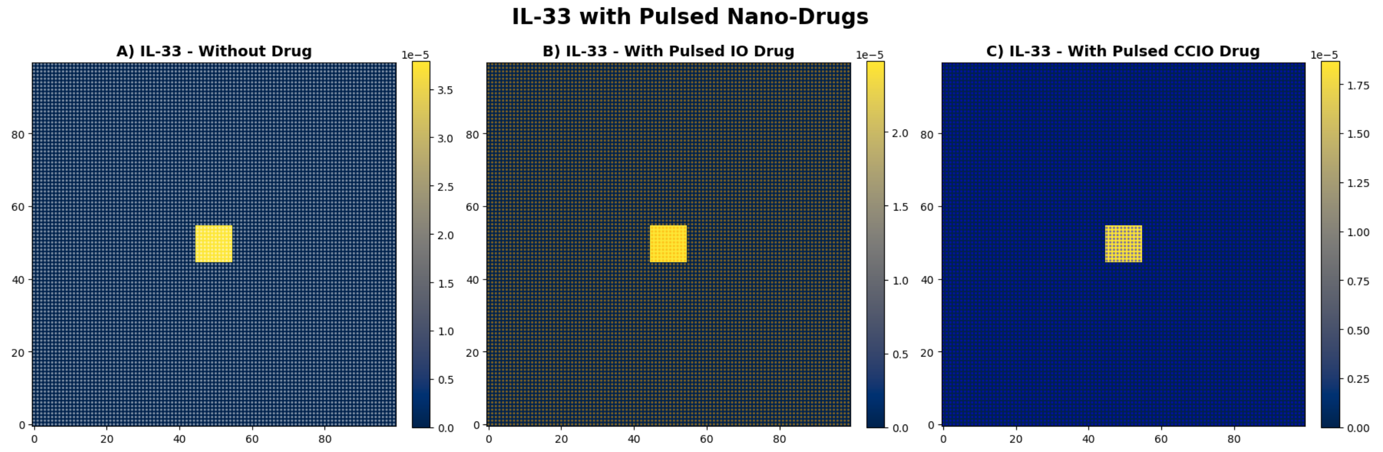


Figure S13: **Comparison of pulsed drug effect for $IL-33$.**

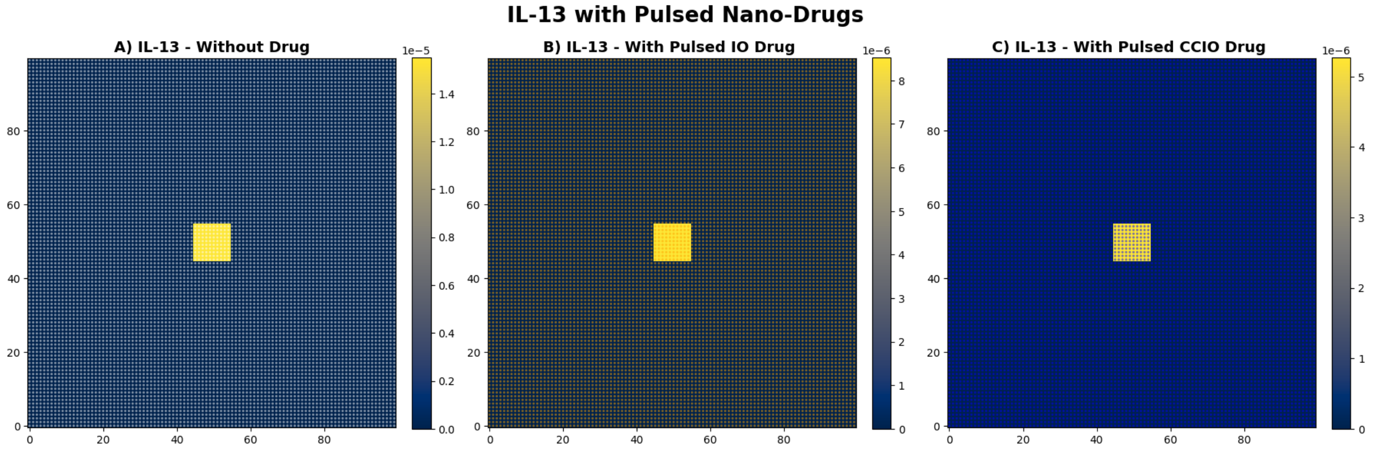


Figure S14: **Comparison of pulsed drug effect for** $IL-13$**.**

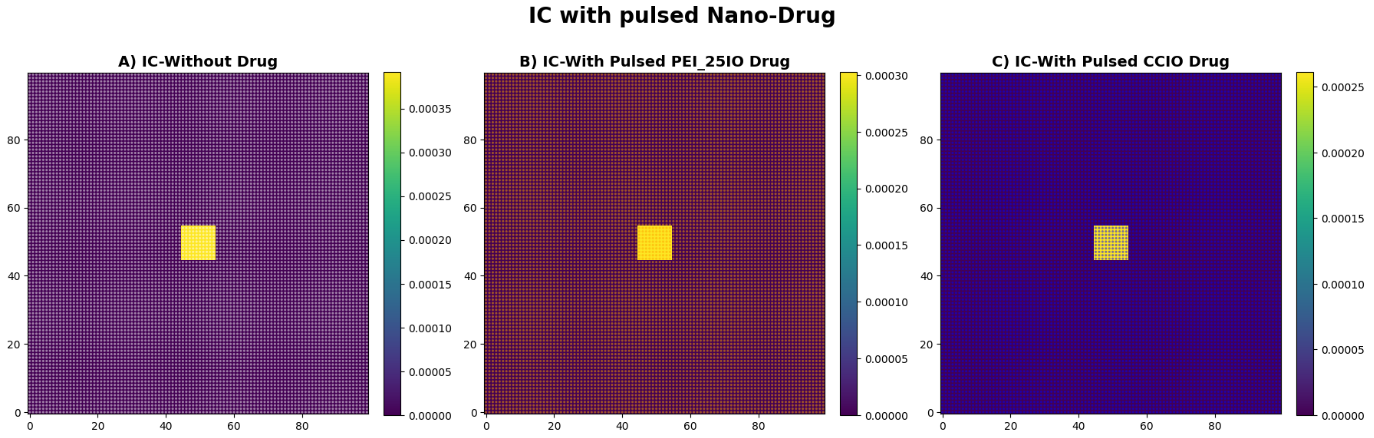


Figure S15: **Comparison of pulsed drug effect for** $IC$**.**

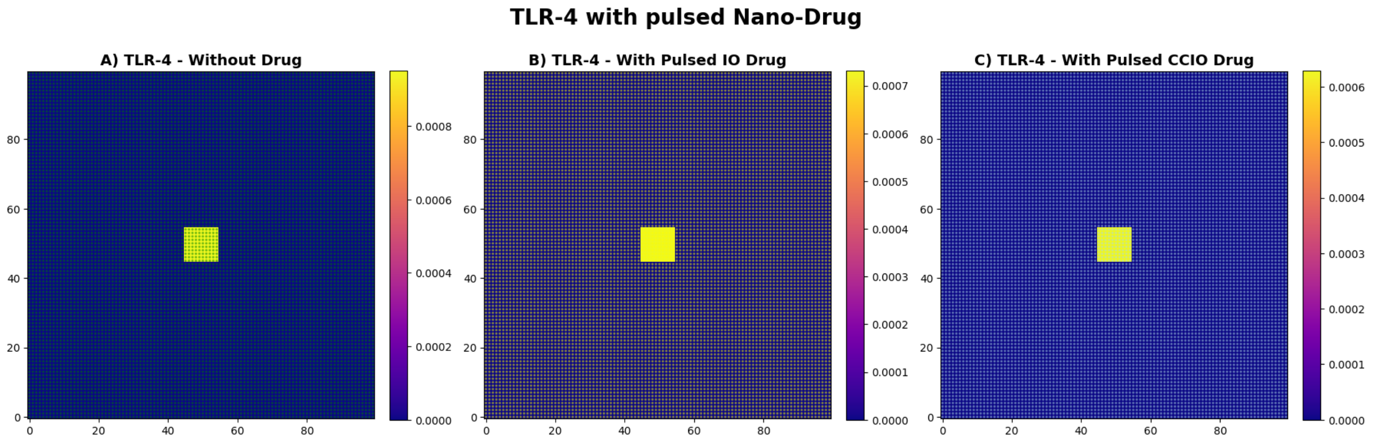


Figure S16: **Comparison of pulsed drug effect for** $TLR-4$**.**

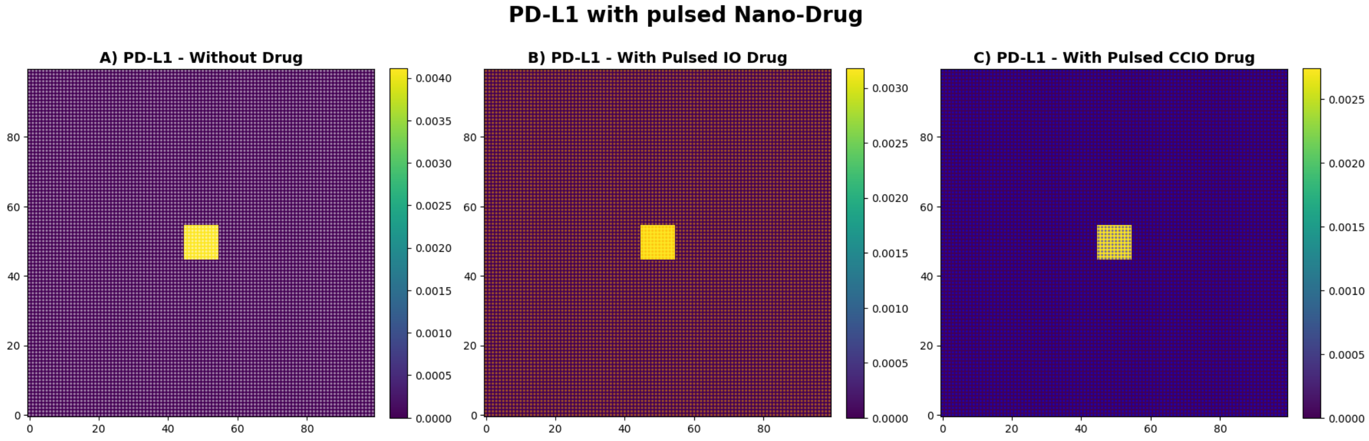


Figure S17: **Comparison of pulsed drug effect for** $PDL1$**.**

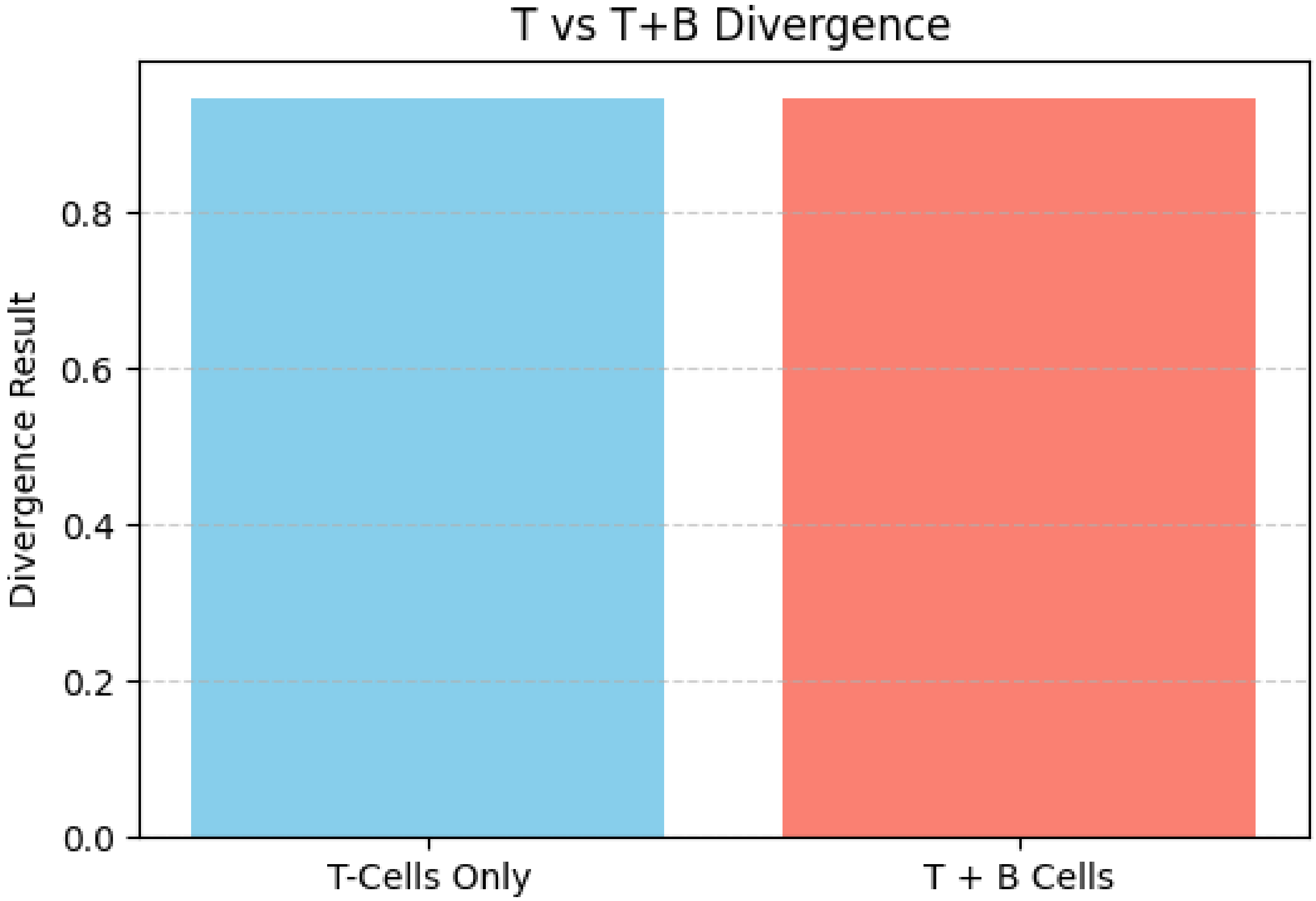


Figure S18: **Divergence of U with T-cells and T+B-cells.**

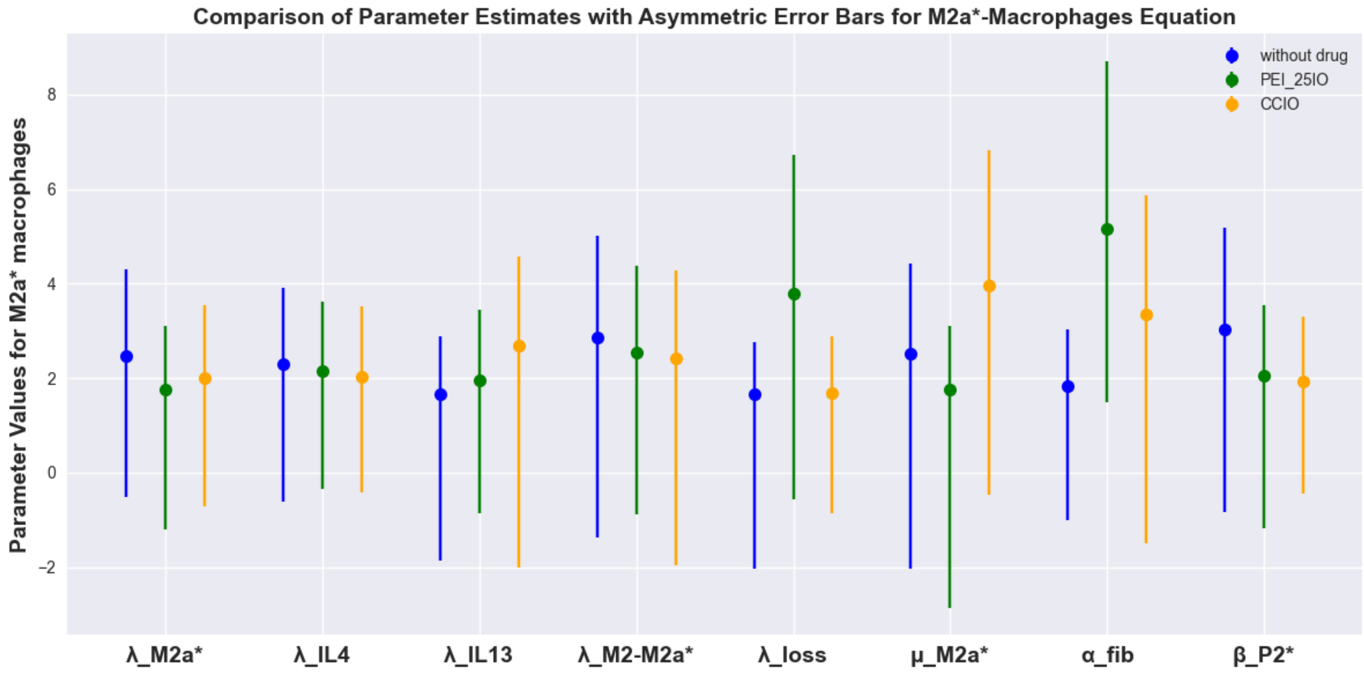


Figure S19: **Asymmetric Errors Bars for M2a*** $-$ *Macrophages*

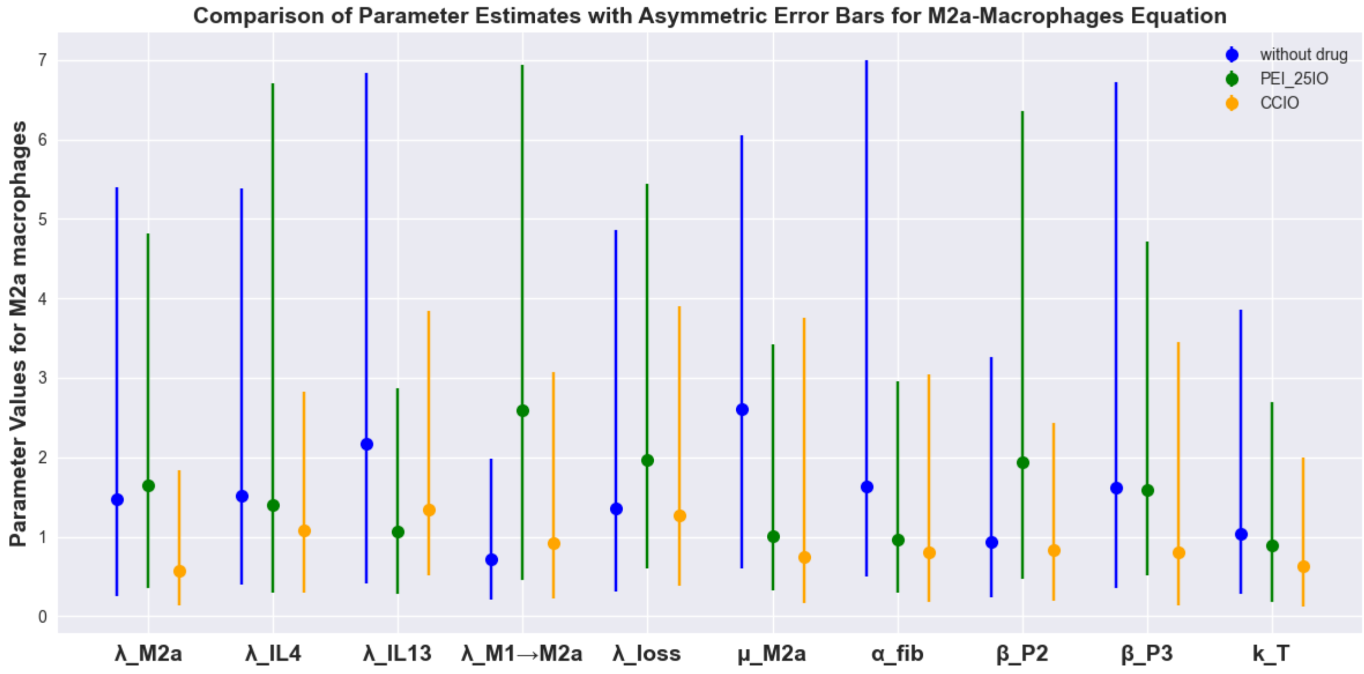


Figure S20: **Asymmetric Errors Bars for M2a-Macrophages**

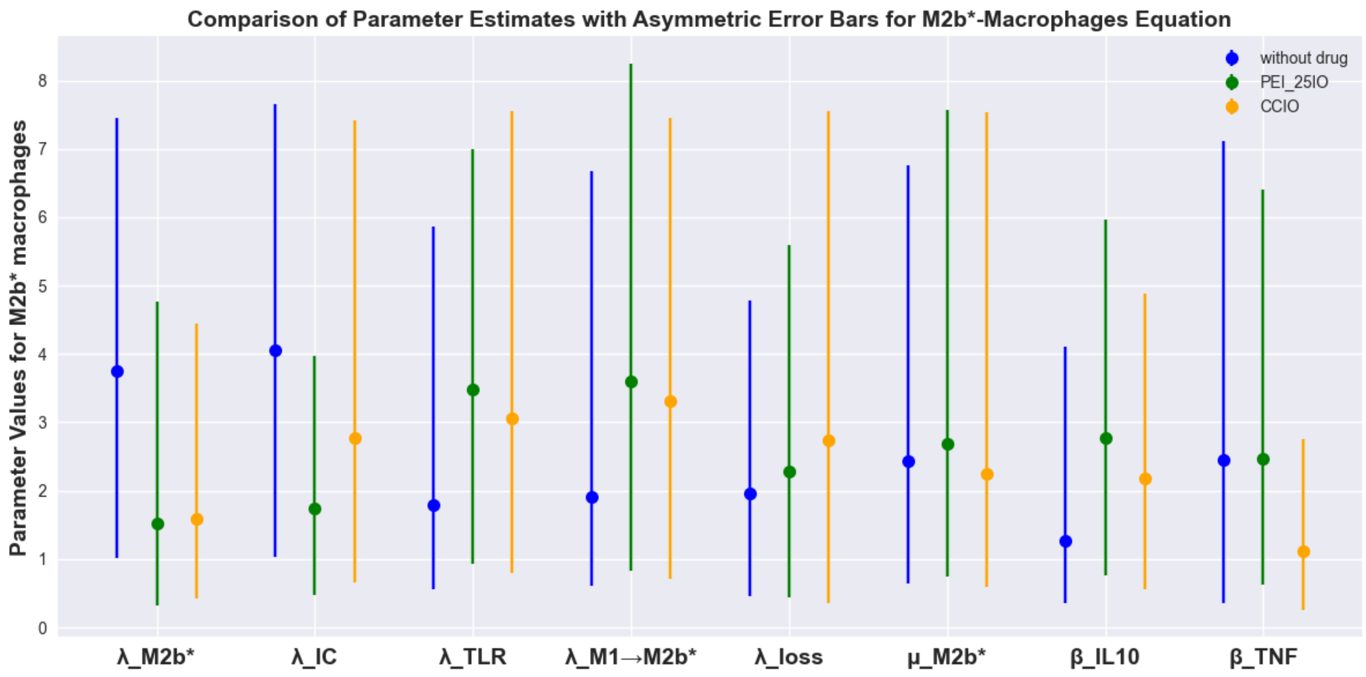


Figure S21: **Asymmetric Errors Bars for M2b*** $- \mathit{Macrophages}$

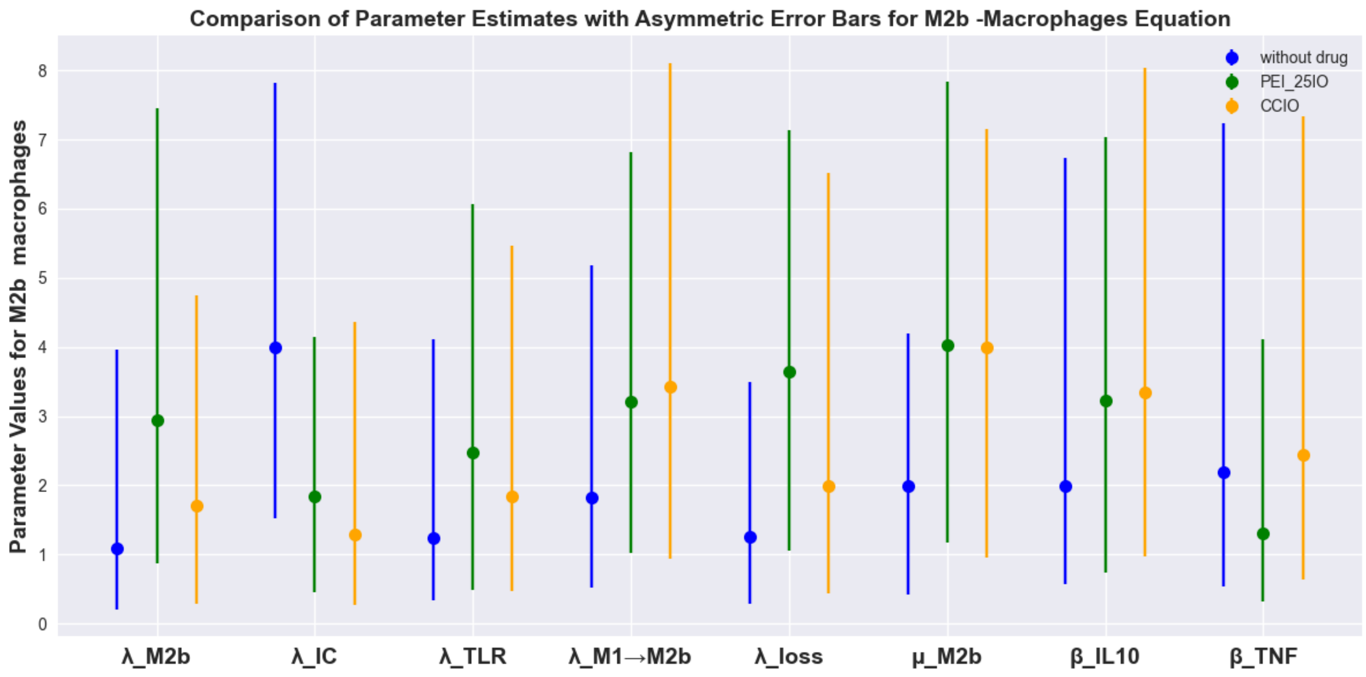


Figure S22: **Asymmetric Errors Bars for M2b-Macrophages**

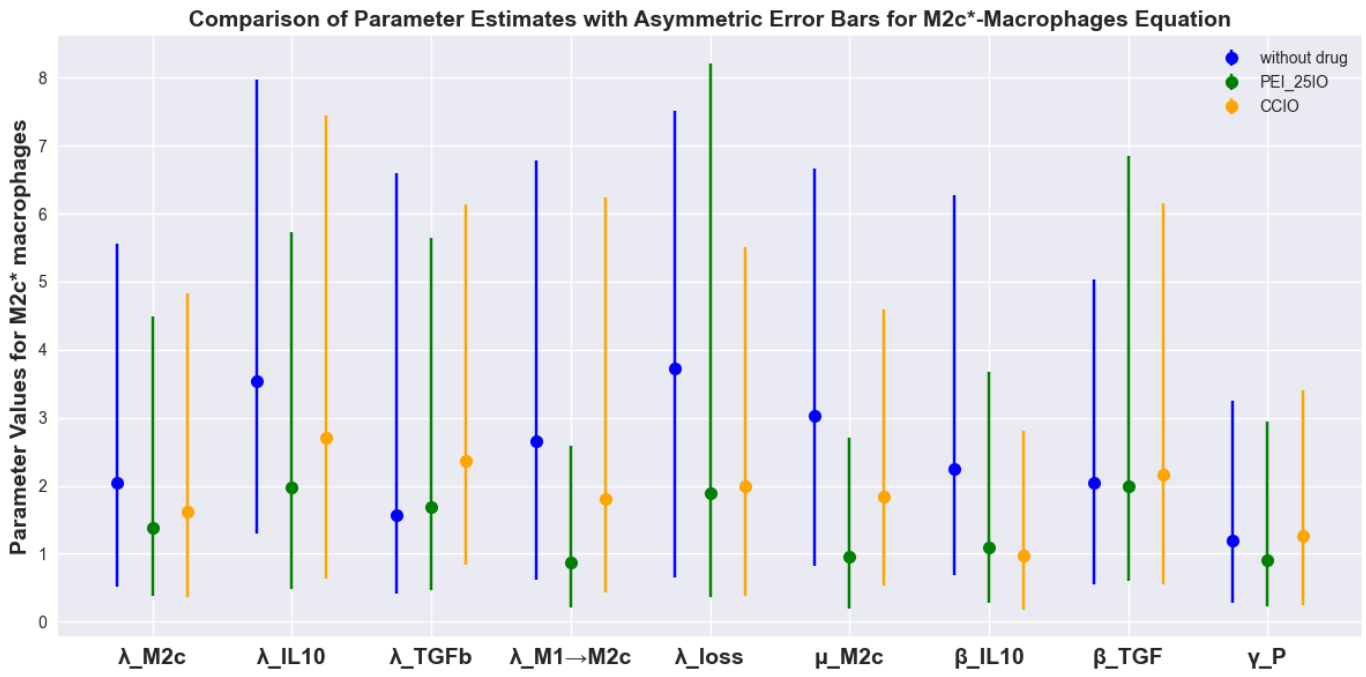


Figure S23: **Asymmetric Errors Bars for M2c*** $- Macrophages$

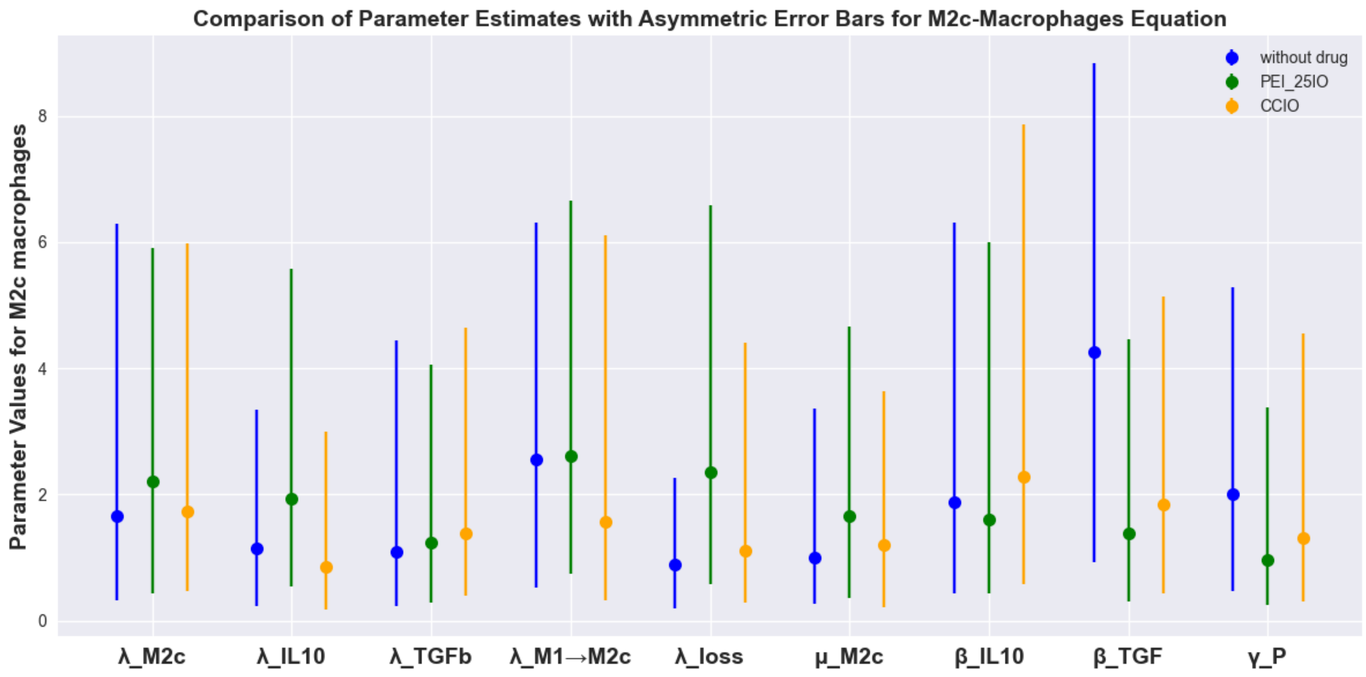


Figure S24: **Asymmetric Errors Bars for M2c-Macrophages**

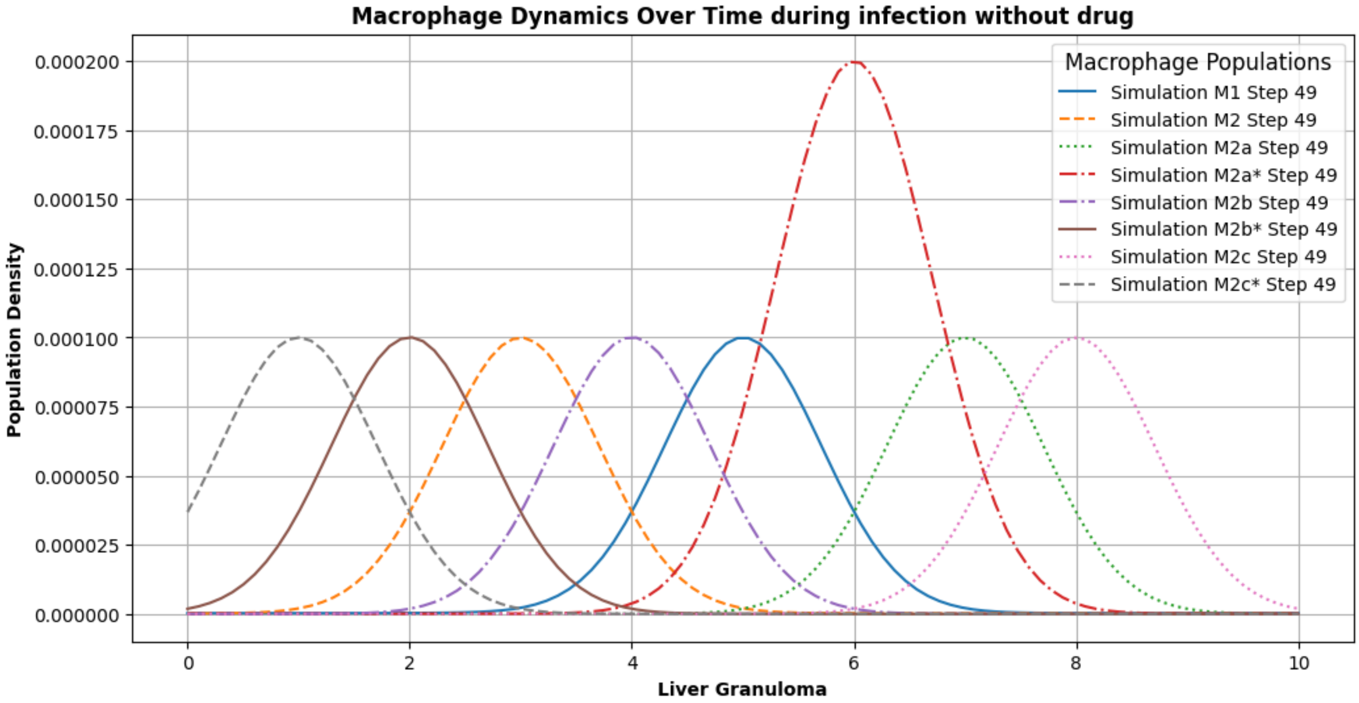


Figure S25: **Macrophage Dynamics Over Time during infection without drug**

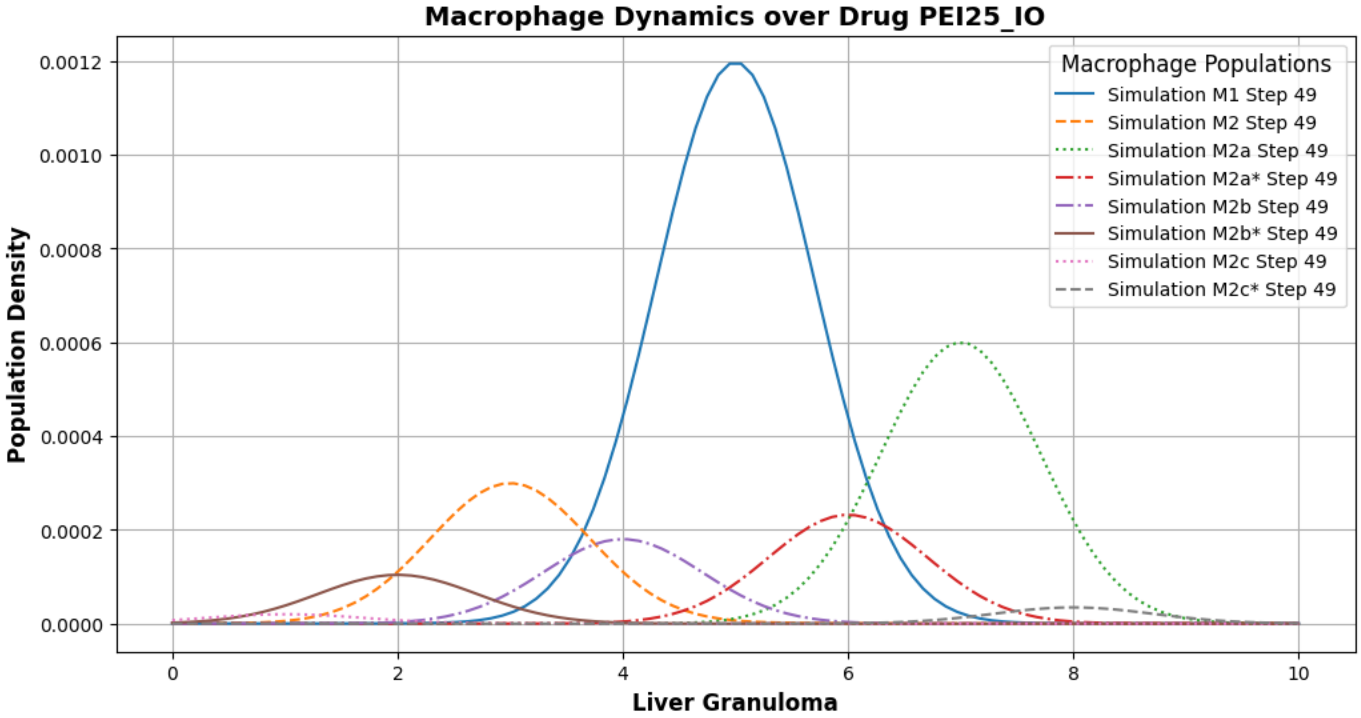


Figure S26: **Macrophage Dynamics Over drug PEI25**$_IO$

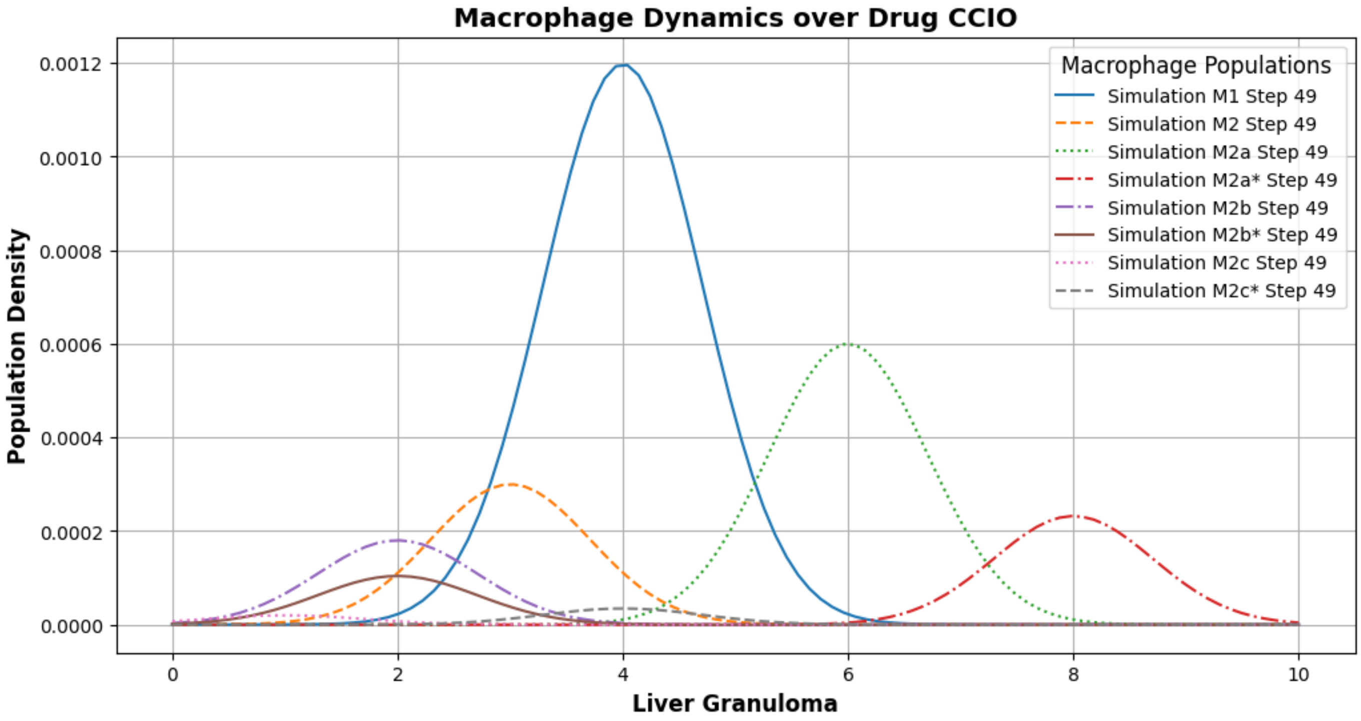


Figure S27: **Macrophage Dynamics Over drug CCIO**

Table 1: Descriptions, values, and sources of additional model parameters.

| Parameter | Description | Value/range | Reference |
|---|---|---|---|
| $k_7$ | Production rate of $I_2$ by T cells | $1.15 \times 10^{-4}$ per day | Siewe et al. (2016) |
| $k_8$ | Production rate of IL-12 by $M_1$ | $2.4 \times 10^{-3}$ per day | Siewe et al. (2016) |
| $k_9$ | Production rate of TNF-$\alpha$ by $M_1$ | $2.86 \times 10^{-3}$ per day | Siewe et al. (2016) |
| $k_{10}$ | Production rate of IL-10 and IL-13 by $M_2$ | $2.25 \times 10^{-4}$ per day | Siewe et al. (2016) |
| $k_{12}$ | Strength of IL-12 effect on activation of T cells | 2 | Siewe et al. (2016) |
| $k_4$ | Strength of IL-4 effect on activation of T cells | $1.5 \times 10^{-4}$ | Siewe et al. (2016) |
| $k_6$ | Strength of IL-6 effect on activation of Macrophage | $0.1 - 1 \times 10^{-4}$ | Siewe et al. (2016) |
| $k_6$ | Strength of IL-6 effect on activation of Macrophage | $0.1 - 1 \times 10^{-4}$ | Siewe et al. (2016) |
| $k_{35}$ | Strength of Treg effect on activation of Macrophage | $0.05 - 0.5 \times 10^{-4}$ | Siewe et al. (2016) |
| $k_{33}$ | Strength of effect on activation of epitheial | $5 \times 10^{-4}$ | Siewe et al. (2016) |
| $k_{13}$ | Strength of effect on activation of Th2 | $2.25 \times 10^{-4}$ | Siewe et al. (2016) |
| $k_{IC}$ | Strength of effect on activation of antibodies | $2 \times 10^{-4}$ | Kumagai et al.2001 |
| $k_{TLR4}$ | Strength of effect on Pamp macrophage | $2 \times 10^{-4}$ | Siewe et al. (2016) |
| $\mu_D$ | Death rate of DCs | 0.1 $d^{-1}$ | Siewe2023 |
| $\mu_{NO}$ | Decay rate of NO | 134.6 $d^{-1}$ | Siewe2023 |
| $\mu_{NO1}$ | Decay rate of NO1 | 1 $d^{-1}$ | [Est.] |
| $\mu_{M1}$ | Death rate of $M_1$ macrophages | 0.056 $d^{-1}$ | Siewe2023 |
| $\mu_{M2}$ | Death rate of $M_2$ macrophages | 0.014 $d^{-1}$ | Siewe2023 |
| $\mu_{P1}$ | Death rate of $P_1$ parasites | 1.85 $d^{-1}$ | Siewe2023 |
| $\mu_{P2}$ | Death rate of $P_2$ parasites | 2.22 $d^{-1}$ | Siewe2023 |
| $\mu_{IL2}$ | Decay rate of IL-2 | 0.33 $d^{-1}$ | Siewe2023 |
| $\mu_{IL4}$ | Decay rate of IL-4 | 330 $d^{-1}$ | Siewe2023 |
| $\mu_{IL35}$ | Decay rate of IL-35 | 2 $d^{-1}$ | Liao2014 |
| $\mu_{IL17}$ | Decay rate of IL-17 | 0.710 $d^{-1}$ | Intosalmi et al.2015 |
| $\mu_{IL33}$ | Decay rate of IL-33 | 1.2 $d^{-1}$ | Sego2020 |
| $\mu_{IL13}$ | Decay rate of IL-13 | 14.5 $d^{-1}$ | Hao2014 |
| $\mu_{IC}$ | Decay rate of IC | 0.01 $d^{-1}$ | Kumagai et al.2001 |
| $\mu_{TLR4}$ | Decay rate of TLR4 | 0.05 $d^{-1}$ | Kropf ei al.2004 |
| $\mu_{PDL1}$ | Decay rate of PDL1 | 0.02 $d^{-1}$ | FonsecaMartins et al.2019 |
| $\mu_{IL6}$ | Decay rate of IL6 | 0.551 $d^{-1}$ | Morel et al.2017 |
| $\mu_{IFN\gamma}$ | Decay rate of IFN-$\gamma$ | 14.5 $d^{-1}$ | Siewe2023 |
| $\mu_{IL12}$ | Decay rate of IL-12 | 13.9 $d^{-1}$ | Siewe2023 |
| $\mu_{TNF\alpha}$ | Decay rate of TNF-$\alpha$ | 3.4 $d^{-1}$ | Siewe2023 |
| $\mu_{TGF\beta}$ | Decay rate of TGF-$\beta$ | 399.25 $d^{-1}$ | Siewe2023 |
| $\mu_{Trans}$ | Transition rate between macrophage phenotypes | 0.5 $d^{-1}$ | [Assumed] |

Table 2: Descriptions, values, and sources of additional model parameters.

| **Parameter** | **Description** | **Value/range** | **Reference** |
|---|---|---|---|
| $\eta_{Trans}$ | Gain Transition rate between macrophage phenotypes | $0.9\ \mathrm{d}^{-1}$ | [Assumed] |
| $\lambda_{M_1}$ | Basal production rate of $M_1$ macrophages | $4.94 \times 10^{-5}\ \mathrm{g\,cm^{-3}\,d^{-1}}$ | Est. |
| $\lambda_{M_2}$ | Basal production rate of $M_2$ macrophages | $2.625 \times 10^{-5}\ \mathrm{g\,cm^{-3}\,d^{-1}}$ | Est. |
| $\lambda_{M_{2a}}$ | Basal production rate of $M_{2a}$ macrophages | $1.8375 \times 10^{-5}\ \mathrm{g\,cm^{-3}\,d^{-1}}$ | Est. |
| $\lambda_{M_{2b}}$ | Basal production rate of $M_{2b}$ macrophages | $5.25 \times 10^{-6}\ \mathrm{g\,cm^{-3}\,d^{-1}}$ | Est. |
| $\lambda_{M_{2c}}$ | Basal production rate of $M_{2c}$ macrophages | $2.625 \times 10^{-6}\ \mathrm{g\,cm^{-3}\,d^{-1}}$ | Est. |
| $\lambda_{P_1}$ | Intrinsic growth rate of parasites within $M_1$ | $2.21\ \mathrm{d}^{-1}$ | Est. |
| $\lambda_{P_2}$ | Intrinsic growth rate of parasites within $M_2$ | $3.82\ \mathrm{d}^{-1}$ | Est. |
| $\lambda_{P_2^*}$ | Intrinsic growth rate of parasites within $M_{2a}$ | $3.42\ \mathrm{d}^{-1}$ | Est. |
| $\lambda_{P_3}$ | Intrinsic growth rate of parasites within $M_{2b}$ | $4.06\ \mathrm{d}^{-1}$ | Est. |
| $\lambda_{P_3^*}$ | Intrinsic growth rate of parasites within $M_{2c}$ | $4.03\ \mathrm{d}^{-1}$ | Est. |
| $\lambda_{P_4}$ | Intrinsic growth rate of parasites within $M_{2b}$ | $3.50\ \mathrm{d}^{-1}$ | Est. |
| $\lambda_{P_4^*}$ | Intrinsic growth rate of parasites within $M_{2c}$ | $3.40\ \mathrm{d}^{-1}$ | Est. |
| $\lambda_{P_5}$ | Intrinsic growth rate of parasites within $M_{2c}$ | $3.90\ \mathrm{d}^{-1}$ | Est. |
| $N_b$ | Average number of parasites for macrophage burst | 30 (5–50) | Siewe2023 |
| $N_1$ | Average number of parasites in $M_1$ macrophages | 2 | Siewe2023 |
| $N_2$ | Average number of parasites in $M_2$ macrophages | 20 | Siewe2023 |
| $N_{2a}$ | Average number of parasites in $M_{2a}$ macrophages | 14 | Est. |
| $N_{2b}$ | Average number of parasites in $M_{2b}$ macrophages | 4 | Est. |
| $N_{2c}$ | Average number of parasites in $M_{2c}$ macrophages | 2 | Est. |
